# Historical contingency shapes zebrafish host-microbiome responses to a subsequent biotic challenge

**DOI:** 10.64898/2026.07.05.734762

**Authors:** Michael J. Sieler, Connor Leong, Kristin D. Kasschau, Michael L. Kent, Thomas J. Sharpton

## Abstract

Environmental change exposes ecosystems, including host-associated microbiomes, to stressors that occur repeatedly and in sequence, yet it remains unclear whether prior stressor history conditions host-microbiome responses to later perturbation. We used adult zebrafish (*Danio rerio*) to test whether sequential exposure to antibiotics, heat stress, the intestinal nematode *Pseudocapillaria tomentosa*, or pairwise stressor combinations altered gut microbiome structure, intestinal host gene expression, and host health outcomes. Across eight exposure regimes, prior stressor history and parasite exposure were associated with gut microbiome composition, while increasing prior stressor history was associated with reduced gut microbial diversity and convergence in community composition. Host intestinal transcriptional responses to parasite exposure were historically contingent, with parasite-associated differential gene expression varying non-linearly across prior stressor histories. Cumulative mortality increased with prior stressor history, whereas infection prevalence among surviving hosts decreased. Integrating microbial abundance, host gene expression, mortality, and neutral-community modeling identified *Cetobacterium*, *Culicoidibacter*, *Flavobacterium*, and *Shewanella* as candidate host-linked taxa associated with host response and survival. Collectively, these findings indicate that prior environmental stressor history shapes vertebrate host-microbiome responses to future perturbation and highlight specific gut microbial members as potential biomarkers or functional targets for follow-up studies.

## Introduction

The Anthropocene is accelerating environmental change, exposing ecosystems and host-associated communities to stressors that increasingly co-occur and recur [1–3]. Because stressors can interact synergistically, antagonistically, or additively, multi-stressor responses are often difficult to predict from single-stressor studies, and empirical syntheses show that stressor suites frequently depart from additivity [4–6]. Climate-linked drivers are expected to increase the repetition and sequencing of perturbations, including in host-microbiome systems [7, 8]. Yet most multi-stressor work emphasizes concurrent exposures, leaving the sequence and accumulation of prior perturbations, and how they shape host-associated microbiomes and host responses to a later biotic challenge, largely untested [9]. Resolving this is central to predicting the resilience of host-associated microbial ecosystems to anthropogenic stress [10].

Ecological theory offers a framework. Historical contingency predicts that the order and timing of past events shape community structure and function [11, 12], and priority effects can amplify these legacies when early perturbations alter the establishment of later organisms through niche preemption or modification [13, 14], steering communities toward divergent ecological states [15]. Host-associated microbiomes are a tractable system for testing how assembly history shapes community and host responses under environmental change [16, 17], and across hosts both stressor combination and cumulative stress history can shape host fitness and microbiome-associated phenotypes [18, 19]. Building on this theory and evidence, we hypothesize that historical stressors act as assembly filters that modify host-microbiome structure and function and thereby condition responses to future disturbance.

We tested this in the zebrafish (*Danio rerio*), a vertebrate model well suited to controlled, longitudinal, multifactorial designs that manipulate stressor order and whose gut microbiota are experimentally accessible and environmentally responsive [20, 21]. Prior work links priority effects and stressor-driven filtering to microbiome assembly and host-relevant outcomes in this system [22], and work from our group shows that abiotic and biotic factors, including warmer rearing temperatures and intestinal nematode exposure, shift gut microbiota composition and assembly and associate with divergent host phenotypes [23–25]. This foundation enables a direct test of whether successive stressor exposure conditions the host-microbiome response to a future perturbation.

Here, we asked whether successive stressor history conditions zebrafish host-microbiome responses to a later perturbation, with three predictions: that prior antibiotic and thermal exposure act as assembly filters, producing differences in gut community composition, diversity, and dispersion across stressor histories; that distinct histories alter the intestinal transcriptional response to a subsequent parasite challenge; and that cumulative mortality and infection prevalence among survivors both increase with prior stressor history. We reared 720 adult zebrafish across eight regimes spanning all combinations of antibiotic exposure, heat stress, and challenge with the intestinal nematode *Pseudocapillaria tomentosa*, applied sequentially (Figure 1; Table 1). Antibiotics and thermal stress represent increasingly common anthropogenic and climate-linked pressures on gut microbial communities [26, 27], while the nematode provides a biotic perturbation relevant to helminth infection of the vertebrate gut [28]. We then integrated gut microbiome composition, intestinal gene expression, and host outcomes to identify taxa associated with coordinated host-microbiome responses, clarifying how prior stressor history shapes vertebrate host-microbiome responses to future perturbation.

**Figure 1:**
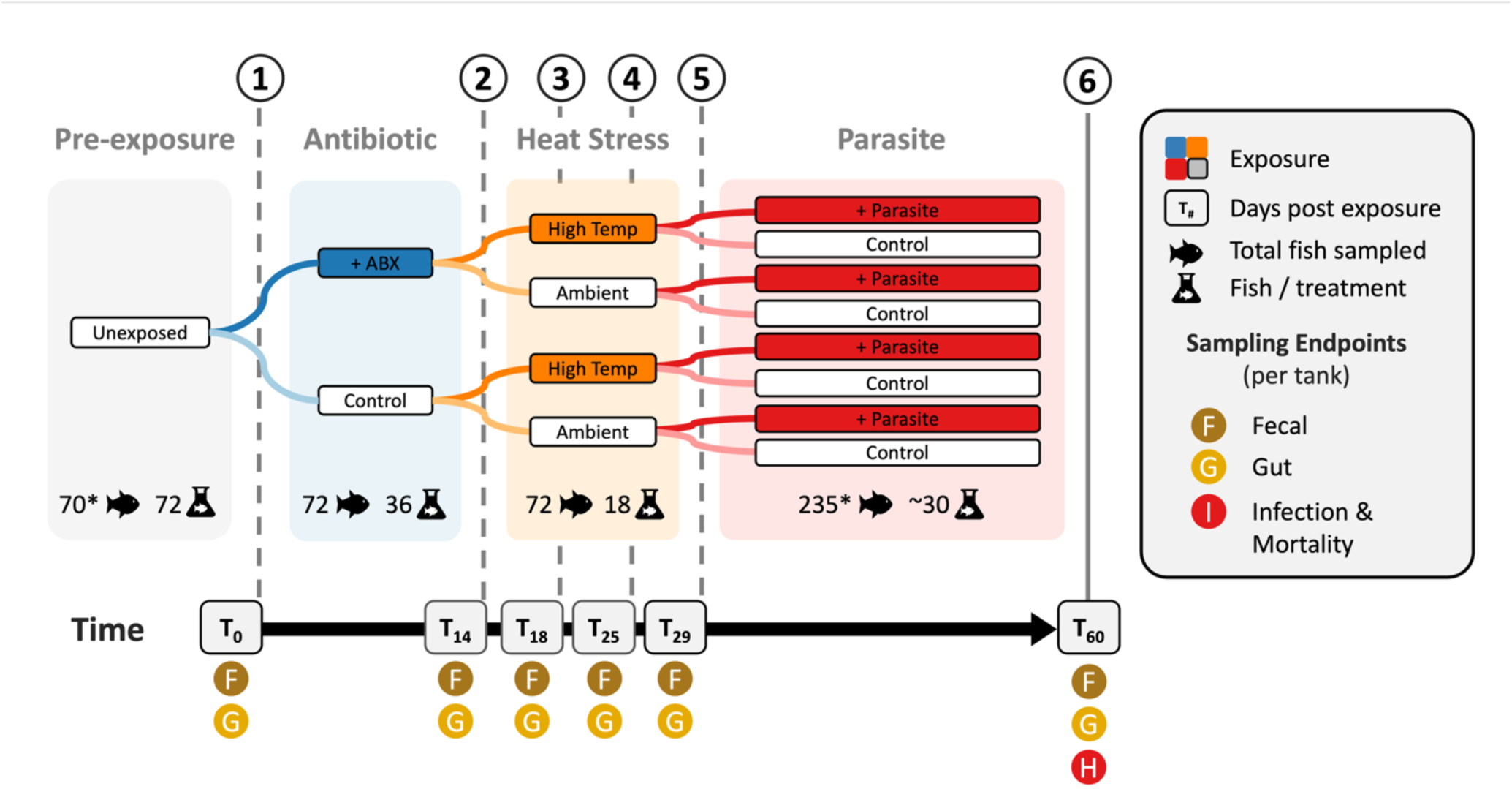
Experimental design schematic. Adult zebrafish (n = 720) were initially grouped into one of eight different cohorts across 24 tanks corresponding to one of the eight different pairwise combinations of exposure regimes, consisting of either no exposure (control), antibiotics (A+/-), temperature (T+/-; heat stress) and/or parasites (P+/-; *Pseudocapillaria tomentosa*) **(1-2)**. For the antibiotic stressor, antibiotic-exposed fish were exposed to antibiotics between days 0 and 14. **(2-5)** For the temperature stressor, **(3-4)** heat stressed fish were exposed to a heat stress between days 18 and 25, with a temperature **(2-3)** ramp up and **(4-5)** ramp down period (1-2°C/day) preceding and following the heat stress period between days 14 to 18 and days 25 to 29, respectively. For the parasite stressor, parasite-exposed fish were exposed to parasites between days 29 and day 60. **(1-5)** From days 0 to 29, fecal and gut microbiota samples were collected (n = 72 samples, or 3 samples per each of the 3 replicate tanks per each exposure regimes). **(6)** At the final time point (day 60), fecal and gut samples were collected from all the remaining surviving fish (n = 235). Additionally, cumulative final mortality was assessed, and surviving fish were inspected for presence of parasitic infection. Asterisks (“*”) indicate fewer samples than expected due to mortality or post-processing filtering due to insufficient read counts, and “∼” indicates average samples across treatments in the case of the final time point number of samples/treatment.

**Table 1:** Exposure regimes and prior stressor history. In addition to the control group, there were seven treatments consisting of different pairwise combinations of exposure regimes to antibiotics, heat stress, or parasite exposure. Prior to the final stressor of parasite exposure, fish in these different exposure regimes were successively exposed to a different number of prior stressor history, either zero, one, or two prior stressors.

| Exposure Regime | Prior Stressor History | Antibiotics | Heat Stress | Parasite |
| --- | --- | --- | --- | --- |
| A- T- P- | 0 | - | - | - |
| A- T- P+ | 0 | - | - | + |
| A+ T- P- | 1 | + | - | - |
| A+ T- P+ | 1 | + | - | + |
| A- T+ P- | 1 | - | + | - |
| A- T+ P+ | 1 | - | + | + |
| A+ T+ P- | 2 | + | + | - |
| A+ T+ P+ | 2 | + | + | + |

## Results

### Gut microbiota community displays historically contingent sensitivity in response to parasite exposure

We first characterized the gut microbiota at day 60 across the eight exposure regimes and prior stressor history levels (Figure 1; Table 1). Differential abundance analysis (MaAsLin3) identified 12 genera that differed from the control regime (A-T-P-) in at least one regime and 11 genera associated with a linear prior-stressor-history contrast (Benjamini-Hochberg q < 0.1; Tables S3.9.2, S3.9.7). Core zebrafish gut taxa were consistently present, with the most prevalent genera including *Aeromonas*, *Cetobacterium*, *Pseudomonas*, *Culicoidibacter*, *Shewanella*, and an unclassified *Paracoccaceae* genus (Figure 2A). Gut microbial diversity tracked prior stressor history more than parasite exposure. In factorial models, temperature was associated with Shannon and Simpson diversity, and an antibiotic-by-temperature interaction with observed richness (Table S1.1.1). Across the prior-stressor-history gradient, Simpson diversity declined significantly, Shannon showed a consistent but non-significant decline, and richness showed no trend (Table S1.5.1; Figure 2B). Parasite exposure did not significantly alter any diversity metric within a stressor-history level after correction (Table S1.2.1). Community composition was shaped by both factors. PERMANOVA associated prior stressor history and parasite exposure with composition under both Bray-Curtis and Canberra distances, with a significant history-by-parasite interaction only for Canberra (Table S2.9.1; Figure 2C). Beta dispersion also differed significantly across stressor-history levels (Table S2.3.1; Figure 2D). Because dispersion heterogeneity can itself drive PERMANOVA, we interpret prior stressor history as associated with both community location and dispersion, parasite exposure as associated with compositional shifts, and the history-dependent parasite effect as metric-dependent. Together, gut microbiota sensitivity to parasite exposure was shaped by prior stressor history.

**Figure 2:**
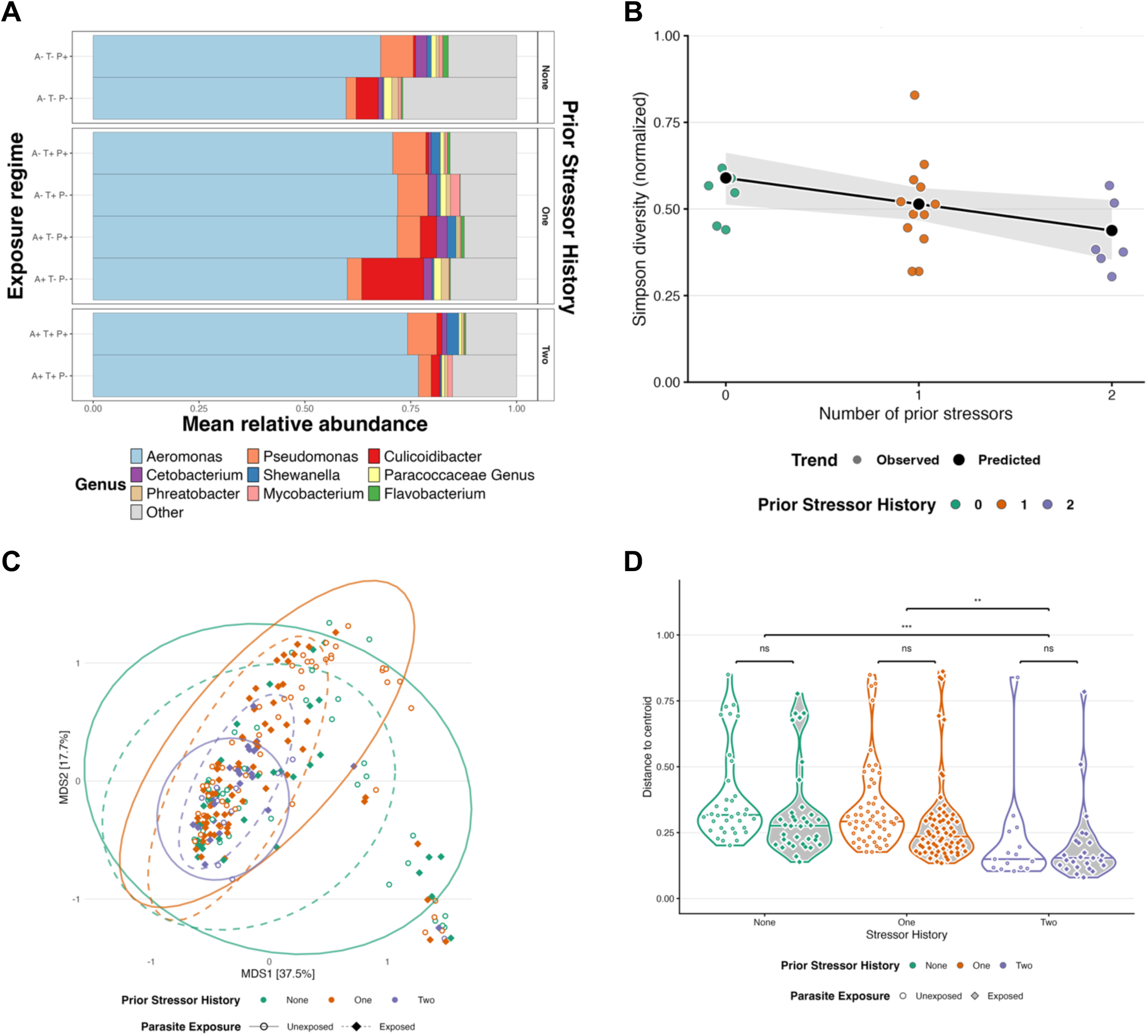
Exposure regimes and prior stress history restructure gut microbiome communities. A) Relative abundance of a subset of the most prevalent genera across exposure regimes and number of prior stressors. Genera are ordered by most to least prevalent. B) Gut microbial diversity displays a significant decreasing trend with increased prior stress history. C) PCoA of gut microbiota community composition varies across prior stressor history in the presence and absence of parasite exposure as measured by Bray-Curtis. D) Beta dispersion varies between fish exposed to different numbers of prior stressor but does not differ parasite exposed and unexposed fish with the same number of prior stressors. Asterisks indicate significance levels (* = p <0.05, ** p < 0.01, *** = p < 0.001; ns = not significant).

### Host gene expression shows non-additive response to parasite exposure

We next profiled intestinal host transcription across regimes at day 60. Because *Pseudocapillaria tomentosa* burrows into the intestinal lining, we expected parasite exposure to elevate differentially expressed gene (DEG) counts, and predicted that this response would scale linearly with prior stressor history. As expected, parasite-exposed fish showed far more DEGs than unexposed fish relative to controls (averaging 5,338 versus 78 DEGs; Benjamini-Hochberg adjusted P < 0.05; Figure 3; Table S4.3.1). Counter to prediction, however, DEG counts did not scale linearly: the response peaked at one prior stressor rather than two, whether quantified per regime (one-stressor parasite regimes averaged 6,283 DEGs versus 4,964 at two prior stressors; Table S4.3.1) or by pooled stressor-history group (9,188 DEGs at one versus 4,440 and 5,871 at zero and two; Table S4.2.1). Thus prior stressor exposure modifies the host transcriptional response to infection non-additively, with the largest response at intermediate stressor history.

**Figure 3:**
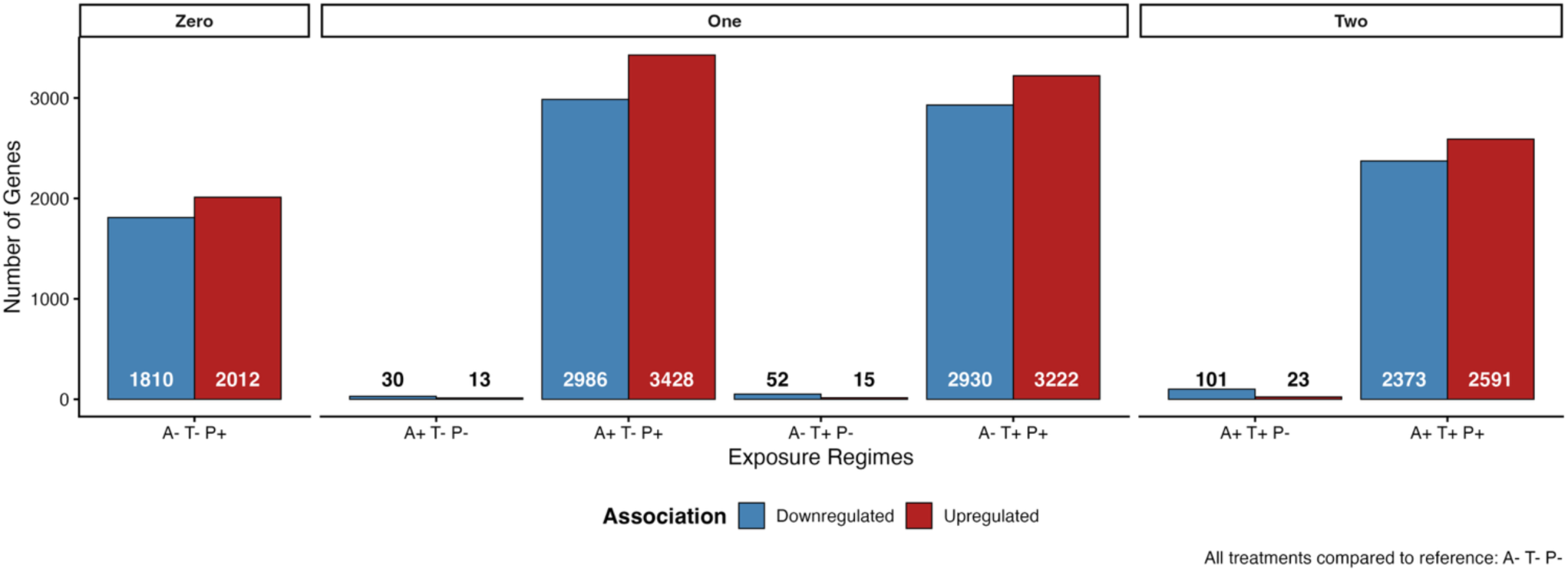
Total number of significant differentially expressed genes (DEGs) across exposure regimes as identified by DESeq2. Blue bars indicate total number of significant negatively (downregulate) associated DEGs and red bars indicate total number of significant positively (upregulated) associated DEGs. Exposure regimes are compared to the control group (A-T-P-). We find a non-additive trend in total number of significant DEGs and number of prior stressor history among fish exposed to parasites.

### Mortality increases with cumulative stressor exposure independent of parasite exposure

To test whether these patterns extended to host health, we examined mortality and infection at day 60. Because cohousing and intermediate terminal sampling preclude continuous mortality tracking, we estimated final mortality from the expected minus surviving fish per regime (about 45 fish expected per regime). Cumulative mortality rose from 18.9% at zero prior stressors to 31.1% at one and 55.6% at two, a nearly three-fold increase (Figure 4A), and a linear trend test confirmed that mortality increased significantly with prior stressor history (binomial GLMM with tank as a random effect; Table S5.13.1). This relationship held independent of parasite exposure, which had no significant main or interaction effect on mortality (Table S5.14.1). Infection prevalence among survivors did not increase with stressor history (83.8%, 59.7%, and 60.0% across zero to two prior stressors; Figure 4B; Table S5.8.1), and neither the linear trend nor pairwise contrasts were significant (Tables S5.5.1, S5.4.1). Mortality therefore increased with cumulative stressor history while infection prevalence did not, an unexpected dissociation we revisit below.

**Figure 4:**
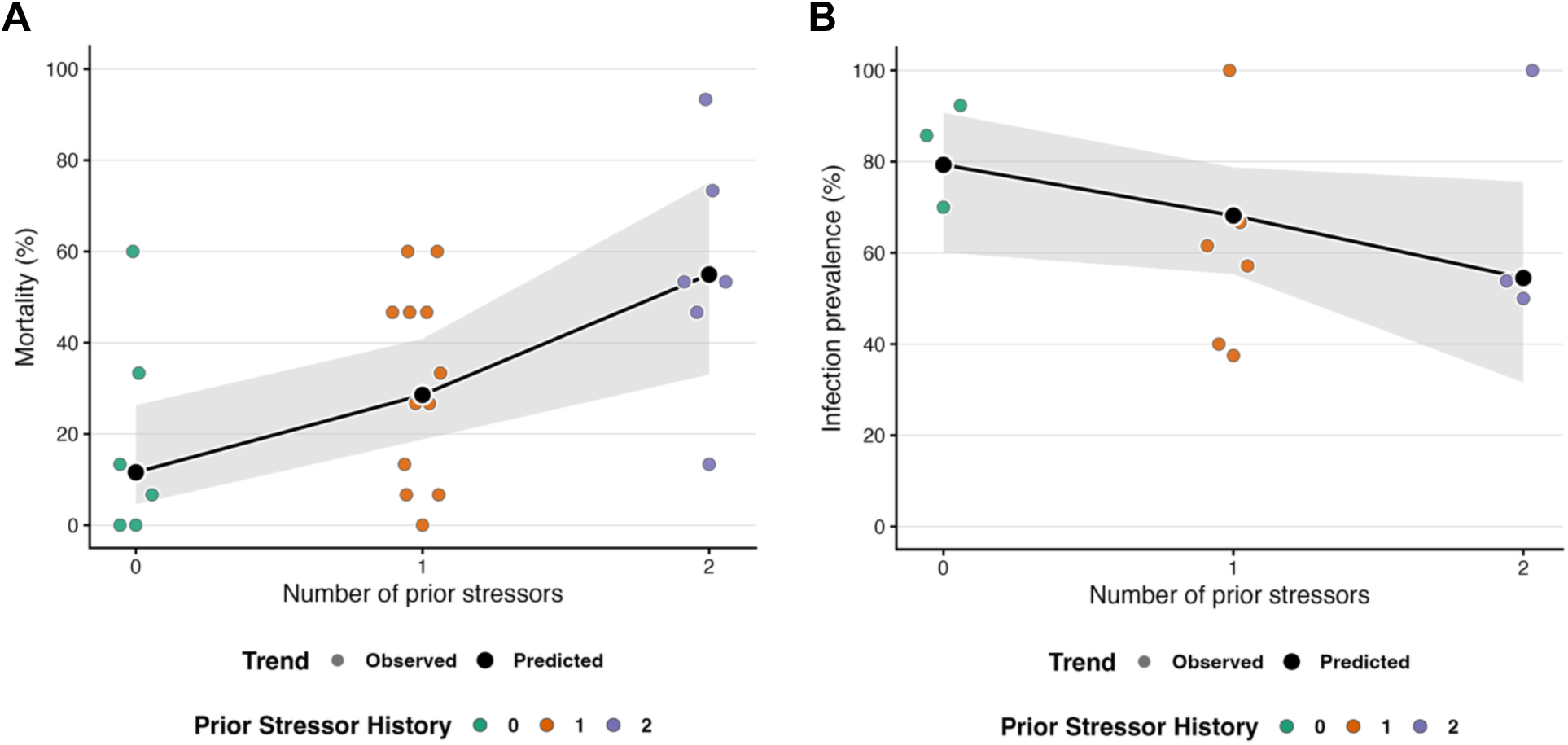
Final cumulative mortality and infection prevalence trends show a dissociation between mortality and infection prevalence trends. Trend analyses between prior stressor history and A) final mortality and B) infection prevalence at day 60. Across both plots, the colored points indicate observed mortality (n = 24 tanks) or infection prevalence (n = 12 tanks), and the black line and points show marginal model predictions, with 95% confidence intervals (gray ribbon). Mortality rose with prior stressor history while infection prevalence among survivors did not increase across the same gradient.

### *Culicoidibacter, Shewanella, Flavobacterium,* and *Cetobacterium* abundance associates with host gene expression and mortality

Building on these mortality patterns, we tested associations between genus abundance and tank-level final mortality (MaAsLin3 with tank as a random effect). Four genera, *Shewanella*, *Culicoidibacter*, *Flavobacterium*, and *Cetobacterium*, showed significant negative abundance-mortality associations (Figure 5A; Tables S3.8.1, S3.9.4), indicating that higher abundance of these taxa tracked lower final mortality.

**Figure 5:**
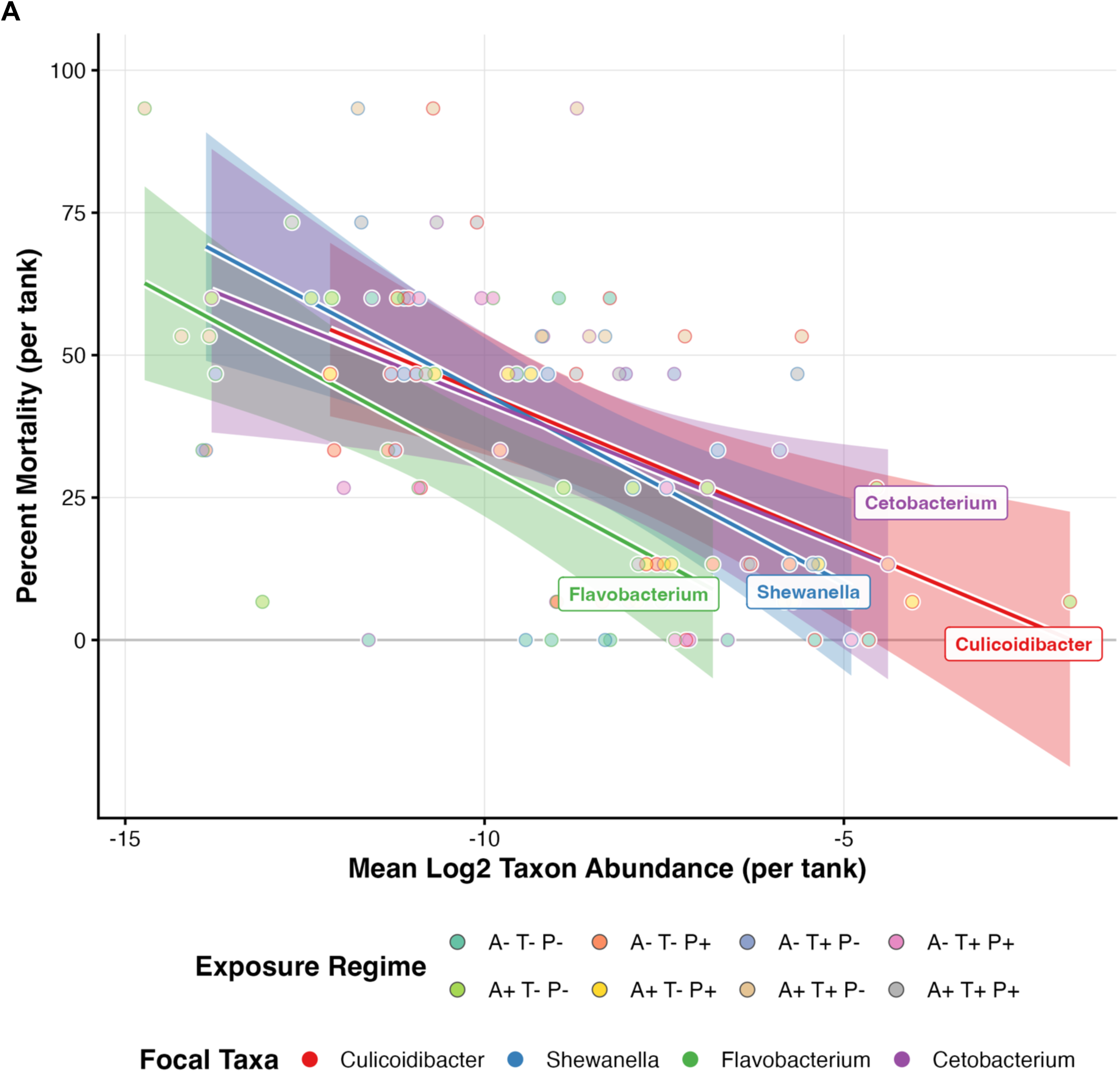
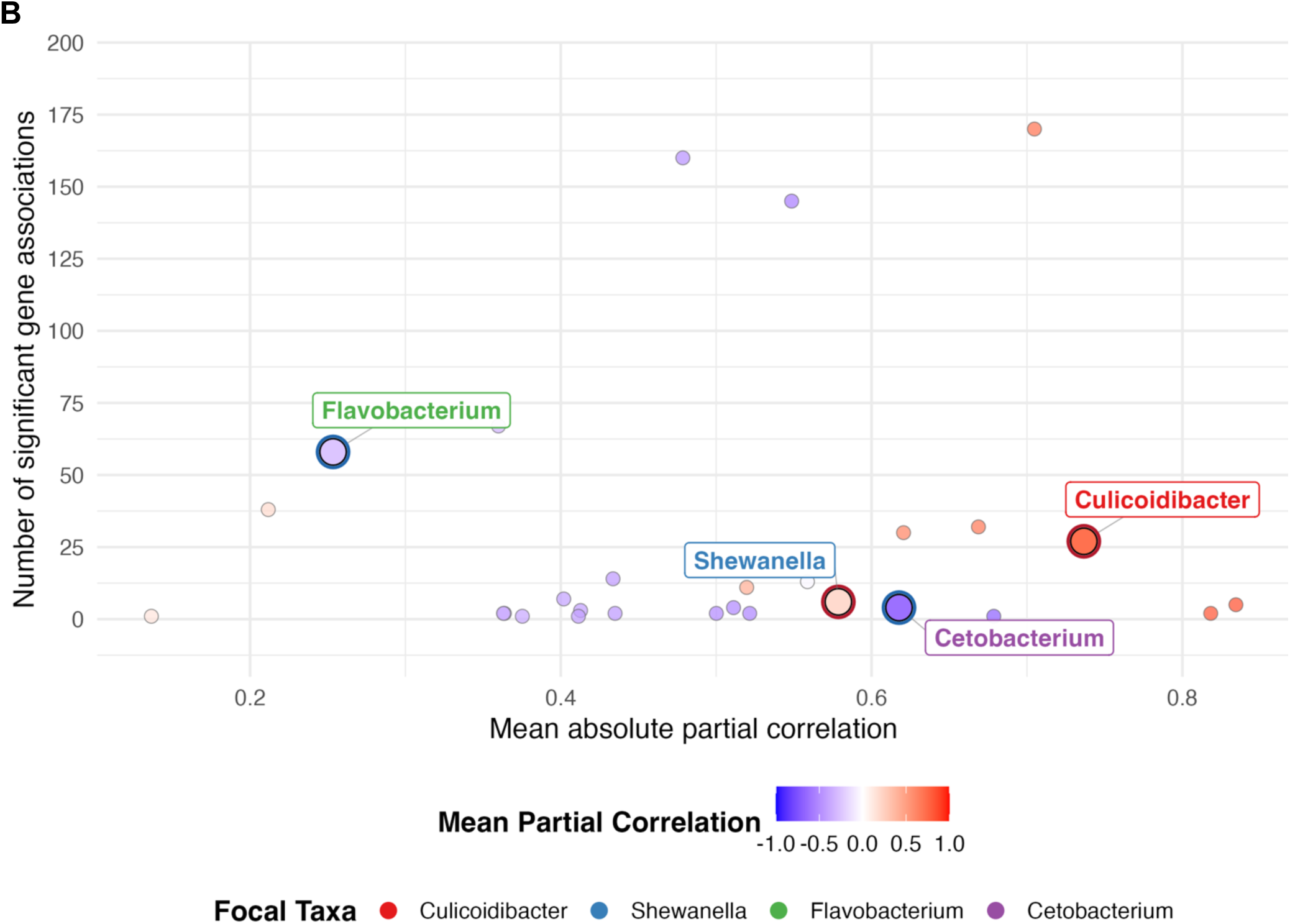
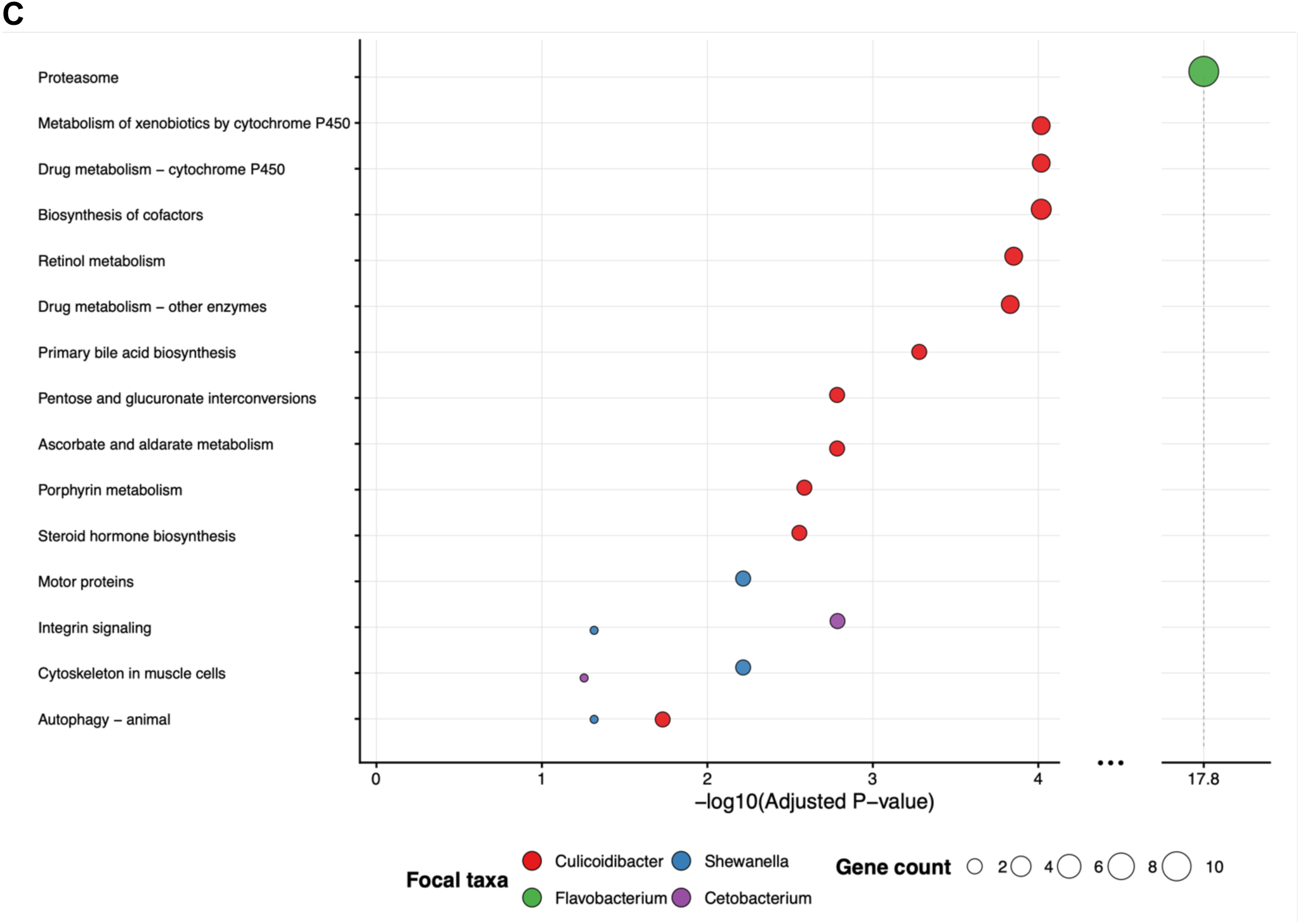
Culicoidibacter, Shewanella, Flavobacterium, and *Cetobacterium* associations with host final mortality and host intestinal gene expression. A) Tank-level final percent mortality versus log2 mean genus abundance of four focal taxa (*Culicoidibacter* (red)*, Shewanella* (blue), Flavobacterium (green), and *Cetobacterium* (purple) across exposure regimes and tanks. Dots are filled by exposure regime and outline color corresponds to taxon identity. Solid lines and ribbons represent model-based trends with 95% confidence intervals colored by taxon identity. **B)** Partial-correlation-based summary of associations between genus abundance (x-axis) and host intestinal gene expression (y-axis). Dots are filled by the mean sign of their significant partial correlations (blue = negative mean partial correlation, red = positive mean partial correlation). **C)** Dot plot showing aggregated enrichment results for the top 15 most significant KEGG pathways across the four focal taxa, dot color indicates focal taxa and dot size indicates the number of enriched genes in that taxon-pathway combination, and horizontal position indicates −log _10_(adjusted *P*-value). The x-axis is truncated between 5 and 17, indicated by a discontinuity marker (…).

We then asked whether these four genera also associated with host intestinal gene expression. Using a two-stage procedure (Spearman screening followed by nonparametric partial correlations adjusted for total worm burden, with Benjamini-Hochberg FDR control; Table S6.6.1), each of the four carried at least one significant gene edge. Their profiles differed: *Culicoidibacter* had few but the strongest edges (mean absolute partial correlation 0.744, nearly all positive), *Flavobacterium* the most edges (83, nearly all negative), and *Shewanella* and *Cetobacterium* intermediate numbers of comparatively strong edges (Figure 5B; Tables S6.5.1–S6.5.4). No host gene was shared among the four (Table S6.4.1). These genera were therefore consistently associated with both host gene expression and final mortality.

To identify host functional themes linked to each genus, we tested KEGG pathway and Gene Ontology over-representation on each genus’s significant partial-correlation gene list (Figure 5C; Tables S7.36.1–S7.39.1 for KEGG; Tables S7.17.1, S7.24.1, and S7.31.1 for GO). The four genera nominated largely distinct programs: *Flavobacterium*-linked genes were enriched for proteasome function, *Culicoidibacter*-linked genes for xenobiotic and drug metabolism via cytochrome P450, cofactor biosynthesis, and related metabolic pathways, and *Shewanella*-and *Cetobacterium*-linked genes for structural and signaling themes such as motor proteins, cytoskeleton, and integrin signaling. Despite non-overlapping input gene sets, each genus mapped to partitioned host functional themes rather than a shared program, consistent with distinct host-microbe relationships among the four taxa.

### Culicoidibacter, Shewanella, Flavobacterium, and Cetobacterium genera appear below Sloan neutral model expectations

Finally, we asked whether the abundances of these four genera reflected neutral processes (random dispersal, drift) or deterministic selection. We fit a Sloan neutral model to genus occurrence frequency versus metacommunity relative abundance at day 60 (Figure 6; Tables S8.6.2, S8.7.2, S8.8.2, and S8.9.2). All four focal genera fell below the neutral prediction, indicating that neutral processes alone cannot explain their distributions and that each occurred less frequently than expected for its abundance, consistent with being selected against. Together with their associations with mortality and host gene expression, these deviations provide a further line of evidence that *Culicoidibacter*, *Shewanella*, *Flavobacterium*, and *Cetobacterium* track biologically meaningful host-microbiome variation. Collectively, the gut microbiota responded to perturbation in a manner contingent on prior stress history, and these taxa may help mediate host outcomes within that historical context.

**Figure 6:**
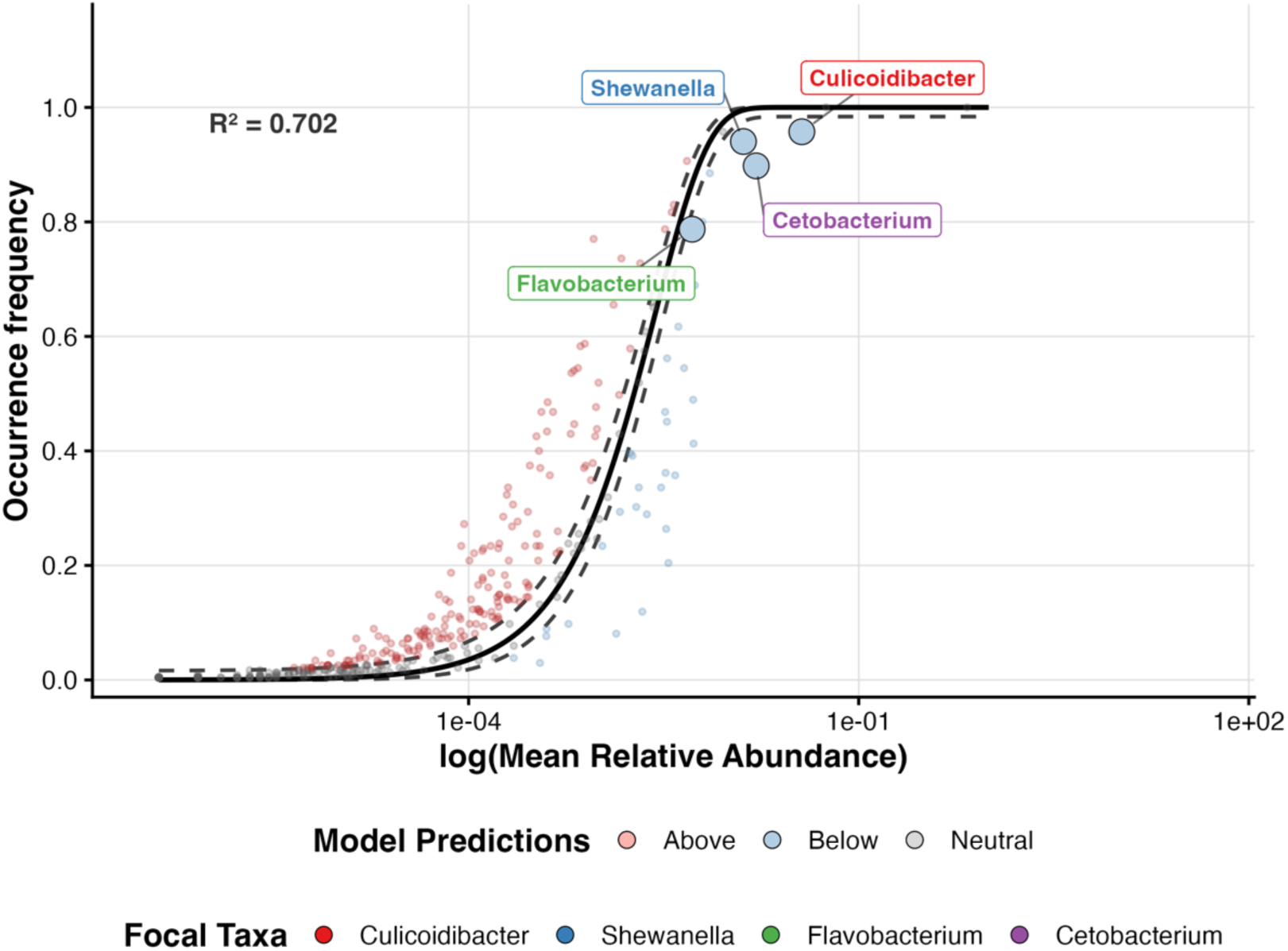
Fit of the Sloan neutral model at day 60. Each dot represents a unique genus across all samples at day 60. The y-axis represents occurrence frequency and the x-axis represents log10 mean relative abundance. Genera that occur more frequently than predicted by the Sloan neutral model are filled red, and those that occur less frequently are filled blue. The black dashed line indicates the model’s 95% confidence interval around the model prediction (solid black line). The focal taxa of *Culicoidibacter* (red)*, Shewanella* (blue)*, Flavobacterium* (green), and *Cetobacterium* (purple) genera occur below the Sloan neutral model’s predictions.

## Discussion

Ecosystems and the organisms that inhabit them face increasing environmental stressors that challenge their resiliency. Ecological theories of historical contingency and priority effects predict that an ecosystem’s prior stressor exposure, by altering community structure and function, shapes its response to future perturbations [12, 14], yet how strongly this applies to vertebrate host-microbiome systems remains unexplored [9, 29]. Here, we asked whether successive abiotic stressors (antibiotics and heat) restructure the zebrafish gut microbiome and condition host intestinal transcription and health outcomes to a subsequent biotic stressor (intestinal parasites).

Prior multi-stressor research spans many environments and host-associated systems [5, 30–33]; our study extends it by asking how multiple successive historical stressors shape a vertebrate host-microbiome system’s response to a later perturbation. We found that cumulative stressor history shapes the endpoint structure of zebrafish gut communities, consistent with priority effects and historical contingency, in which the order and timing of colonization influence community structure and function. Each prior stressor may act as an assembly and successional filter that narrows the set of viable community configurations (the species pool) able to withstand later stressors. Consistent with host traits contributing to this filtering, Stagaman et al. found that adaptive immunity filters the zebrafish gut microbiota by reducing neutral assembly (e.g., dispersal) and increasing interindividual differences, though such effects can be sensitive to shared-environment transmission [34]. Our observation that gut diversity decreases and composition converges with more prior stressors supports this view, indicating that microbiota structuring reflects historical exposure to successive stressors, not only the number of co-occurring ones.

How strongly parasite exposure registers at the community level depended on prior stressor history. Based on our earlier work with *P. tomentosa* [24, 35], we expected parasite exposure to produce larger differences in diversity and composition, especially in fish with more prior stressors. Instead, fish with fewer prior stressors showed greater interindividual heterogeneity, regardless of parasite exposure. This contrasts with the Anna Karenina Principle (AKP) [36], under which stressed hosts should be more variable, and with our prior work in which temperature acted as a stronger selective force and parasite exposure added stochasticity, with microbiota diverging toward multiple temperature-dependent stable states rather than an AKP-like dysbiosis [24]. The pattern instead fits a homeorhetic framework [37], in which stability reflects change along trajectories rather than return to a single static state. Two factors may explain the discrepancy with our expectations: parasite exposure here was chronic rather than single-dose, and, lacking intermediate sampling, endpoint patterns may be biased toward survivors. Thus prior stressor history conditions how parasite exposure is expressed at the community level, with endpoint structure reflecting both the final perturbation and the preceding sequence of stressors.

Host outcomes also tracked cumulative stress history, indicating organism-level consequences of sequential perturbation. As predicted, mortality rose with prior stressor history, consistent with work showing that multiple stressors alter host fitness [38–40] and that microbiome context shapes responses to combined stressors [19, 41]. In *Daphnia*, for example, the interaction between a cyanobacterial stressor and an oomycete parasite depended on microbiome inoculum type, with an antagonistic survival effect only in hosts carrying a laboratory-derived microbiome [19]. We similarly observed a non-significant antagonistic relationship in which parasite-exposed fish had lower mortality than unexposed fish of similar stressor history, though this differs from Godwin et al., who found heat stress exacerbates mortality in parasite-exposed salmon [38]. The hygiene hypothesis offers one interpretive frame, whereby limited exposure to immune-stimulating organisms can promote dysregulated immunity and vulnerability to later challenges [42, 43]. Together these results suggest multi-stressor survival effects are context-dependent and non-additive, not predictable from single-stressor responses. Contrary to our third prediction, infection prevalence among survivors decreased with prior stressor history. We interpret this as selective mortality: fish most vulnerable to cumulative stress and parasite exposure likely died before final sampling, leaving a more resilient surviving subset and producing an inverse mortality-infection relationship. This implies a host-level filtering effect analogous to the microbiome filtering above, in which successive stressors constrain not only which microbiota configurations persist but also which hosts survive to be assessed.

Host intestinal transcription was likewise historically contingent and non-additive. Parasite-associated differential gene expression (DEG) peaked in fish with one prior stressor and was lower at zero or two, so the response did not scale linearly with cumulative stress. Among parasite-unexposed fish, transcriptional divergence from the reference regime instead increased with prior stressor number, indicating that cumulative stress alone progressively altered the intestinal state, and our joint model identified genes with significant parasite-by-stress-history interactions, supporting a host response that depends on prior context. Because gut microbiota and host intestinal gene expression are tightly linked through immune, metabolic, and barrier processes, and microbiome state can influence transcriptional plasticity and stress tolerance [41, 44], the non-additive response we observed may partly reflect differences in host-microbiome state across stress-history backgrounds. However, sampling transcription only at the endpoint means we cannot resolve expression trajectories or separate direct effects of cumulative stress, microbiome-associated shifts, selective mortality, and their interactions. Overall, prior stress history appears to coordinate shifts in both microbiome structure and host response, prompting us to ask which microbial taxa link these patterns to host outcomes.

Integrating microbial abundance with host gene expression and mortality highlighted four gut taxa as candidate host-linked members. Lower abundance of *Cetobacterium*, *Culicoidibacter*, *Flavobacterium*, and *Shewanella* was associated with higher mortality, and each showed significant partial-correlation associations with intestinal gene expression after adjusting for worm burden. All four are common zebrafish gut residents [21, 22, 45, 46]. *Culicoidibacter* had few but particularly strong gene edges, enriched for xenobiotic and drug metabolism and heme- and oxidoreductase-related functions plausibly relevant to stress tolerance; it corresponds to the ZOR0006 lineage reported in zebrafish [47] and to a representative characterized from *Culicoides sonorensis* larvae [48], and although its gut role is poorly resolved, its recurrence and strong host associations make it a follow-up candidate. *Cetobacterium* was linked to a small gene set implicating tissue remodeling and stress-response programs, consistent with prior evidence that it modulates host metabolism and improves zebrafish health as a probiotic [49–51]. *Shewanella* likewise showed gene-expression associations, and gnotobiotic and community work indicates it can modulate intestinal immune cell recruitment in zebrafish larvae [52, 53], supporting a context-dependent host-interactive role.

*Flavobacterium*, by contrast, was linked to the largest host-gene set among the four, with predominantly negative associations enriched for proteasome- and ubiquitin-associated protein catabolism, RNA processing and splicing, and clathrin adaptor and coat complexes. This requires caution: *Flavobacterium* is a diverse genus that includes fish-associated opportunistic pathogens [54, 55], some reported in zebrafish [56], so the association between lower abundance and higher mortality is not evidence of a uniformly beneficial role. It may instead be a context-dependent member whose abundance reflects host physiological state, community stability, or strain-level differences. Consistent with this, our partial correlations identify single-taxon associations after worm-burden adjustment but do not resolve higher-order interactions among taxa, so reduced *Flavobacterium* may signal broader disruption of community structure rather than an isolated protective effect, making it a possible biomarker of host or community state whose pathogenic outcomes arise only under particular conditions. Together, these analyses highlight *Cetobacterium*, *Culicoidibacter*, *Flavobacterium*, and *Shewanella* as candidate host-linked members whose abundance patterns are consistent with keystone-like roles, context-dependent host interactions, or use as biomarkers of host state under cumulative stress. Targeted follow-up will be needed to distinguish whether they contribute causally to host outcomes, reflect host physiological condition, participate in community interaction networks, or some combination.

Neutral-community modeling provided a complementary test of whether the four taxa’s occurrence patterns were consistent with neutral assembly. Comparing each taxon’s metacommunity occurrence to Sloan neutral predictions, as applied previously to the zebrafish gut [22], all four fell below expectation, occurring less frequently than predicted from their mean relative abundance. Neutral processes such as dispersal and drift are therefore insufficient to explain their distributions, which may instead be shaped by non-neutral processes including host or environmental filtering, stressor-associated niche constraints, competition, or priority effects from prior community history. Alongside their associations with mortality and host gene expression, these deviations add evidence that *Cetobacterium*, *Culicoidibacter*, *Flavobacterium*, and *Shewanella* track biologically meaningful host-microbiome variation. These analyses do not, however, establish whether reduced occurrence contributed to poor outcomes, resulted from stress-associated changes in the gut environment, or reflected broader community restructuring; resolving this will require temporal sampling, mono- and co-colonization experiments, and defined-community perturbations.

Collectively, host-microbiome responses to perturbation were historically contingent on prior stressor history. Integrating gut microbiota, host gene expression, health outcomes, and neutral-model analyses, we nominate *Cetobacterium*, *Culicoidibacter*, *Flavobacterium*, and *Shewanella* as candidates for follow-up, either as putative keystone-like taxa or as biomarkers of host state under cumulative stress. These results argue that prior stressor history, not only concurrent stressor combinations, should be considered when interpreting multi-stressor experiments in host-associated systems, and that stressor sequence should be incorporated when forecasting host-microbiome responses to Anthropocene-linked pressures. Testing whether specific taxa causally shape host outcomes, and building predictive models, will require temporal sampling and functional assays.

## Methods

### Ethical approval

All animal procedures were approved by the Oregon State University Institutional Animal Care and Use Committee (IACUC protocol 2022-0280) and conducted in accordance with institutional and national guidelines for the care and use of laboratory zebrafish.

### Experimental design and stressor history

We tested whether sequential environmental stressor history conditions host-microbiome responses to a later biotic challenge in adult zebrafish (*Danio rerio*, 5D strain). Seven hundred twenty fish were distributed across 24 tanks (three replicate tanks per treatment) in a fully factorial design crossing antibiotic exposure (A+ or A-), heat stress (T+ or T-), and intestinal nematode exposure (*Pseudocapillaria tomentosa*; P+ or P-), yielding eight exposure regimes (A±T±P±). Stressors were applied sequentially over 60 days rather than concurrently, so that regimes differed in the number of prior stressors experienced before the final parasite phase (0, 1, or 2 prior stressors; Table 1).

Antibiotic-exposed fish received medicated gelatin feed containing streptomycin, ciprofloxacin, and ampicillin (50 µg g⁻¹ body weight per day) from days 0-14. Heat-stressed fish experienced a ramp from 28 °C to 35 °C between days 14-18, maintenance at 35 °C through day 25, and return to 28 °C by day 29. Parasite-exposed fish received continuous exposure to *P. tomentosa* eggs via flow-through effluent from infected donor tanks from days 29-60, following established Sharpton and Kent Lab helminth challenge procedures [24, 25, 28]. Control fish (A-T-P-) experienced matched handling without antibiotics, without heat ramp, and without parasite effluent.

### Animal husbandry

Fish were reared under standard Sharpton and Kent Lab zebrafish husbandry (Supplementary Methods): recirculating aquaculture, 14:10 h light:dark cycle, and diet consistent across tanks unless otherwise noted for antibiotic delivery. Tanks were cohoused by exposure regime; individual fish were not uniquely tagged. This cohousing structure motivated endpoint-only mortality estimation and tank-level inferential models (see below).

### Sampling and host phenotyping

Fecal and intestinal microbiota samples were collected at days 0, 14, 29, and 60 (three fish per tank per time point when available). At day 60, all remaining survivors were sampled, yielding 235 fish with sufficient 16S data after quality filtering. Intestinal tissue for RNA sequencing was collected from a subset of day-60 fish with paired microbiome data (Supplementary Methods).

Cumulative mortality was assessed at day 60 only. Because intermediate sampling removed fish from tanks and fish were cohoused without individual IDs, daily mortality could not be tracked continuously. We therefore estimated final cumulative percent mortality per exposure regime as:

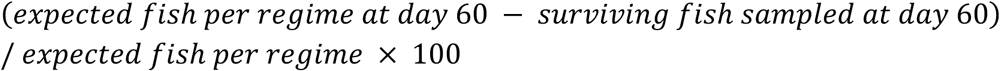

where expected fish per regime reflects starting stocking (90 fish per regime across three tanks) minus documented removals at intermediate sampling time points (∼45 fish expected per regime at day 60 after intermediate terminal sampling). Tank was the unit of observation for mortality models (24 tanks).

Infection prevalence and worm burden were assessed for all fish. No infection was detected among parasite unexposed fish, so infection prevalence and worm burden were quantified among parasite-exposed (P+) day-60 survivors only. Unexposed (P-) fish were not included in infection prevalence or burden summaries because they were not challenged with *P. tomentosa*. For each P+ tank with day-60 survivors, infection prevalence was computed as infected fish divided by sampled survivors. Worm burden was summarized descriptively as mean total worm count per sampled P+ survivor and, separately, mean burden among infected fish only. Infection prevalence was modeled with tank-level binomial GLMMs on P+ day-60 survivors (∼12 P+ tanks with survivors at endpoint). Parasite exposure (P+ vs P-) entered mortality models as a binary fixed effect but was not used as a worm-burden covariate in infection summaries.

### 16S rRNA gene sequencing and microbiome processing

Gut microbiome profiling followed our published zebrafish 16S workflow (Sieler et al. 2023, 2025; see Supplementary Methods for PCR, sequencing platform, DADA2 parameters, taxonomy assignment, and filtering). Briefly, V4-region amplicons were processed to amplicon sequence variants (ASVs), assigned with the SILVA reference, and aggregated to genus level for downstream analyses. Analyses presented in the main text focus on day-60 samples unless noted.

### RNA sequencing and differential gene expression

Intestinal RNA was extracted from day-60 samples and processed with standard Illumina library preparation and paired-end sequencing (Supplementary Methods). Salmon quantification yielded gene-level counts. We conducted two complementary DESeq2 designs on filtered genes: (1) per exposure-regime contrasts versus the A-T-P-reference (eight regimes; main-text Figure 3 spine), and (2) a pooled-by-prior-stressor-history model grouping parasite-exposed fish by history level (0, 1, or 2 prior stressors) to test history-dependent transcriptional responses (Table S4.2.1). Significant differentially expressed genes (DEGs) were declared at Benjamini-Hochberg adjusted *P* < 0.05. The A-T-P-reference regime was omitted from bar-chart display because it is the contrast baseline.

### Microbiome community analyses

#### Alpha diversity

Genus-level Shannon diversity [57], inverse Simpson diversity [58], and observed genus richness were analyzed with beta regression GLMMs (glmmTMB [59]; logit link) including random intercepts for tank: y ∼ HistoryLevelNum × Parasite + (1 | Tank.ID) for stress-history gradients, and factorial Antibiotics × Temperature × Parasite models with tank random effects for exposure-regime effects. Type-III Wald tests and emmeans contrasts were used for fixed effects [60, 61]; pairwise parasite contrasts within each history level received Benjamini-Hochberg false discovery rate (FDR) correction across strata [62, 63].

#### Compositional variation and dispersion

Bray-Curtis [64] and Canberra [65] dissimilarities among genus-relative-abundance profiles were tested with PERMANOVA (distance ∼ HistoryLevelNum × Parasite; 999 permutations, seed 42) and betadisper ANOVAs on group dispersions [66, 67]. Main-text models treat fish as exchangeable units under permutation (no tank stratum).

#### Differential abundance

Genus-level differential abundance at day 60 was estimated with MaAsLin3 [68] (TSS normalization, log transform) using tank random-effect models for primary manuscript tables: exposure-regime contrasts versus A-T-P-, linear prior-stressor-history trends, and parasite effects (∼ Treatment + (1 | Tank.ID) or ∼ HistoryLevelNum + (1 | Tank.ID) as appropriate). Cited main-text results use tank models (e.g., Tables S3.8.1, S3.9.2, S3.9.4, S3.9.7). Sensitivity models without tank random effects are retained in supplements where noted.

### Host-microbiome integration

#### Taxon-mortality associations

Tank-level final percent mortality was regressed on genus log₂ abundance with MaAsLin3 [68] including (1 | Tank.ID) to identify taxa whose abundance tracked mortality (Table S3.8.1).

#### Partial correlation networks

We tested associations between genus relative abundance and host intestinal gene expression using a two-stage procedure on day-60 samples with paired RNA and 16S data. First, Spearman correlations were screened across exposure-regime pairwise contrasts versus A-T-P-; second, retained gene-taxon pairs were tested with nonparametric partial correlations (nptest::np.cor.test) adjusting for total worm count as a continuous covariate, as previously described [25]. Partial-correlation *P* values were FDR-controlled at 0.10 for network edges (Table S6.6.1). Worm burden was set to zero for unexposed (P-) RNA samples with missing burden (structural zeros); mortality and infection models instead use parasite exposure as a binary indicator and P+-only subsets for burden summaries, respectively (see Sampling and host phenotyping).

#### Functional enrichment

For each focal genus with significant partial-correlation edges, host genes passing the partial-correlation FDR threshold were tested for over-representation of KEGG pathways and Gene Ontology terms (biological process, molecular function, cellular component) using clusterProfiler (org.Dr.eg.db; organism dre). Enrichment used the submitted gene list per genus without an explicit genome-wide expression universe (Supplementary Methods); interpretation focuses on ranked terms (Figure 5C; Tables S7.36.1–S7.39.1 for KEGG; Tables S7.17.1, S7.24.1, and S7.31.1 for GO).

#### Sloan neutral community model

At day 60 we fit a Sloan neutral model relating genus occurrence frequency to metacommunity mean relative abundance [69, 70], with bootstrap confidence intervals for the neutral expectation (1,000 resamples, seed 42). Focal genera were compared to the model prediction to classify occurrence above or below neutral expectation (Figure 6; Tables S8.6.2, S8.7.2, S8.8.2, and S8.9.2).

#### Significance testing and reporting

Unless stated otherwise, two-sided tests were evaluated at α = 0.05. Multiple testing was controlled within analysis families by Benjamini-Hochberg FDR: alpha-diversity Wald and pairwise contrasts; PERMANOVA and betadisper follow-up tests where noted; MaAsLin3 qval_individual < 0.05 for primary taxon screens (joint q < 0.1 reported for multi-feature MaAsLin summaries in supplements); DESeq2 adjusted *P* < 0.05 for DEGs; partial-correlation FDR < 0.10 for network edges (Spearman screen FDR < 0.05); enrichment adjusted *P* < 0.10 for reporting ranked pathways. Main-text Results report effect direction and magnitude; test statistics, degrees of freedom, and exact *P* values appear in cited supplementary tables.

#### Mortality trend contrast

Tank-level mortality was modeled as cbind(dead, alive) ∼ HistoryLevelNum + (1 | Tank.ID) (main trend) and cbind(dead, alive) ∼ HistoryLevelNum × Parasite + (1 | Tank.ID) (joint test). The reported odds ratio for the history gradient is from a linear polynomial contrast on the log-odds scale (emmeans::contrast(…, “poly”)) testing monotonic increase across history levels 0 → 1 → 2. This OR is not equivalent to a simple two-level marginal odds ratio between extreme groups on the response scale (Table S5.13.1).

## Supporting information

Supplementary Tables and Figures Index

## Data availability

Raw 16S and RNA-seq reads are available from NCBI BioProject PRJNA1482558 (SRA). All code and associated files can be found on the project github page: https://github.com/sielerjm/zebrafish-stress-contingency-2026.

## Author Contribution

TJS and MLK conceived and designed the study. MJS, CL, and KK conducted the experiments. MJS and TJS performed the gut microbiome and integrated analyses. MJS prepared the figures. MJS, TJS, CL, KK, and MLK contributed to the preparation and editing of the manuscript. All authors read and approved the final manuscript.

