## Supplementary Figures for "Historical contingency shapes zebrafish host-microbiome responses to a subsequent biotic challenge": Supplementary_Figures__01__Diversity__submission__2026-06-26.pdf

### Supplementary figures (combined)

Build (UTC): 2026−06−26 12:36:43 UTC

Git: 55d6b9a4a7bf134b3db15f0777c389e2da8c31be

Module: 01__Diversity

Figures dir: /Sieler2026/Results/01__Diversity/Figures

Figures (combined PDF order; p. = first page of that figure in this PDF):

p. 2: Figure S1.1.1 − alpha_factorial_ATP_richness

p. 3: Figure S1.1.2 − alpha_factorial_ATP_shannon

p. 4: Figure S1.1.3 − alpha_factorial_ATP_simpson

p. 5: Figure S1.6.1 − observed_diversity_trend_quasirandom

p. 6: Figure S1.7.1 − richness_parasite_within_history

p. 7: Figure S1.8.1 − shannon_diversity_trend_quasirandom

p. 8: Figure S1.9.1 − shannon_parasite_within_history

p. 9: Figure S1.10.1 − simpson_diversity_trend_quasirandom

p. 10: Figure S1.11.1 − simpson_parasite_within_history

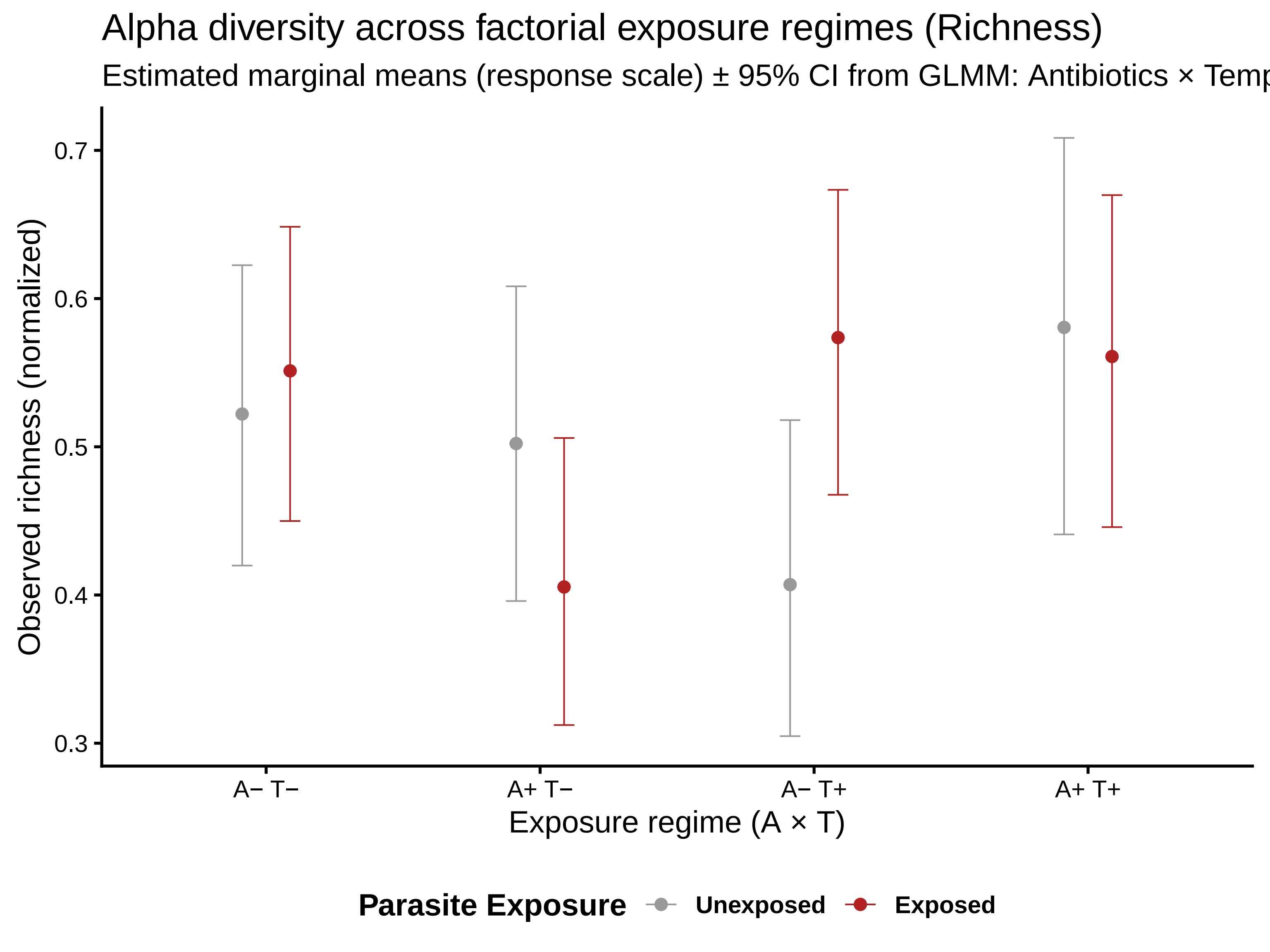

Alpha diversity across factorial exposure regimes (Richness)
Estimated marginal means (response scale) ± 95% CI from GLMM: Antibiotics × Temp
Observed richness (normalized)
0.7
0.6
0.5
0.4
0.3
A− T−
A+ T−
A− T+
A+ T+
Exposure regime (A × T)
Parasite Exposure
Unexposed
Exposed

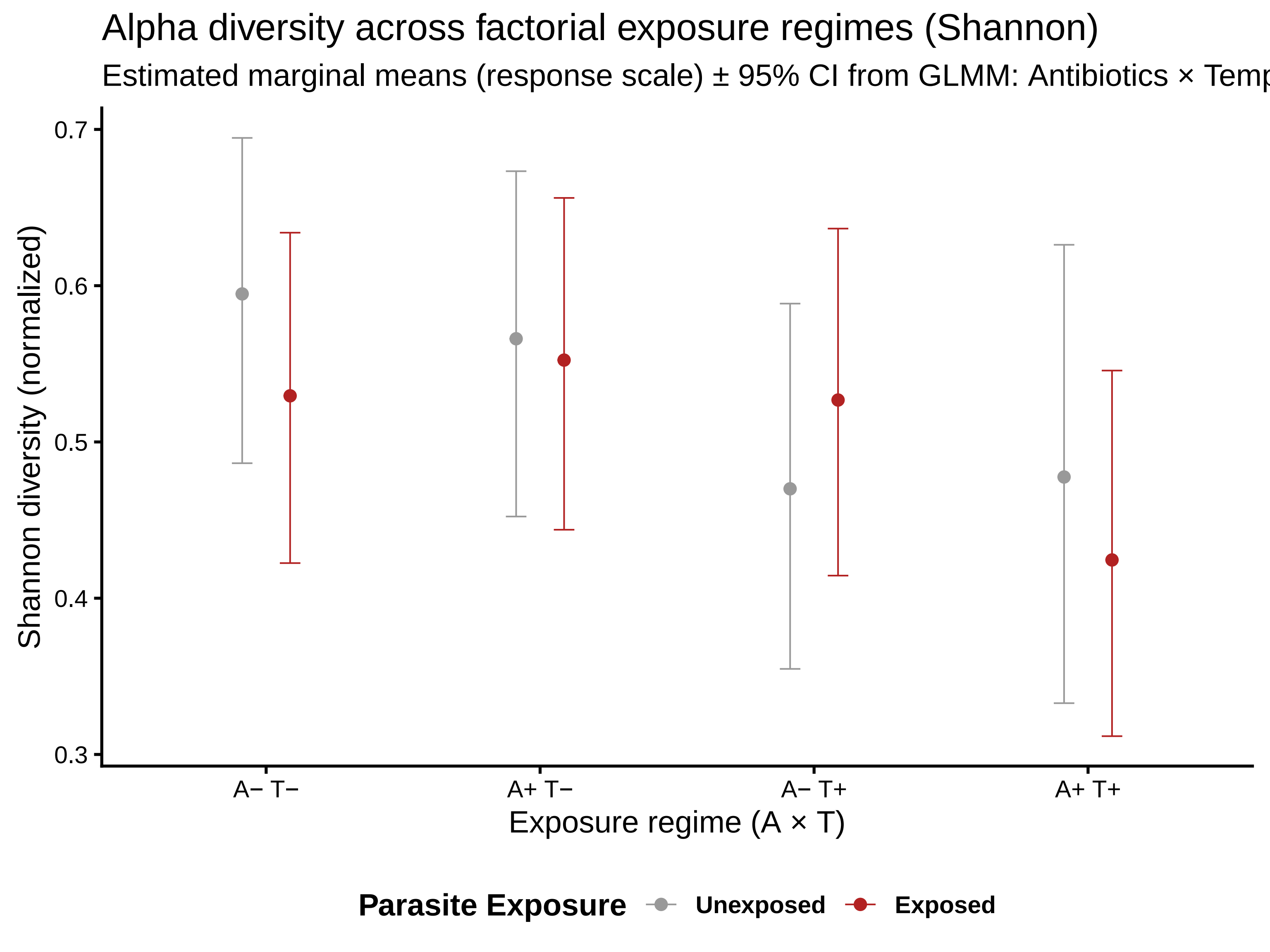
Alpha diversity across factorial exposure regimes (Shannon)
Estimated marginal means (response scale) ± 95% CI from GLMM: Antibiotics × Temp
Shannon diversity (normalized)
0.7
0.6
0.5
0.4
0.3
A− T−
A+ T−
A− T+
A+ T+
Exposure regime (A × T)
Parasite Exposure
Unexposed
Exposed

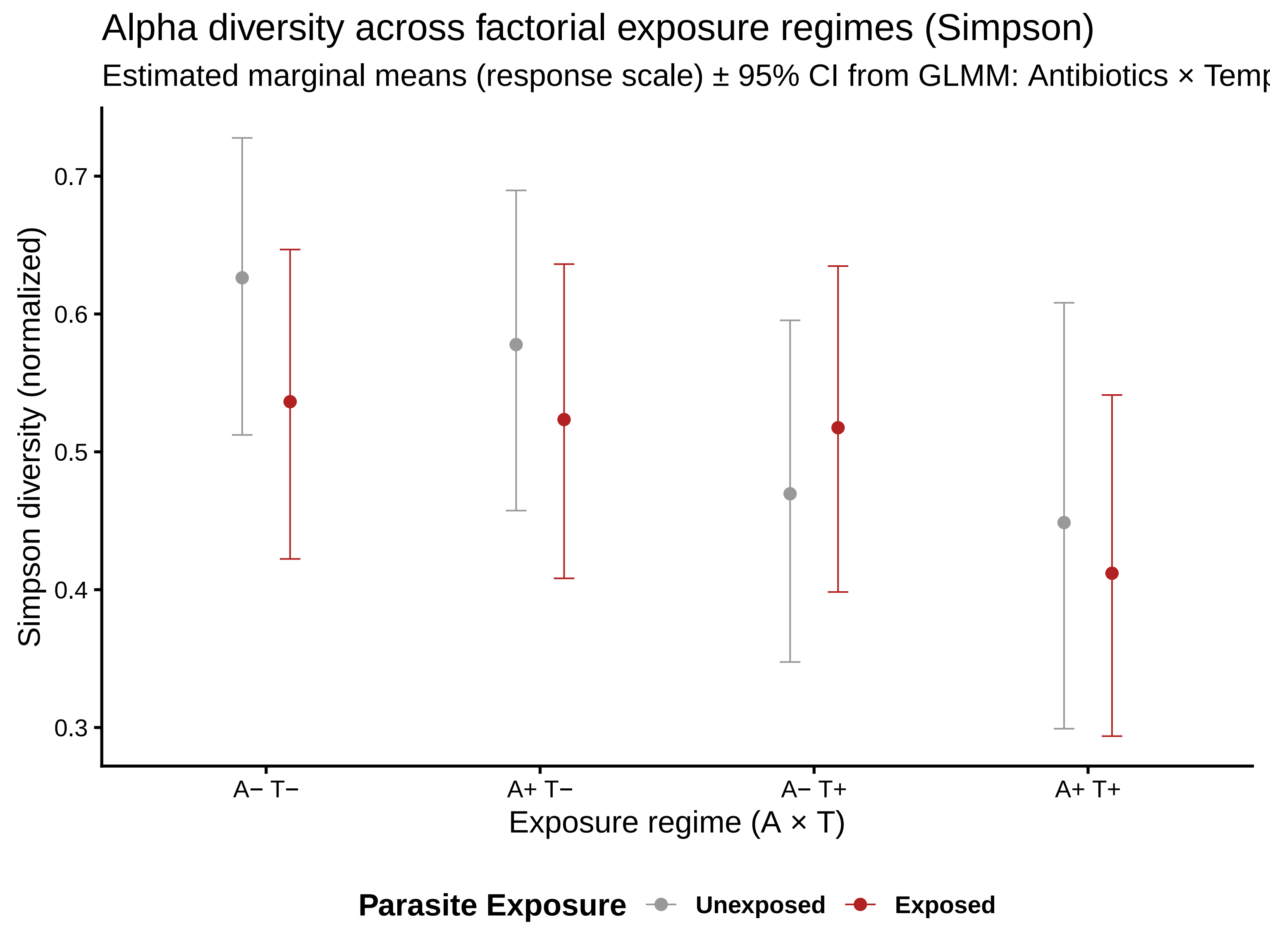
Alpha diversity across factorial exposure regimes (Simpson)
Estimated marginal means (response scale) ± 95% CI from GLMM: Antibiotics × Temp
Simpson diversity (normalized)
0.7
0.6
0.5
0.4
0.3
A− T−
A+ T−
A− T+
A+ T+
Exposure regime (A × T)
Parasite Exposure
Unexposed
Exposed

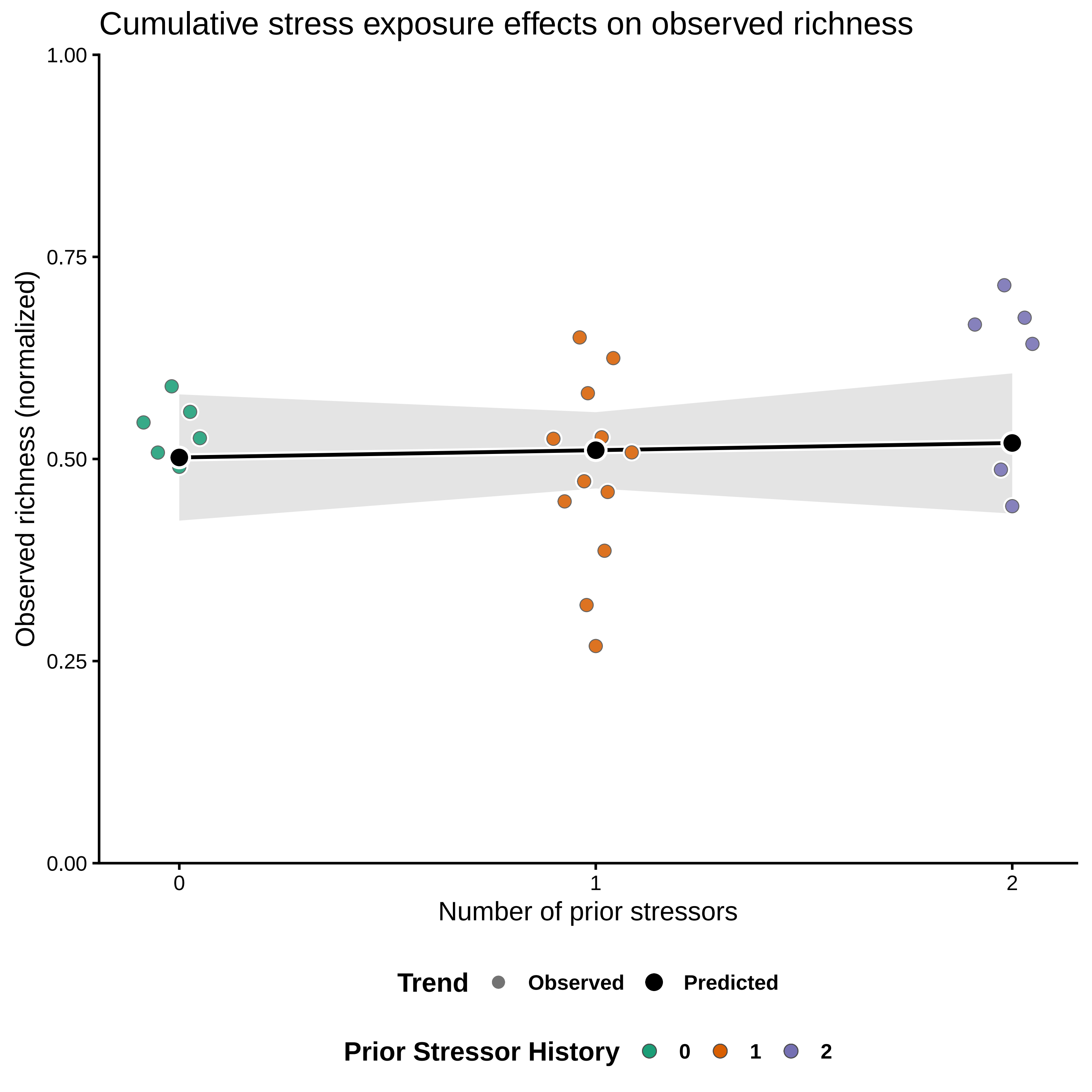
Cumulative stress exposure effects on observed richness
1.00
0.75
0.50
0.25
0.00
Observed richness (normalized)
0
1
2
Number of prior stressors
Trend
Observed
Predicted
Prior Stressor History
0
1
2

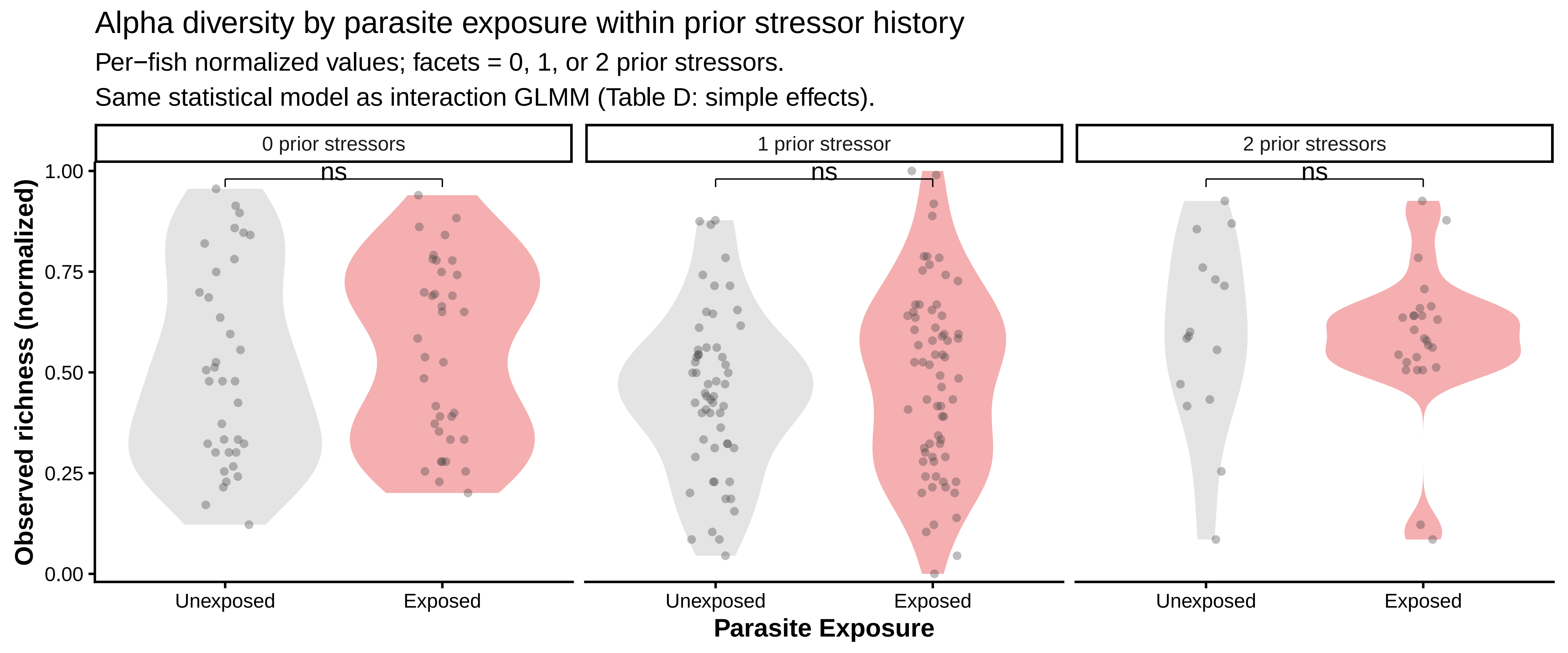

Alpha diversity by parasite exposure within prior stressor history
Per−fish normalized values; facets = 0, 1, or 2 prior stressors.
Same statistical model as interaction GLMM (Table D: simple effects).
0 prior stressors
1 prior stressor
2 prior stressors
ns
ns
ns
Observed richness (normalized)
1.00
0.75
0.50
0.25
0.00
Unexposed
Exposed
Unexposed
Exposed
Unexposed
Exposed
Parasite Exposure

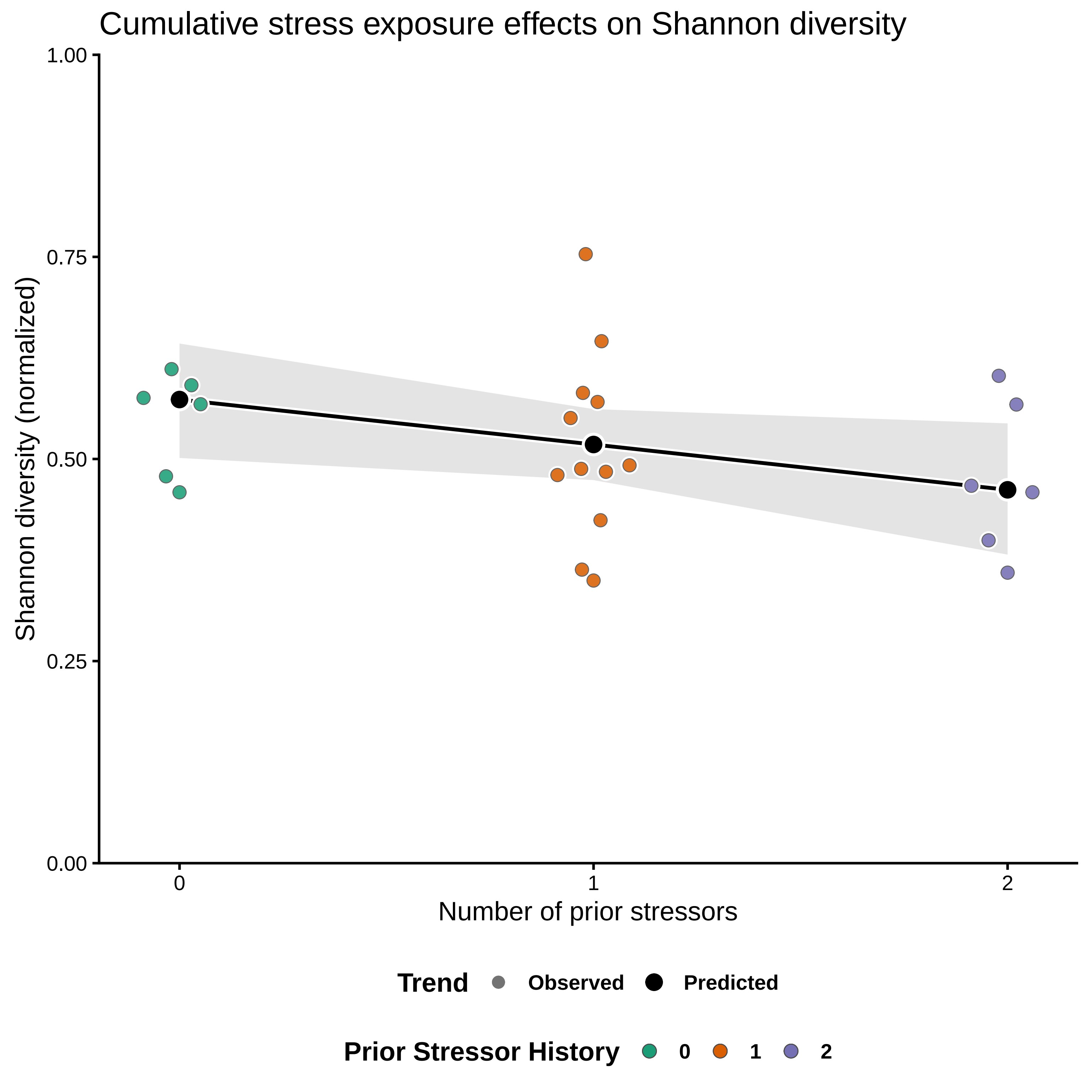

Cumulative stress exposure effects on Shannon diversity
Shannon diversity (normalized)
1.00
0.75
0.50
0.25
0.00
0
1
2
Number of prior stressors
Trend
Observed
Predicted
Prior Stressor History
0
1
2

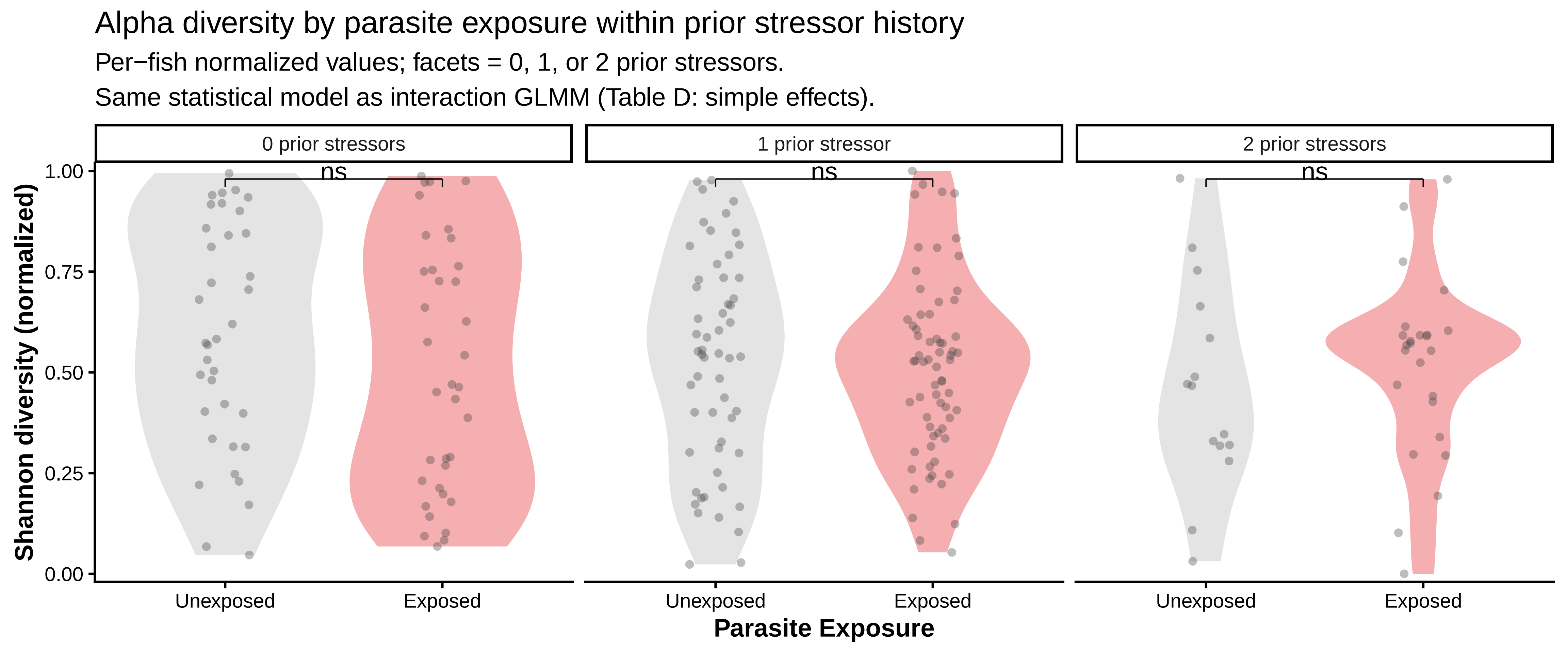
Alpha diversity by parasite exposure within prior stressor history
Per−fish normalized values; facets = 0, 1, or 2 prior stressors.
Same statistical model as interaction GLMM (Table D: simple effects).
0 prior stressors
1 prior stressor
2 prior stressors
ns
ns
ns
Shannon diversity (normalized)
1.00
0.75
0.50
0.25
0.00
Unexposed
Exposed
Unexposed
Exposed
Unexposed
Exposed
Parasite Exposure

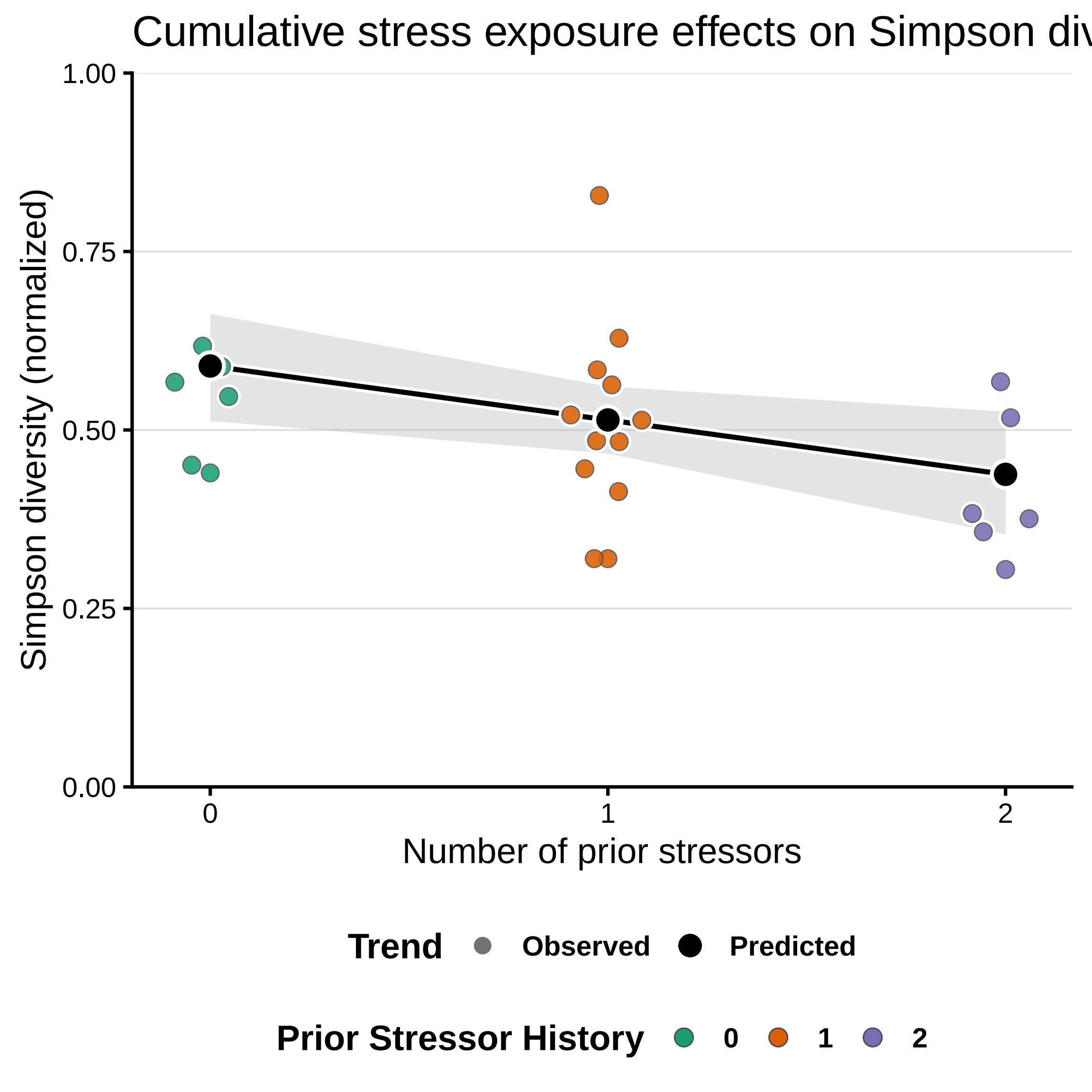
Cumulative stress exposure effects on Simpson div
Simpson diversity (normalized)
1.00
0.75
0.50
0.25
0.00
0
1
2
Number of prior stressors
Trend
Observed
Predicted
Prior Stressor History
0
1
2

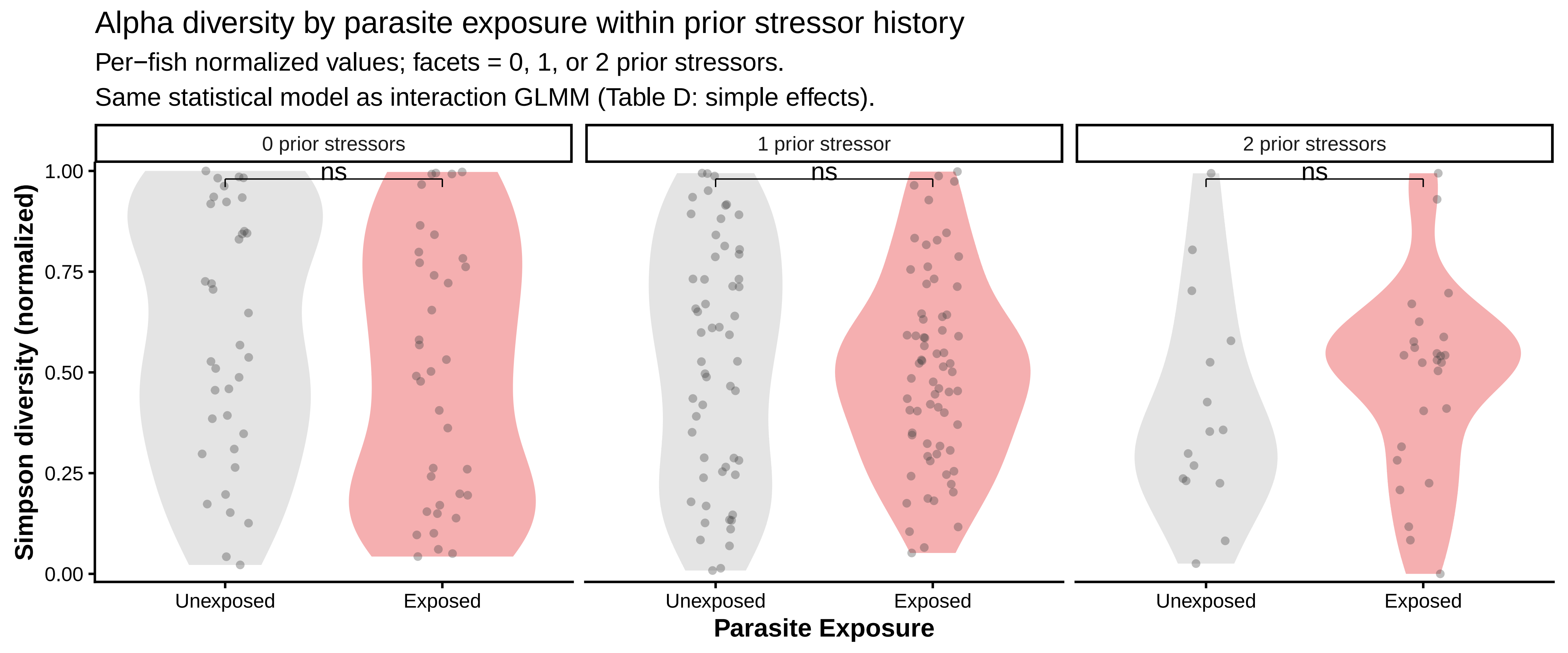
Alpha diversity by parasite exposure within prior stressor history
Per−fish normalized values; facets = 0, 1, or 2 prior stressors.
Same statistical model as interaction GLMM (Table D: simple effects).
0 prior stressors
1 prior stressor
2 prior stressors
ns
ns
ns
Simpson diversity (normalized)
1.00
0.75
0.50
0.25
0.00
Unexposed
Exposed
Parasite Exposure
