## Supplementary Figures for "Historical contingency shapes zebrafish host-microbiome responses to a subsequent biotic challenge": Supplementary_Figures__02__Composition__submission__2026-06-26.pdf

### Supplementary figures (combined)

Build (UTC): 2026−06−26 12:36:44 UTC

Git: 55d6b9a4a7bf134b3db15f0777c389e2da8c31be

Module: 02__Composition

Figures dir: /Sieler2026/Results/02__Composition/Figures

Figures (combined PDF order; p. = first page of that figure in this PDF):

p. 2: Figure S2.2.1 − betadisper_bray_stress_history

p. 3: Figure S2.2.2 − betadisper_bray_stress_history_parasite

p. 4: Figure S2.2.3 − betadisper_canberra_stress_history

p. 5: Figure S2.2.4 − betadisper_canberra_stress_history_parasite

p. 6: Figure S2.2.5 − betadisper_bray_factorial_ATP

p. 7: Figure S2.2.6 − betadisper_canberra_factorial_ATP

p. 8: Figure S2.5.1 − genus_relative_abundance_by_treatment

p. 9: Figure S2.10.1 − pcoa_bray_factorial_ATP

p. 10: Figure S2.11.1 − pcoa_bray_parasite_faceted_by_history

p. 11: Figure S2.12.1 − pcoa_bray_parasite_faceted_by_history_regime_points

p. 12: Figure S2.13.1 − pcoa_bray_stress_history

p. 13: Figure S2.14.1 − pcoa_bray_stress_history_parasite

p. 14: Figure S2.15.1 − pcoa_canberra_factorial_ATP

p. 15: Figure S2.16.1 − pcoa_canberra_parasite_faceted_by_history

p. 16: Figure S2.17.1 − pcoa_canberra_parasite_faceted_by_history_regime_points

p. 17: Figure S2.18.1 − pcoa_canberra_stress_history

p. 18: Figure S2.19.1 − pcoa_canberra_stress_history_parasite

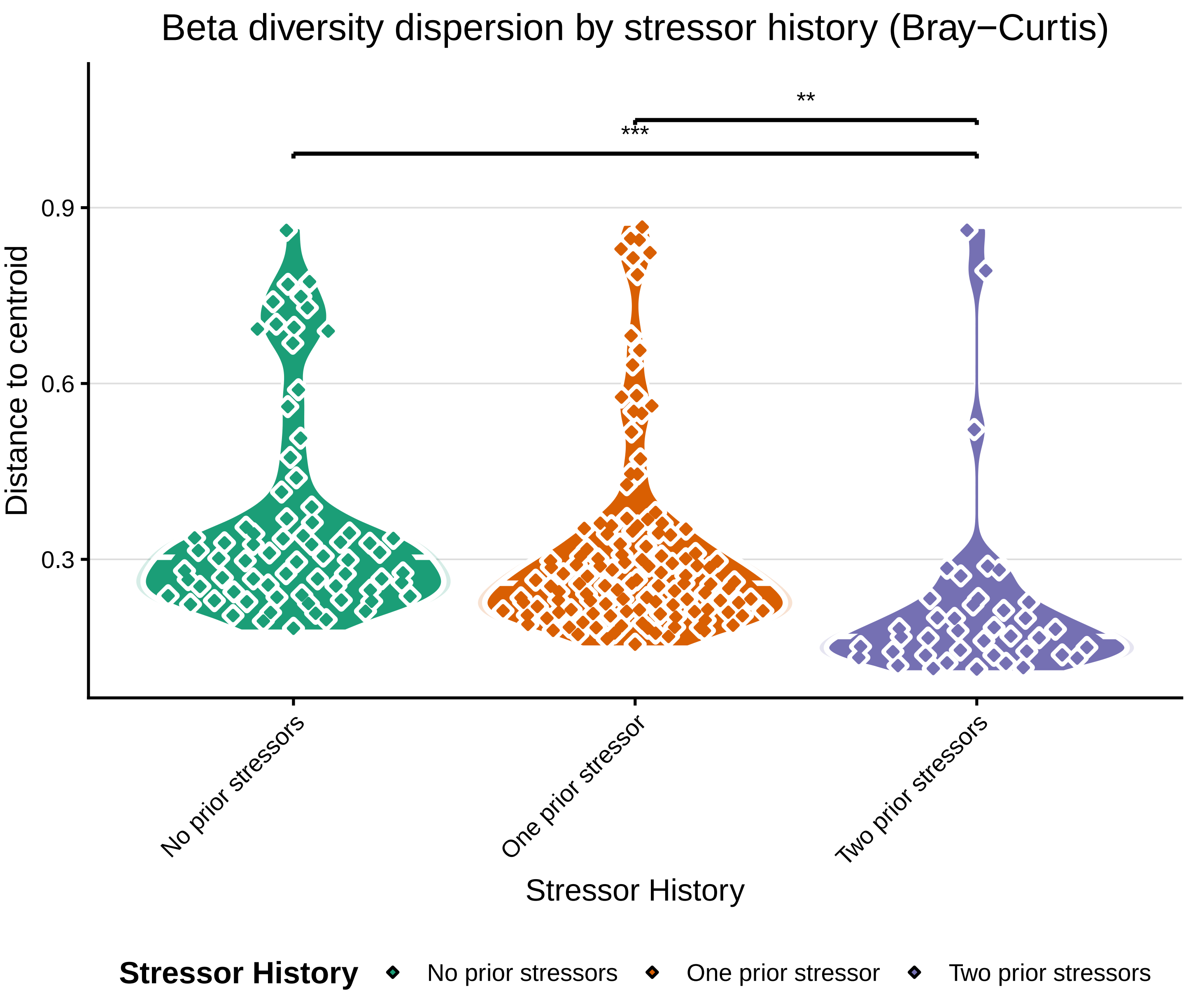

Beta diversity dispersion by stressor history (Bray−Curtis)
**
***
0.9
0.6
0.3
Distance to centroid
No prior stressors
One prior stressor
Two prior stressors
Stressor History
Stressor History
No prior stressors
One prior stressor
Two prior stressors

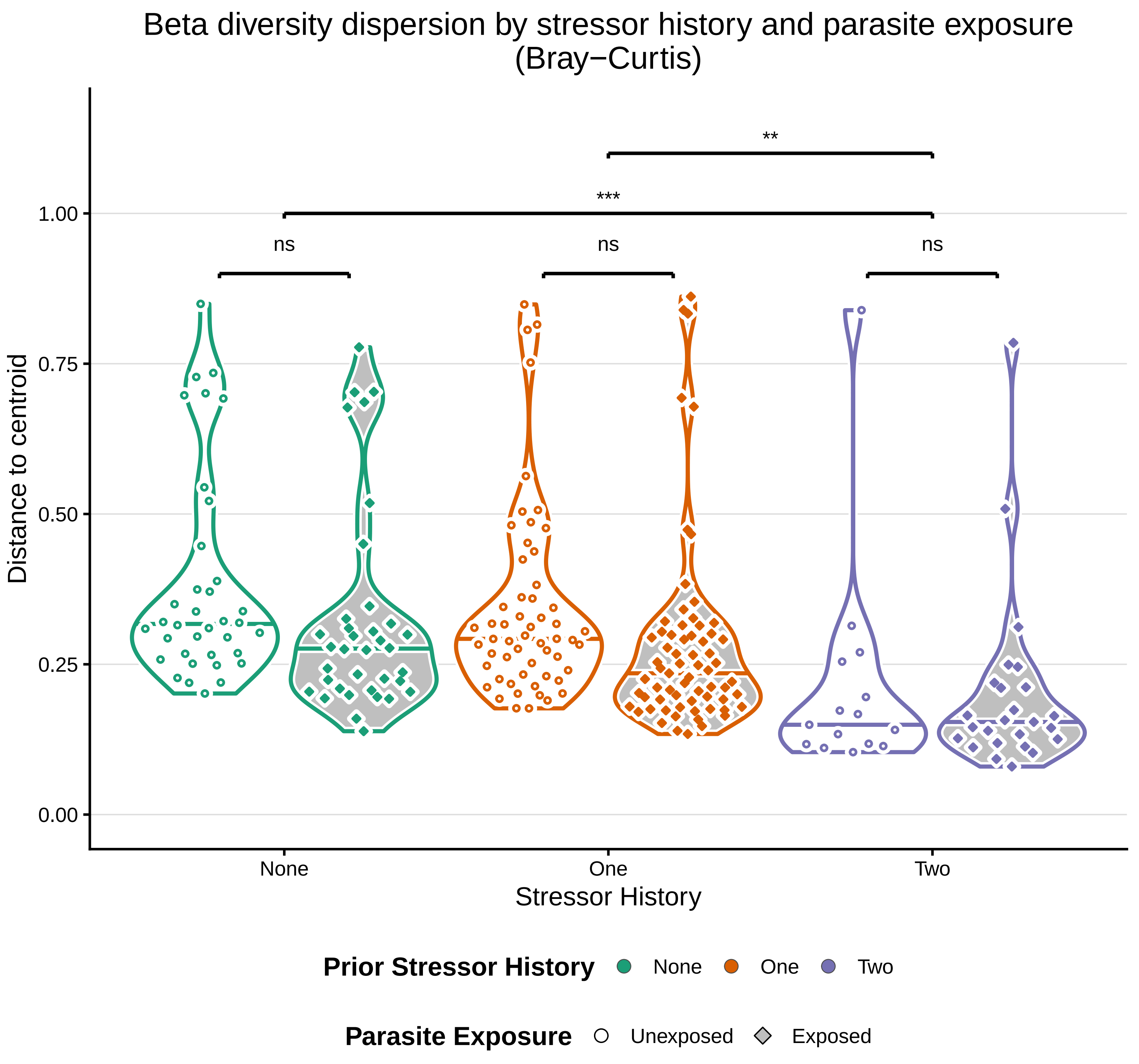

Beta diversity dispersion by stressor history and parasite exposure
(Bray−Curtis)
Distance to centroid
1.00
0.75
0.50
0.25
0.00
**
***
ns
ns
ns
None
One
Two
Stressor History
Prior Stressor History
None
One
Two
Parasite Exposure
Unexposed
Exposed

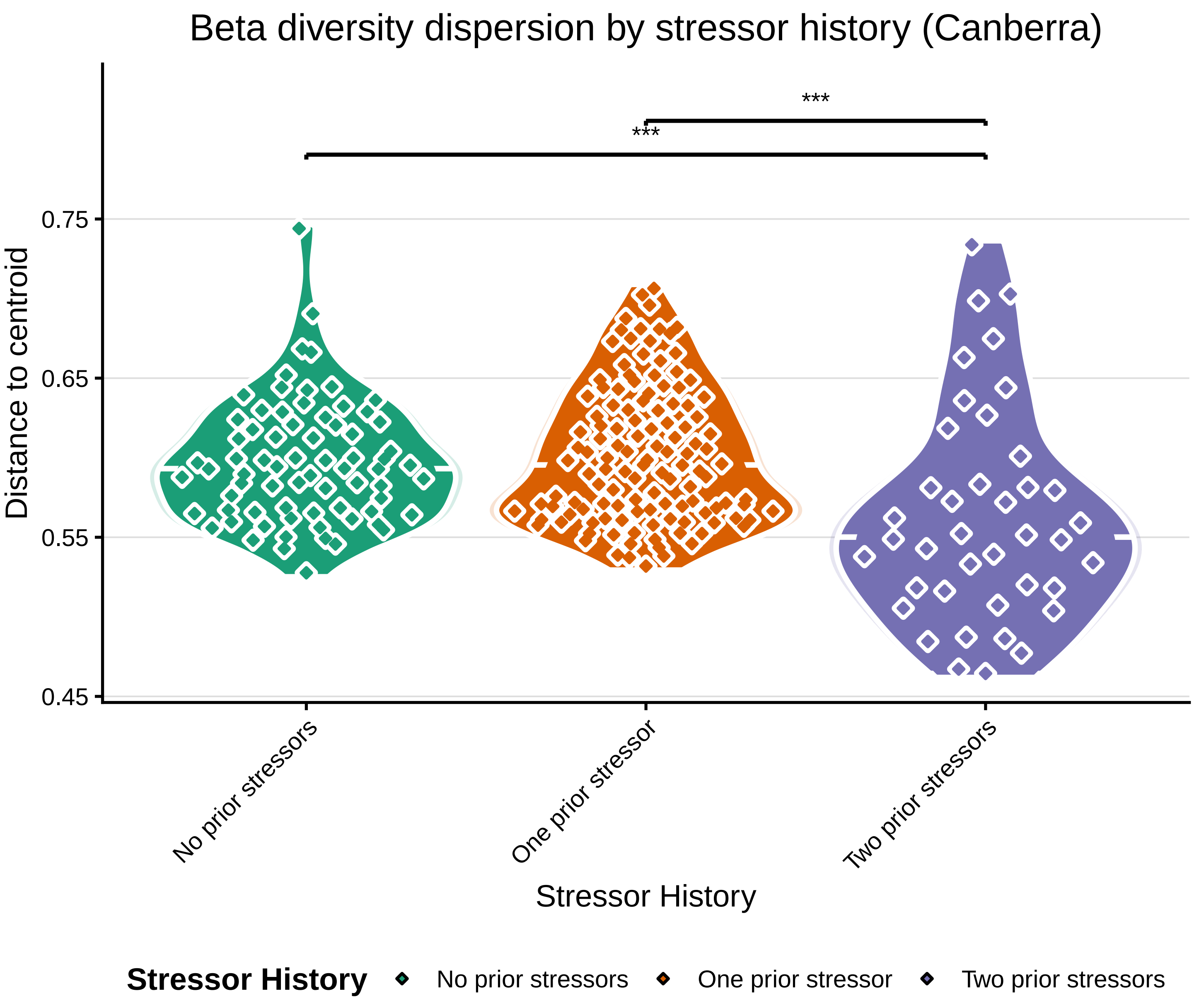

Beta diversity dispersion by stressor history (Canberra)
***
***
Distance to centroid
0.75
0.65
0.55
0.45
No prior stressors
One prior stressor
Two prior stressors
Stressor History
Stressor History
No prior stressors
One prior stressor
Two prior stressors

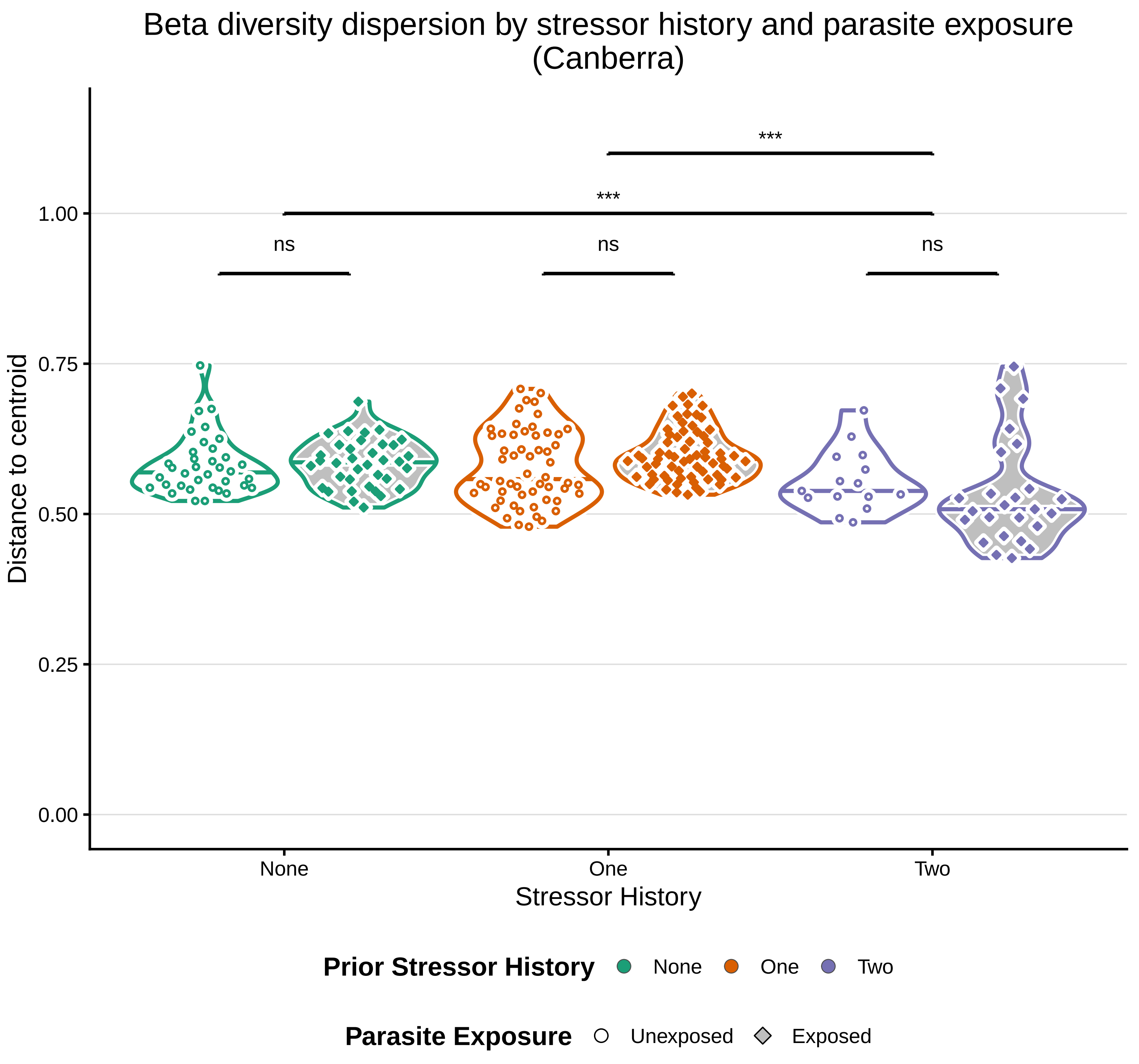

Beta diversity dispersion by stressor history and parasite exposure
(Canberra)
Distance to centroid
1.00
0.75
0.50
0.25
0.00
***
***
ns
ns
ns
None
One
Two
Stressor History
Prior Stressor History
None
One
Two
Parasite Exposure
Unexposed
Exposed

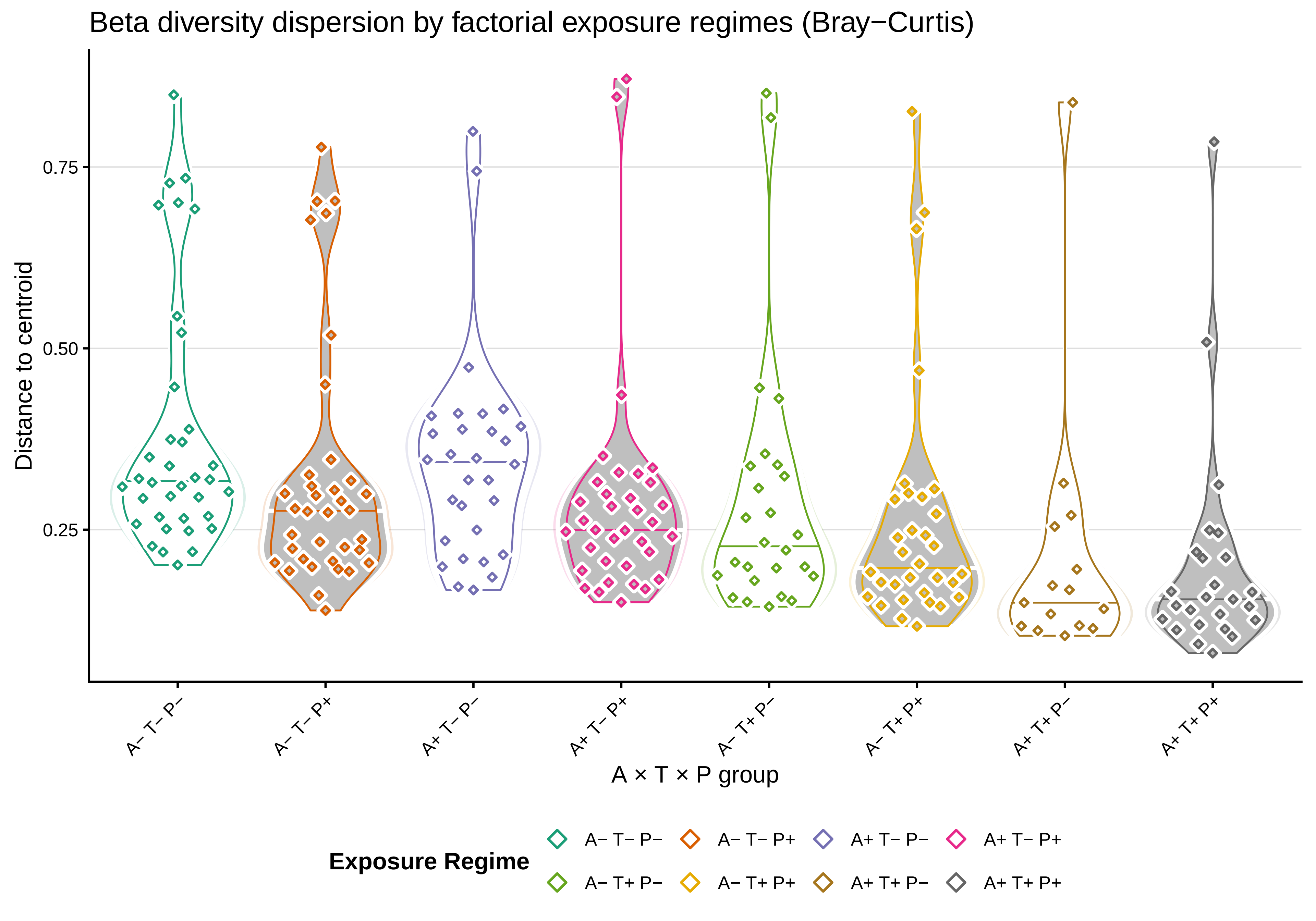
Beta diversity dispersion by factorial exposure regimes (Bray−Curtis)
Distance to centroid
0.75
0.50
0.25
A− T− P−
A− T− P+
A+ T− P−
A+ T− P+
A− T+ P−
A− T+ P+
A+ T+ P−
A+ T+ P+
A × T × P group
Exposure Regime
A− T− P−
A− T− P+
A+ T− P−
A+ T− P+
A− T+ P−
A− T+ P+
A+ T+ P−
A+ T+ P+

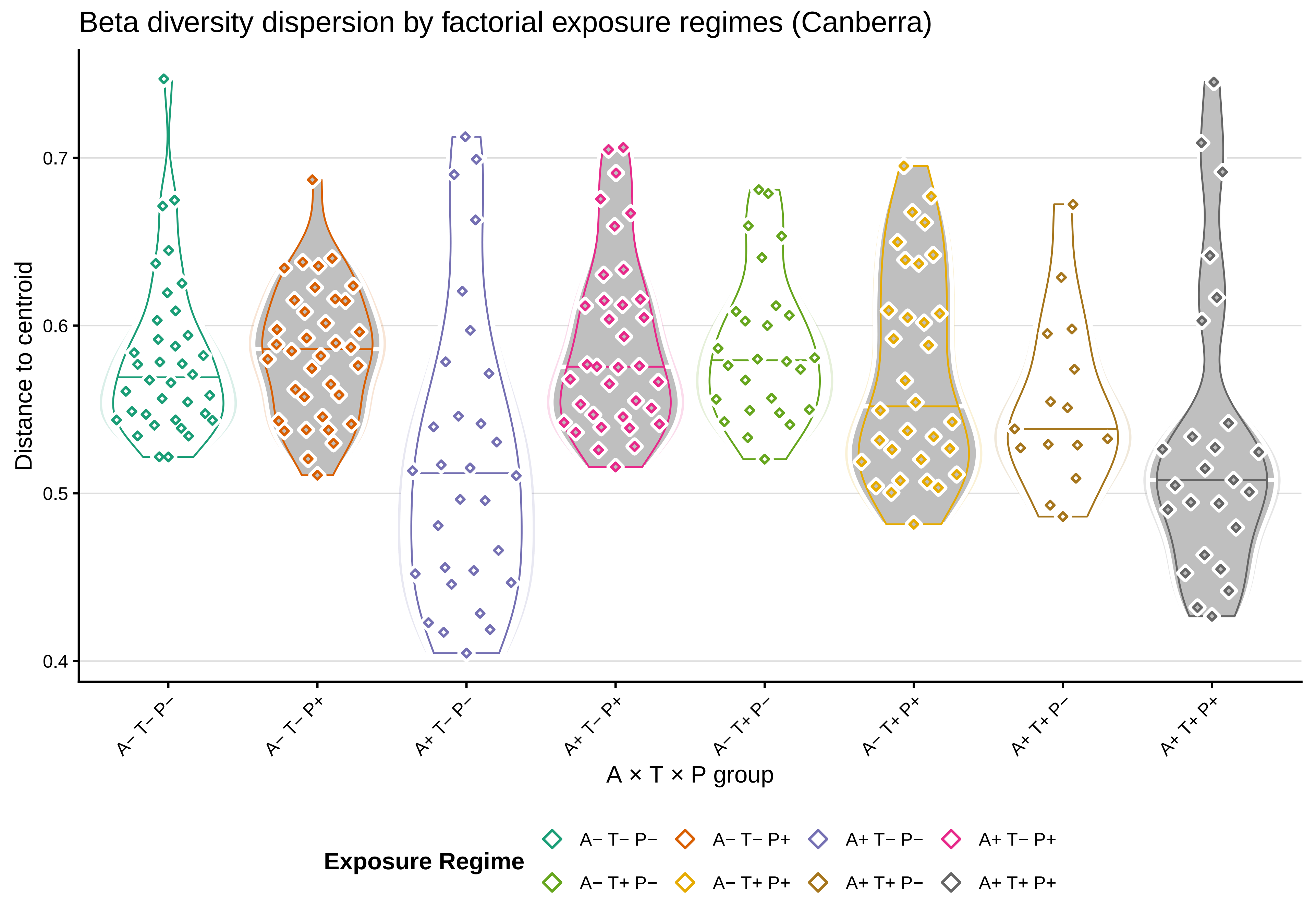

Beta diversity dispersion by factorial exposure regimes (Canberra)
Distance to centroid
0.7
0.6
0.5
0.4
A− T− P−
A− T− P+
A+ T− P−
A+ T− P+
A− T+ P−
A− T+ P+
A+ T+ P−
A+ T+ P+
A × T × P group
Exposure Regime
A− T− P−
A− T− P+
A+ T− P−
A+ T− P+
A− T+ P−
A− T+ P+
A+ T+ P−
A+ T+ P+

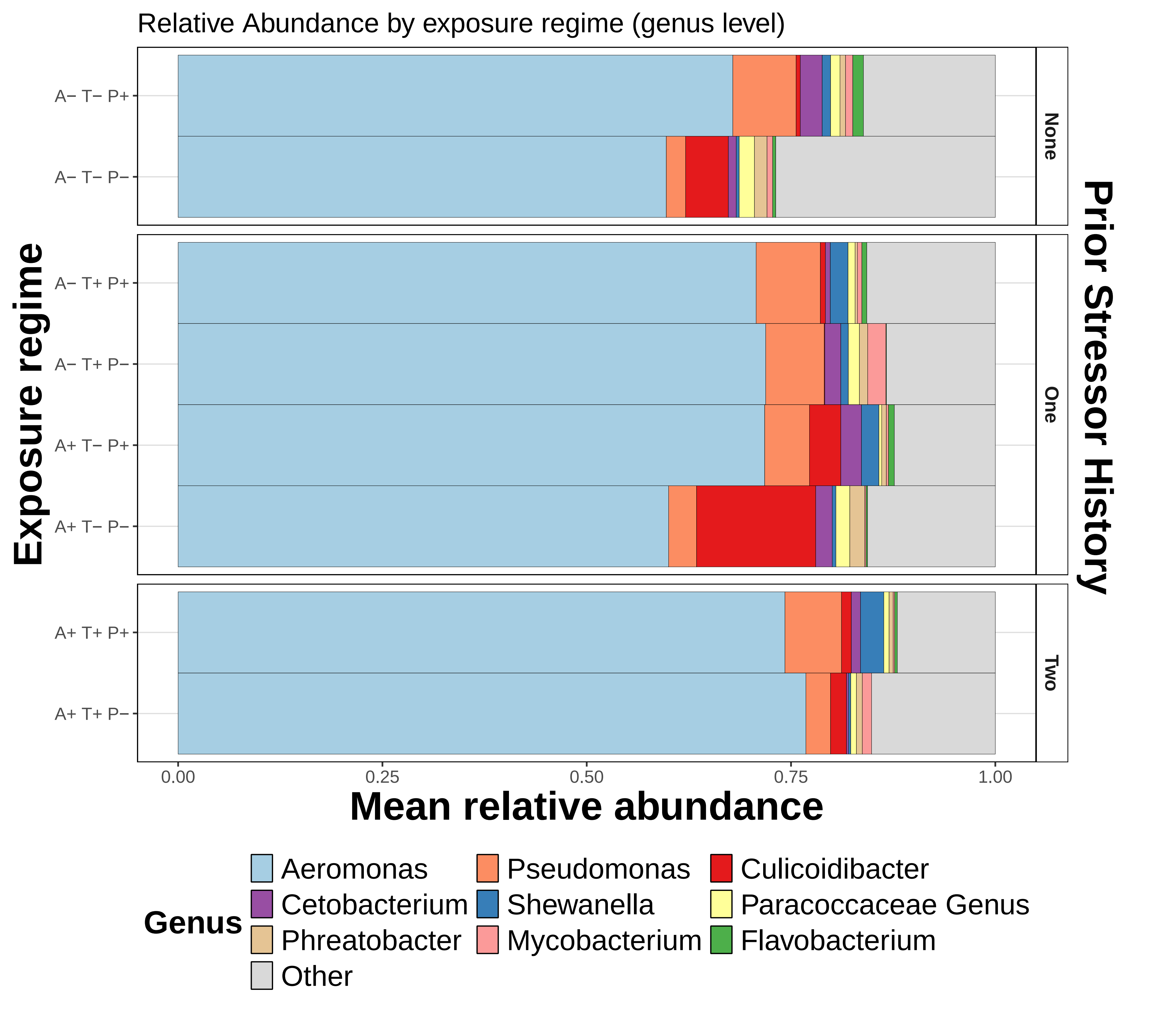
Relative Abundance by exposure regime (genus level)
Exposure regime
A− T− P+
A− T− P−
A− T+ P+
A− T+ P−
A+ T− P+
A+ T− P−
A+ T+ P+
A+ T+ P−
None
One
Two
Prior Stressor History
0.00
0.25
0.50
0.75
1.00
Mean relative abundance
Genus
Aeromonas
Pseudomonas
Culicoidibacter
Cetobacterium
Shewanella
Paracoccaceae Genus
Phreatobacter
Mycobacterium
Flavobacterium
Other

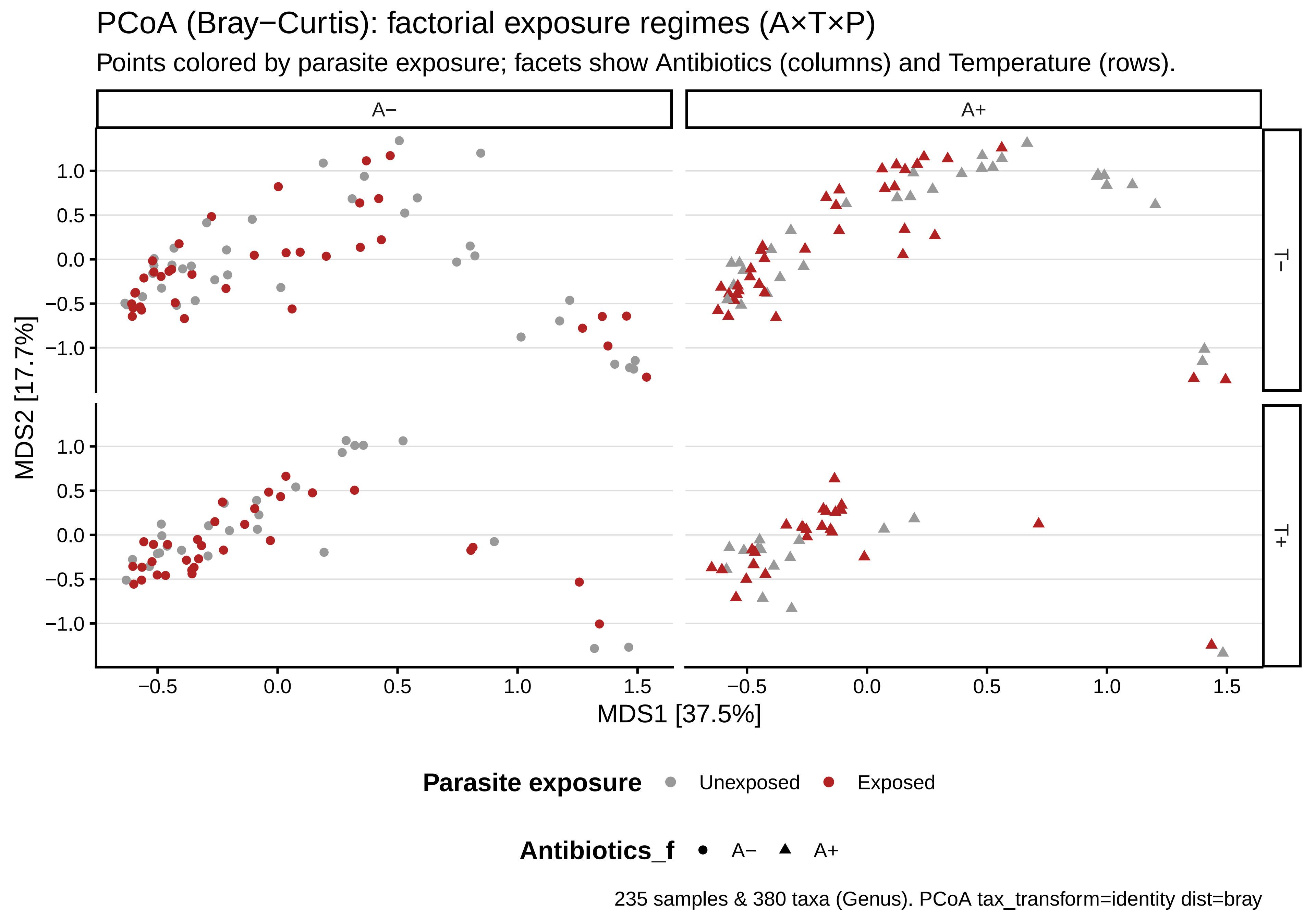
PCoA (Bray−Curtis): factorial exposure regimes (A×T×P)
Points colored by parasite exposure; facets show Antibiotics (columns) and Temperature (rows).
A−
A+
T−
T+
MDS2 [17.7%]
MDS1 [37.5%]
1.0
0.5
0.0
−0.5
−1.0
−0.5
0.0
0.5
1.0
1.5
Parasite exposure
Unexposed
Exposed
Antibiotics_f
A−
A+
235 samples & 380 taxa (Genus). PCoA tax_transform=identity dist=bray

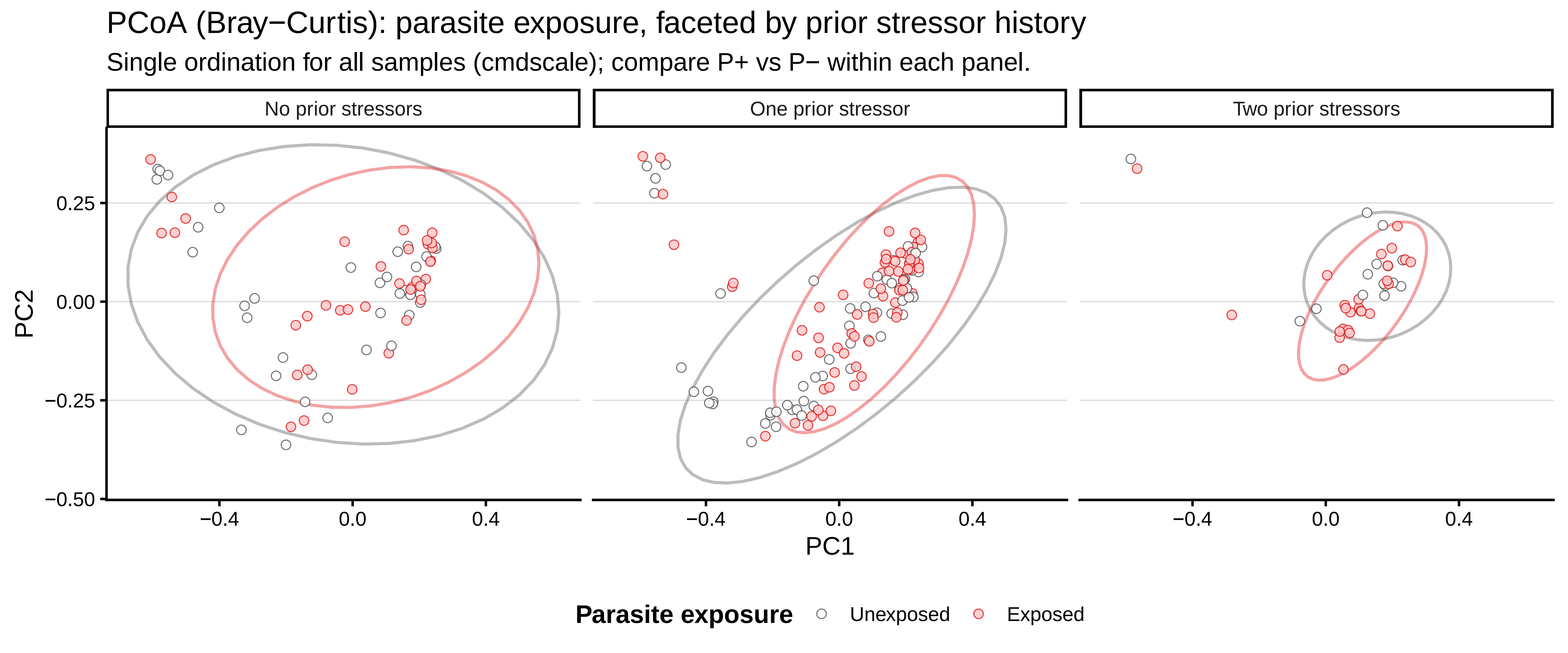

PCoA (Bray−Curtis): parasite exposure, faceted by prior stressor history
Single ordination for all samples (cmdscale); compare P+ vs P− within each panel.
No prior stressors
One prior stressor
Two prior stressors
PC2
0.25
0.00
−0.25
−0.50
−0.4
0.0
0.4
PC1
Parasite exposure
Unexposed
Exposed

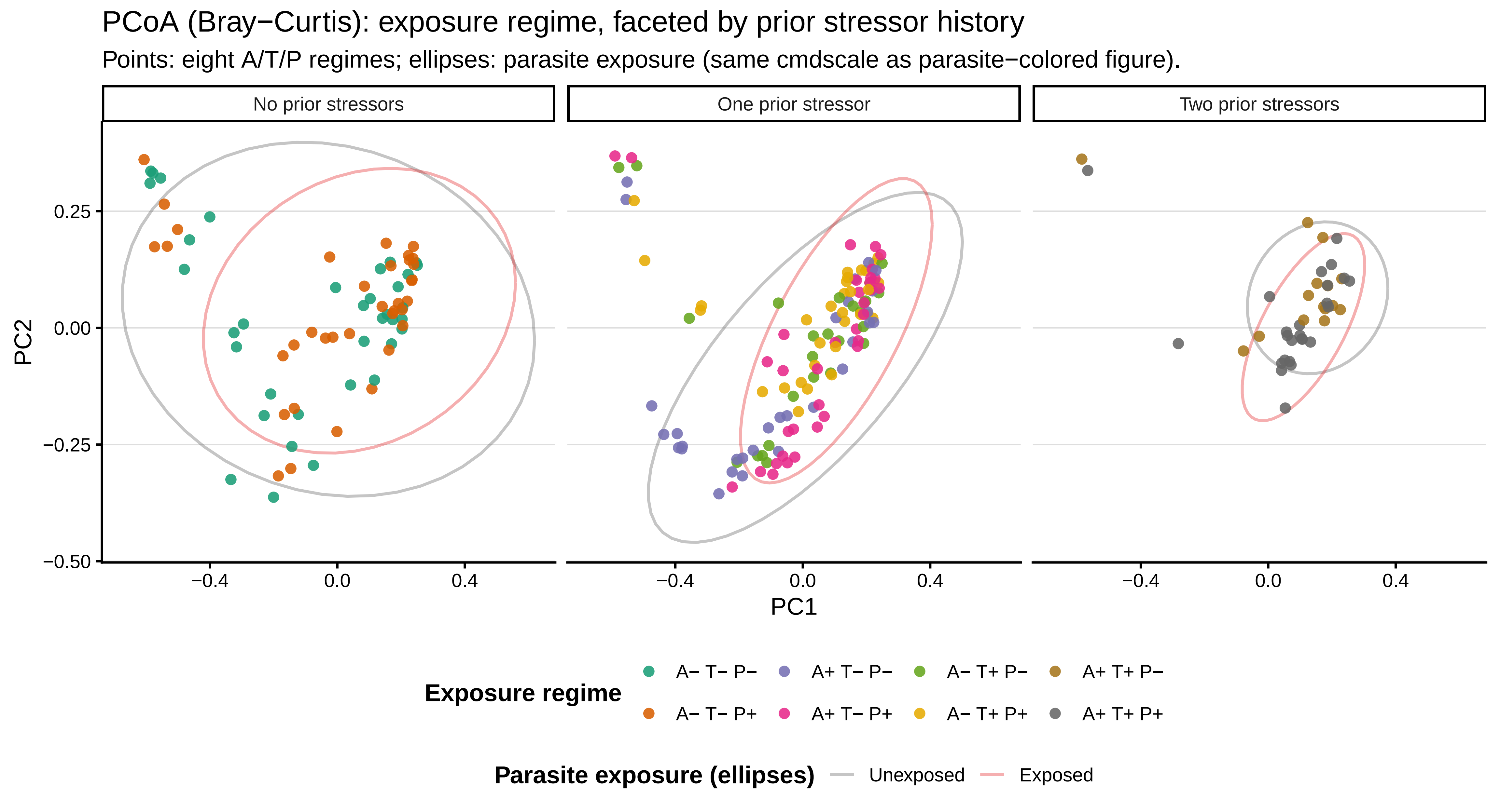
PCoA (Bray−Curtis): exposure regime, faceted by prior stressor history
Points: eight A/T/P regimes; ellipses: parasite exposure (same cmdscale as parasite−colored figure).
No prior stressors
One prior stressor
Two prior stressors
PC2
0.25
0.00
−0.25
−0.50
−0.4
0.0
0.4
PC1
Exposure regime
A− T− P−
A+ T− P−
A− T+ P−
A+ T+ P−
A− T− P+
A+ T− P+
A− T+ P+
A+ T+ P+
Parasite exposure (ellipses)
Unexposed
Exposed

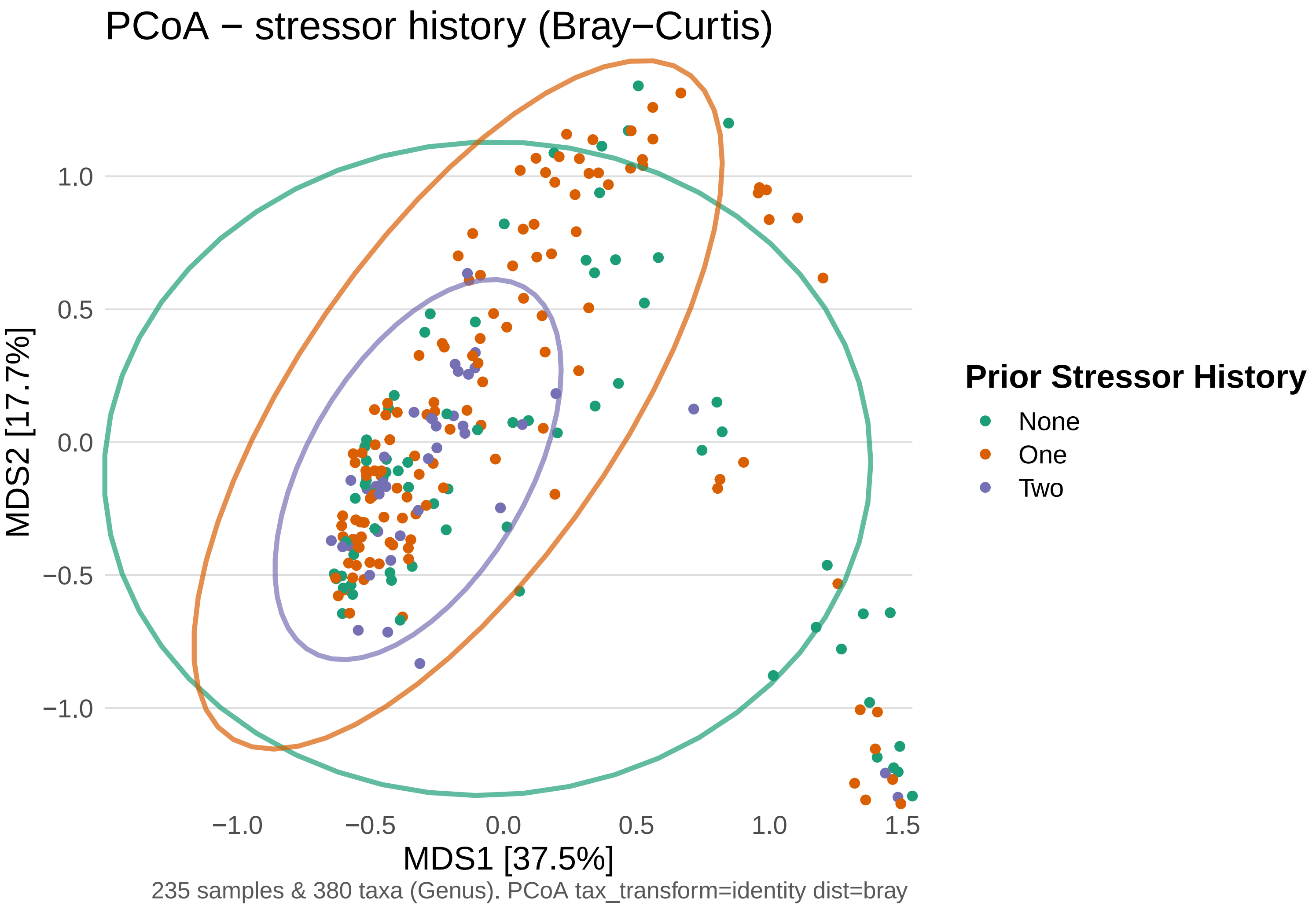

PCoA − stressor history (Bray−Curtis)
MDS2 [17.7%]
1.0
0.5
0.0
−0.5
−1.0
−1.0
−0.5
0.0
0.5
1.0
1.5
MDS1 [37.5%]
235 samples & 380 taxa (Genus). PCoA tax_transform=identity dist=bray
Prior Stressor History
None
One
Two

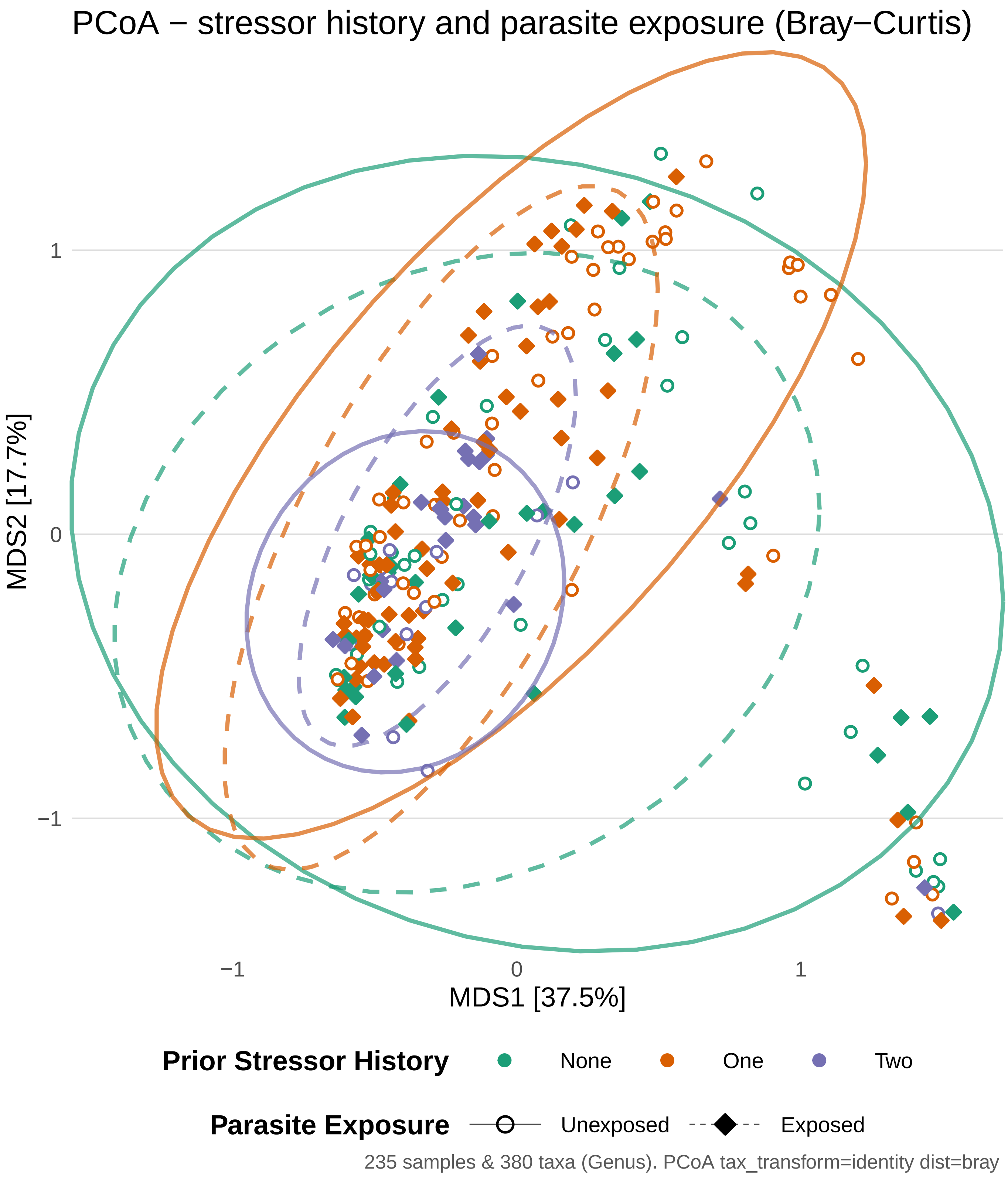
PCoA − stressor history and parasite exposure (Bray−Curtis)
MDS2 [17.7%]
1
0
−1
−1
0
1
MDS1 [37.5%]
Prior Stressor History
None
One
Two
Parasite Exposure
Unexposed
Exposed
235 samples & 380 taxa (Genus). PCoA tax_transform=identity dist=bray

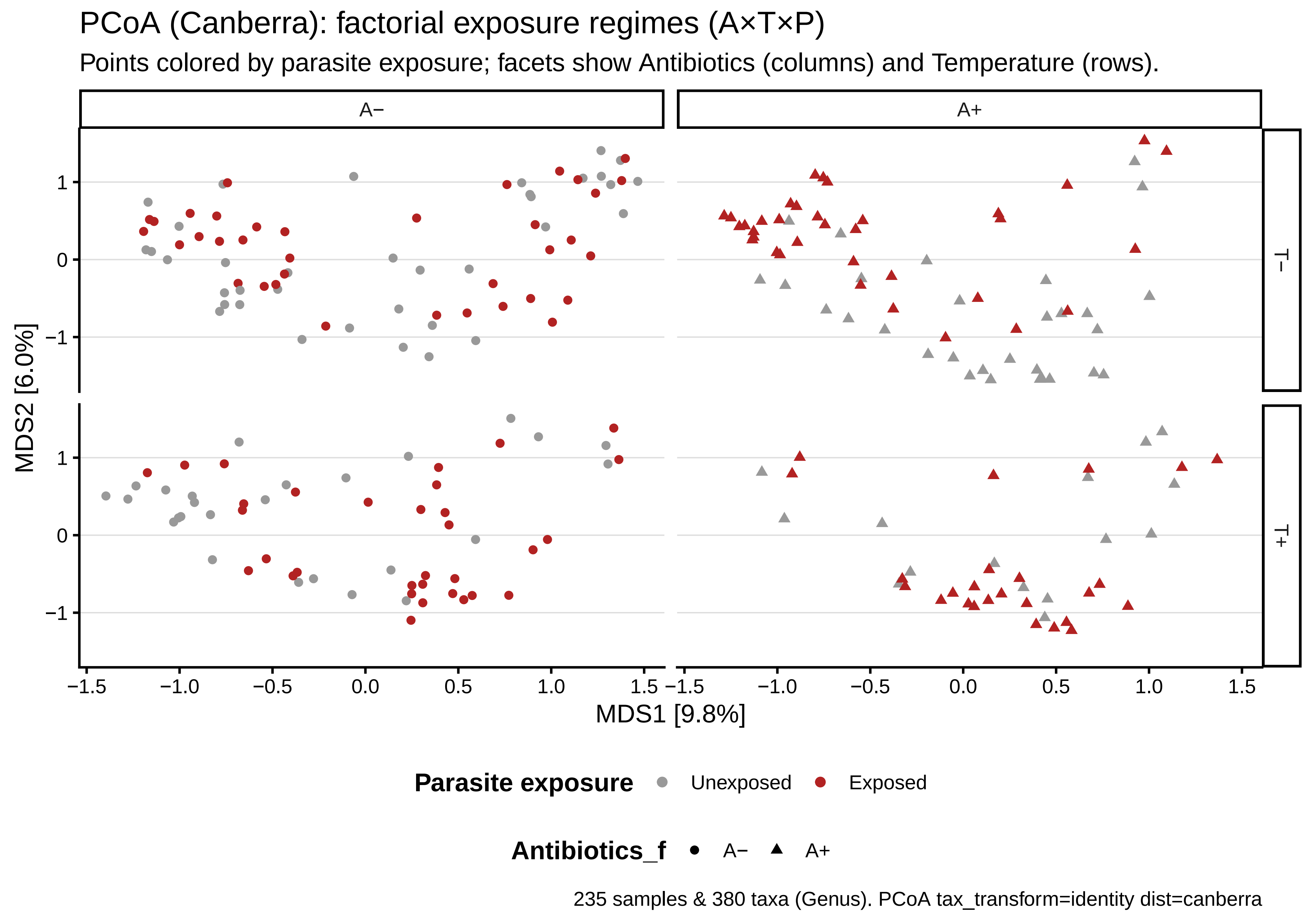

PCoA (Canberra): factorial exposure regimes (A×T×P)
Points colored by parasite exposure; facets show Antibiotics (columns) and Temperature (rows).
A−
A+
T−
T+
MDS2 [6.0%]
MDS1 [9.8%]
1
0
−1
−1.5
−1.0
−0.5
0.0
0.5
1.0
1.5
Parasite exposure
Unexposed
Exposed
Antibiotics_f
A−
A+
235 samples & 380 taxa (Genus). PCoA tax_transform=identity dist=canberra

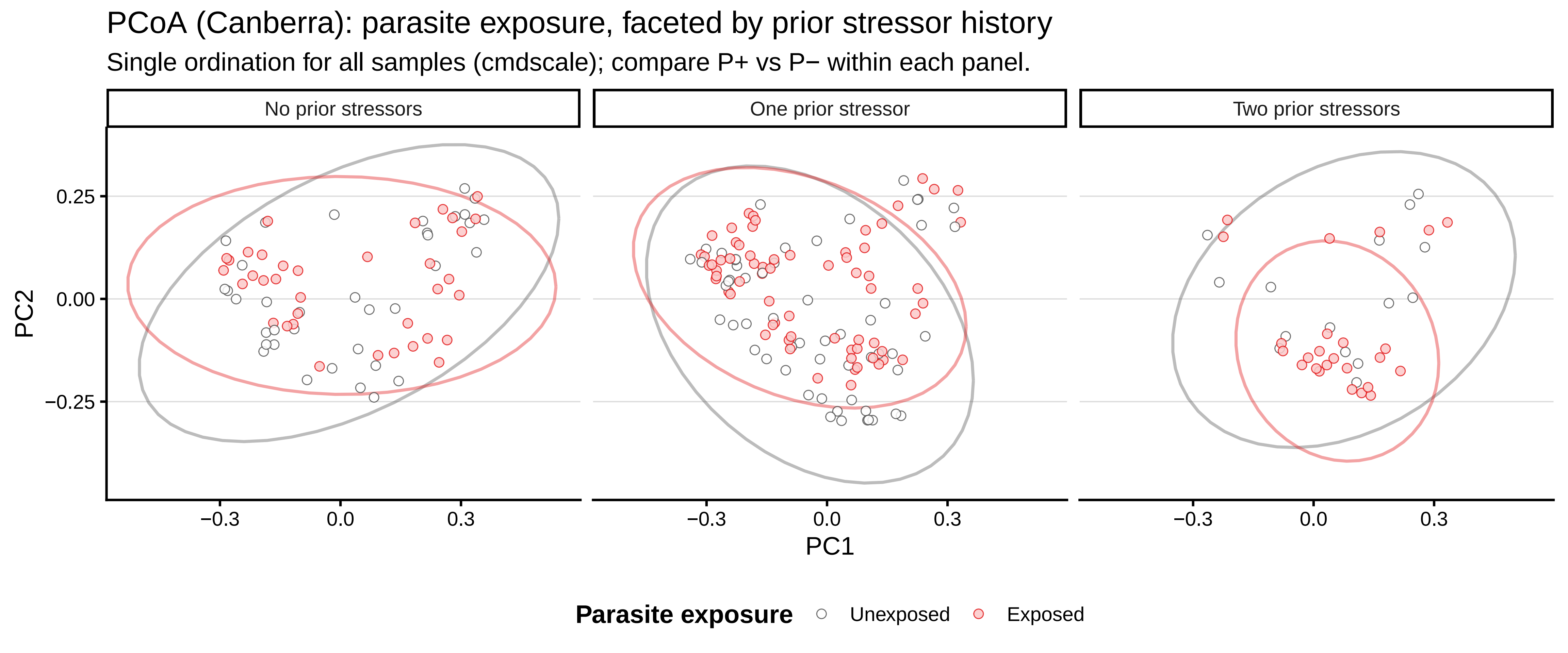
PCoA (Canberra): parasite exposure, faceted by prior stressor history
Single ordination for all samples (cmdscale); compare P+ vs P− within each panel.
No prior stressors
One prior stressor
Two prior stressors
PC2
0.25
0.00
−0.25
−0.3
0.0
0.3
PC1
Parasite exposure
Unexposed
Exposed

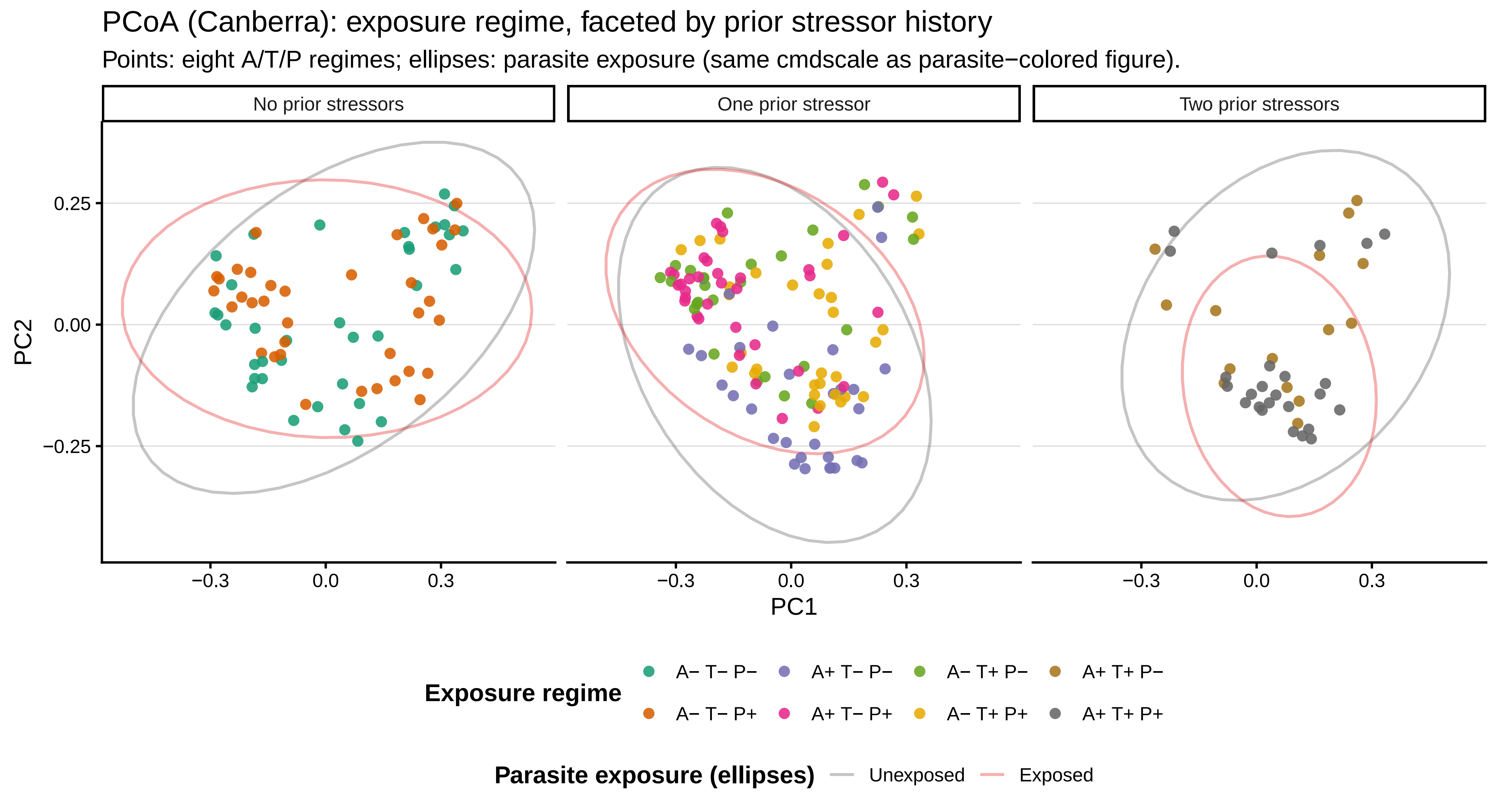
PCoA (Canberra): exposure regime, faceted by prior stressor history
Points: eight A/T/P regimes; ellipses: parasite exposure (same cmdscale as parasite−colored figure).
No prior stressors
One prior stressor
Two prior stressors
0.25
0.00
−0.25
PC2
−0.3
0.0
0.3
PC1
Exposure regime
A− T− P−
A+ T− P−
A− T+ P−
A+ T+ P−
A− T− P+
A+ T− P+
A− T+ P+
A+ T+ P+
Parasite exposure (ellipses)
Unexposed
Exposed

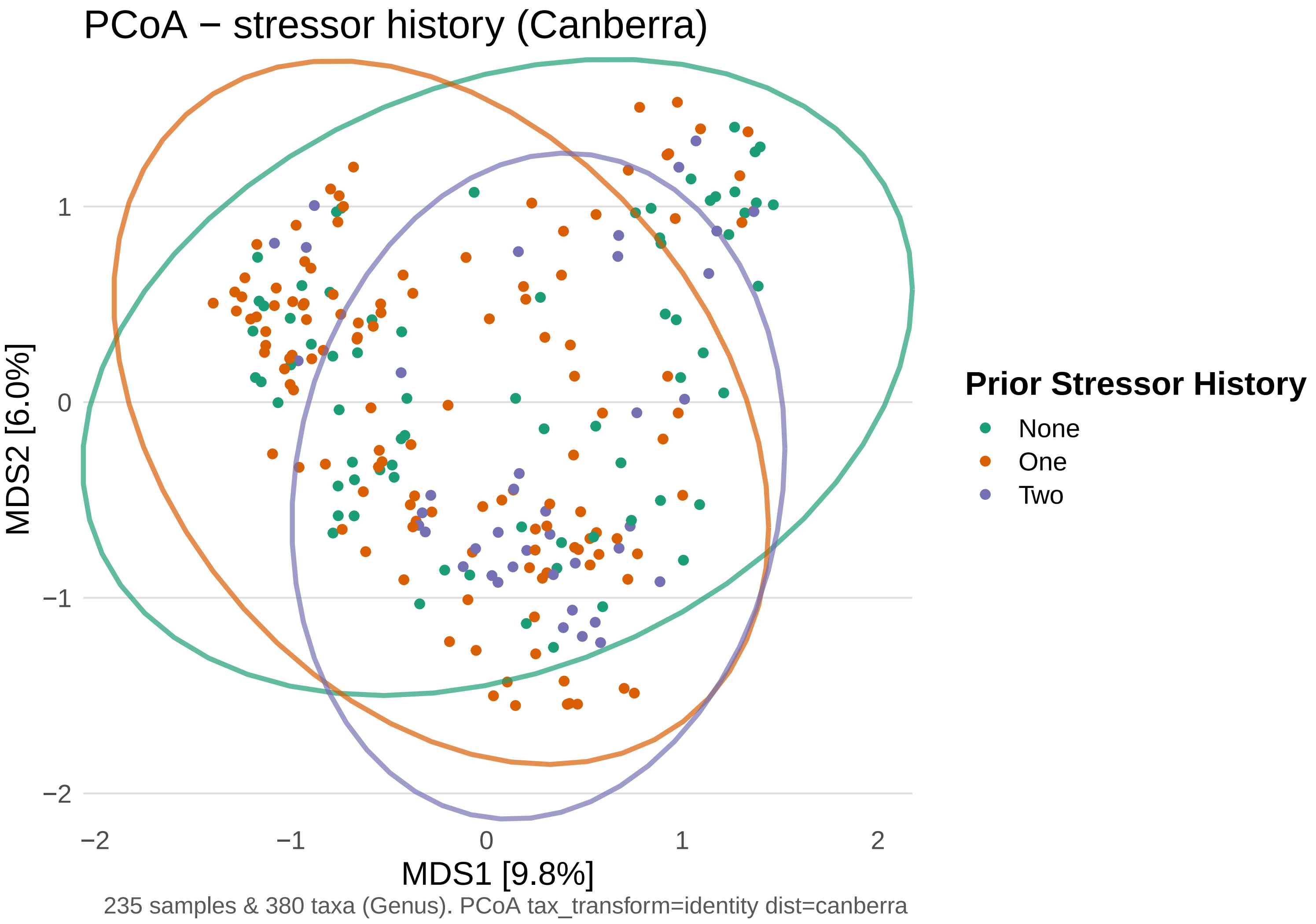
PCoA − stressor history (Canberra)
MDS2 [6.0%]
MDS1 [9.8%]
1
0
−1
−2
−2
−1
0
1
2
Prior Stressor History
None
One
Two
235 samples & 380 taxa (Genus). PCoA tax_transform=identity dist=canberra

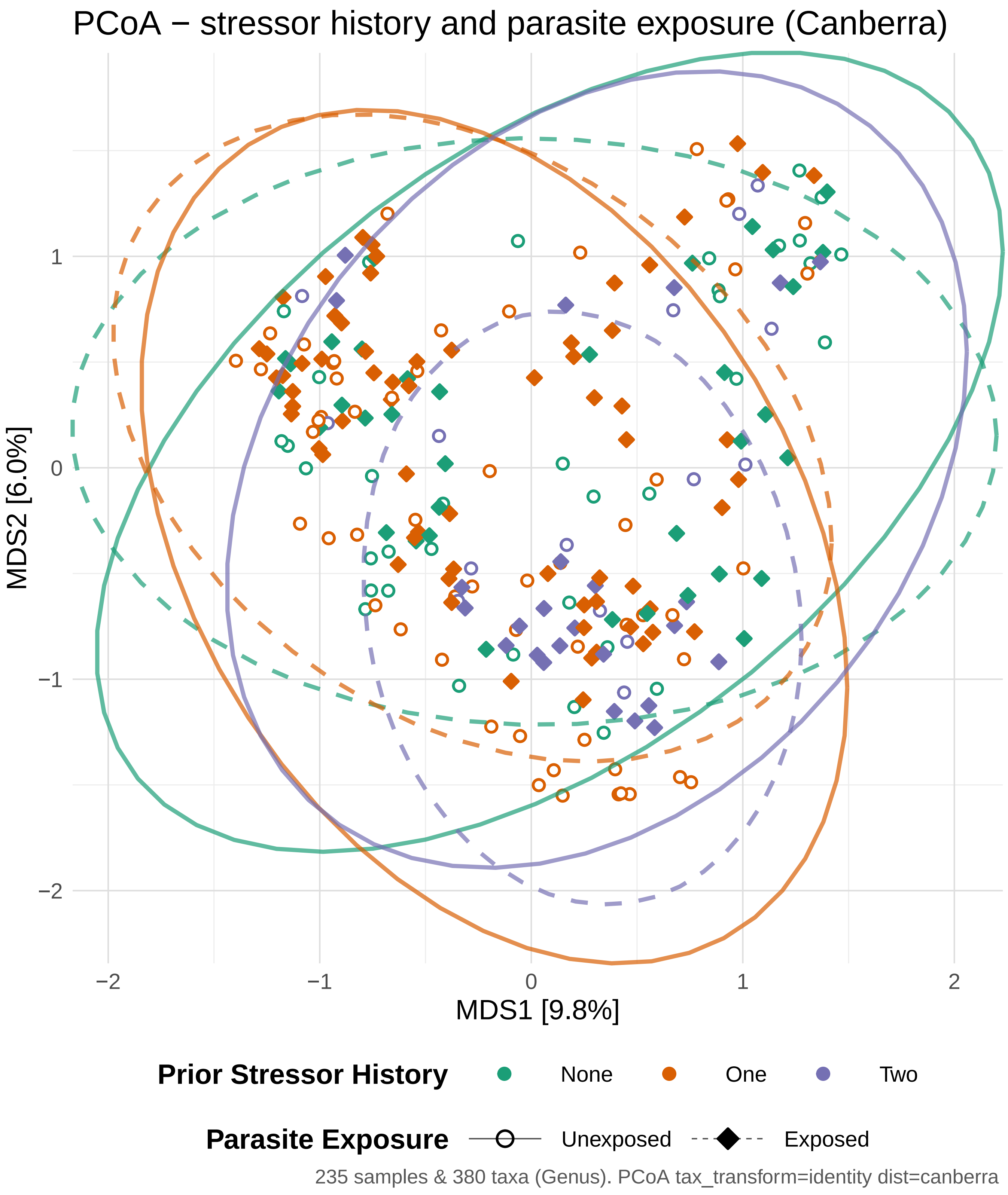

PCoA − stressor history and parasite exposure (Canberra)
MDS2 [6.0%]
MDS1 [9.8%]
1
0
−1
−2
−2
−1
0
1
2
Prior Stressor History
None
One
Two
Parasite Exposure
Unexposed
Exposed
235 samples & 380 taxa (Genus). PCoA tax_transform=identity dist=canberra
