## Supplementary Figures for "Historical contingency shapes zebrafish host-microbiome responses to a subsequent biotic challenge": Supplementary_Figures__03__DiffAbund__submission__2026-06-26.pdf

### Supplementary figures (combined)

Build (UTC): 2026−06−26 12:36:45 UTC

Git: 55d6b9a4a7bf134b3db15f0777c389e2da8c31be

Module: 03__DiffAbund

Figures dir: /Sieler2026/Results/03__DiffAbund/Figures

Figures (combined PDF order; p. = first page of that figure in this PDF):

p. 2: Figure S3.10.1 − maaslin_mortality_top_taxa_coef_bar__Tank

p. 3: Figure S3.11.1 − maaslin_mortality_top_taxa_scatter_combined__Tank

p. 4: Figure S3.12.1 − maaslin_mortality_top_taxa_scatter_facets__Tank

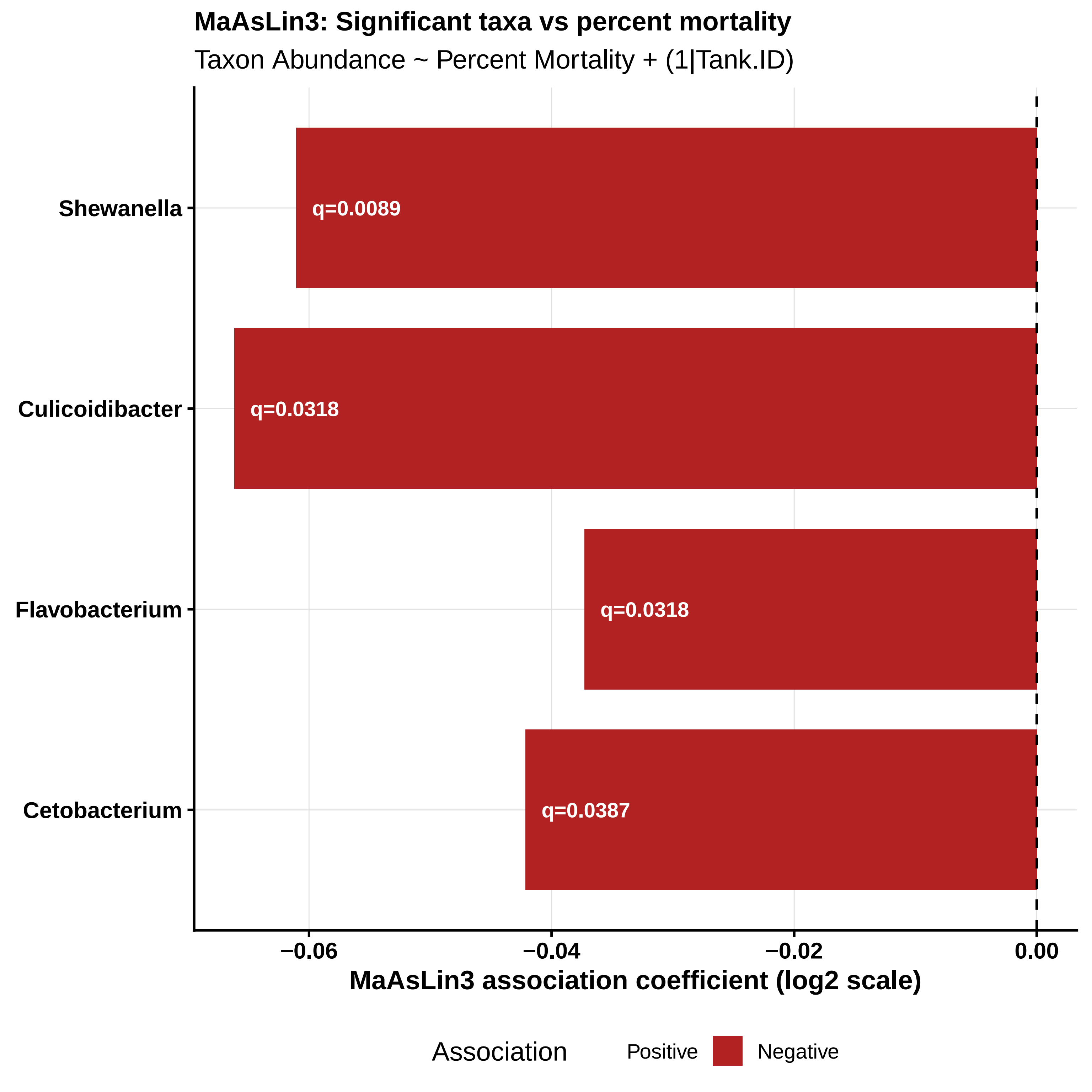
MaAsLin3: Significant taxa vs percent mortality
Taxon Abundance ~ Percent Mortality + (1|Tank.ID)
Shewanella
q=0.0089
Culicoidibacter
q=0.0318
Flavobacterium
q=0.0318
Cetobacterium
q=0.0387
−0.06
−0.04
−0.02
0.00
MaAsLin3 association coefficient (log2 scale)
Association
Positive
Negative

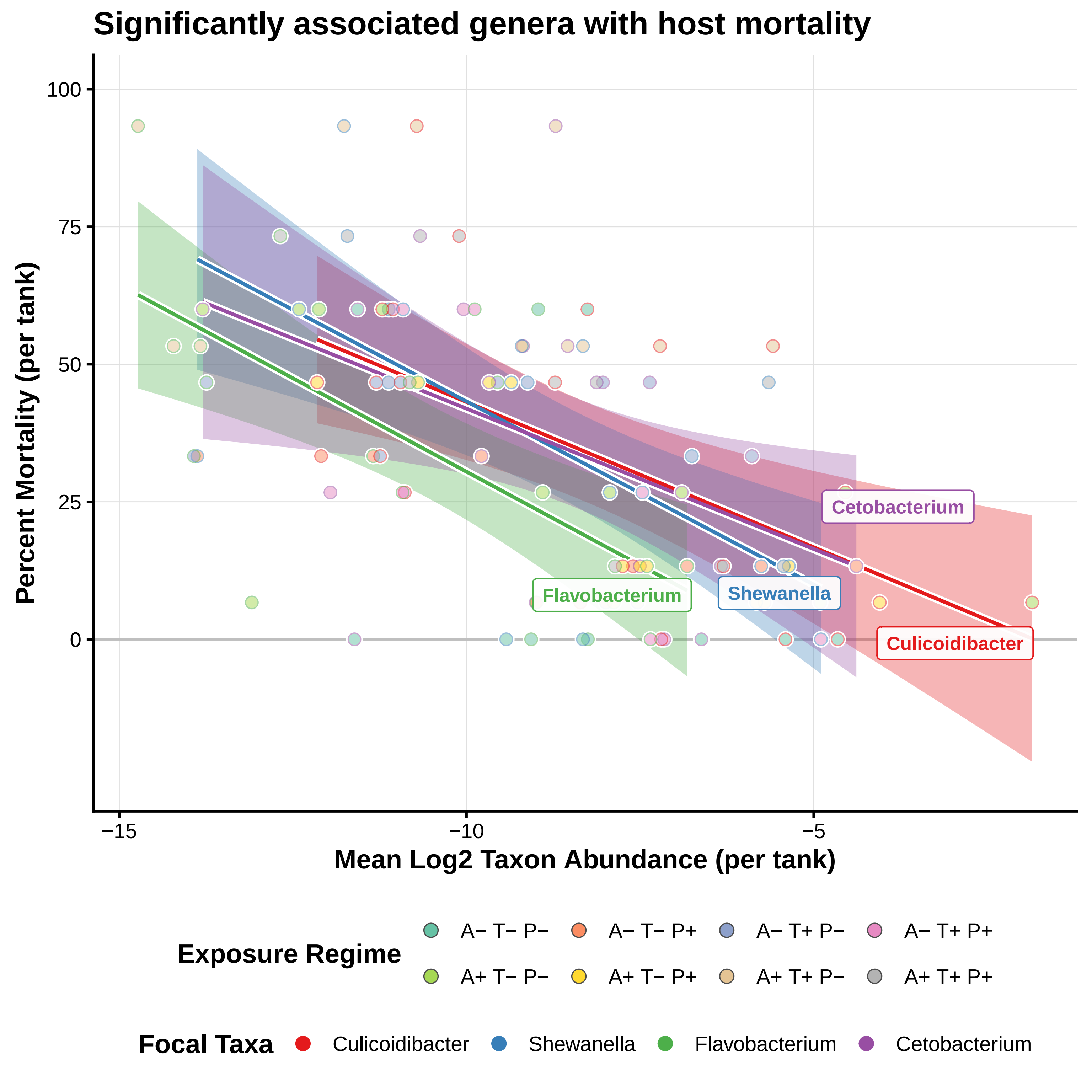

Significantly associated genera with host mortality
Percent Mortality (per tank)
100
75
50
25
0
−15
−10
−5
Mean Log2 Taxon Abundance (per tank)
Cetobacterium
Flavobacterium
Shewanella
Culicoidibacter
Exposure Regime
A− T− P−
A− T− P+
A− T+ P−
A− T+ P+
A+ T− P−
A+ T− P+
A+ T+ P−
A+ T+ P+
Focal Taxa
Culicoidibacter
Shewanella
Flavobacterium
Cetobacterium

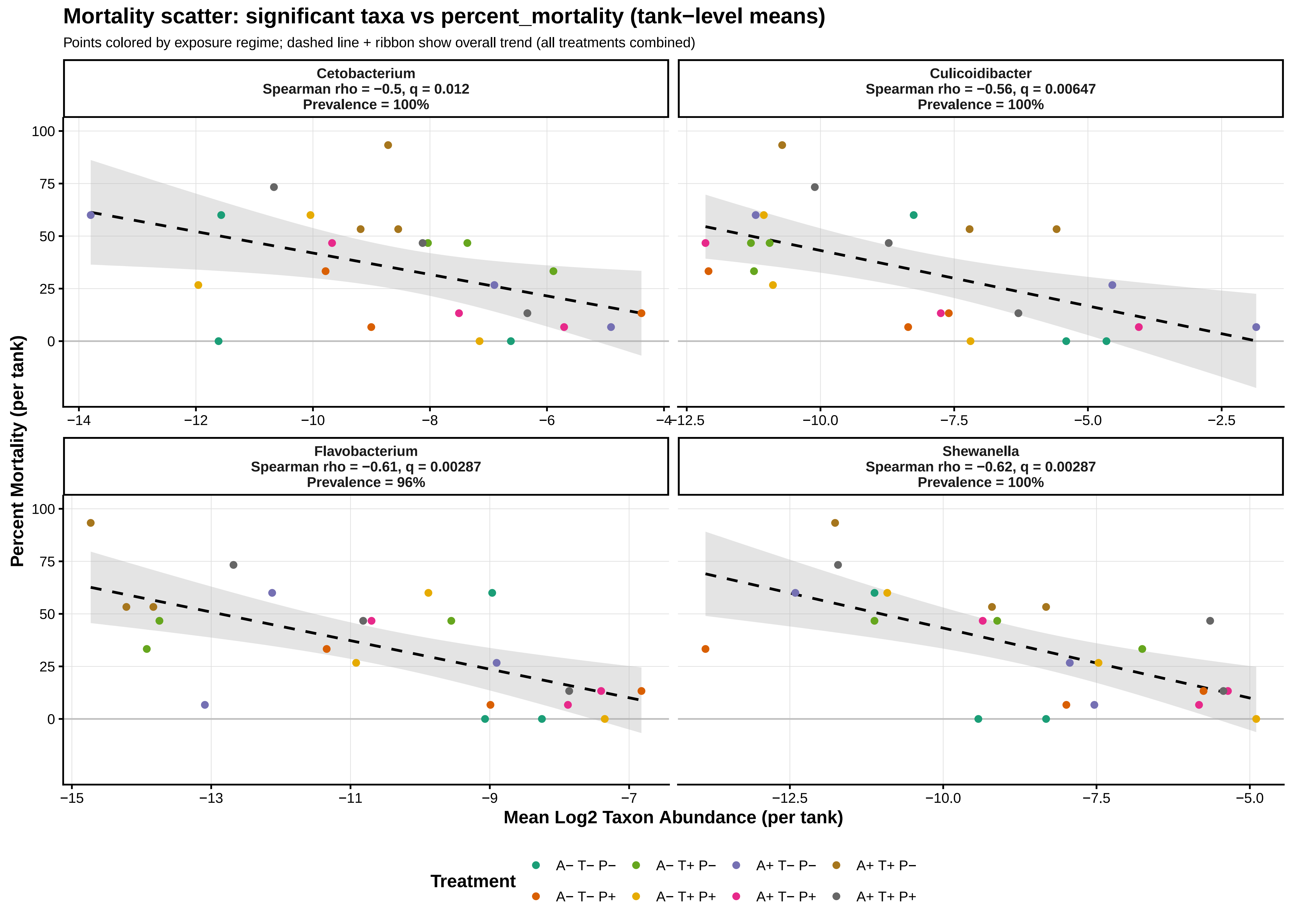

Mortality scatter: significant taxa vs percent_mortality (tank−level means)
Points colored by exposure regime; dashed line + ribbon show overall trend (all treatments combined)
Cetobacterium
Spearman rho = −0.5, q = 0.012
Prevalence = 100%
Culicoidibacter
Spearman rho = −0.56, q = 0.00647
Prevalence = 100%
Flavobacterium
Spearman rho = −0.61, q = 0.00287
Prevalence = 96%
Shewanella
Spearman rho = −0.62, q = 0.00287
Prevalence = 100%
Percent Mortality (per tank)
Mean Log2 Taxon Abundance (per tank)
Treatment
A− T− P−
A− T+ P−
A+ T− P−
A+ T+ P−
A− T− P+
A− T+ P+
A+ T− P+
A+ T+ P+
