## Supplementary Figures for "Historical contingency shapes zebrafish host-microbiome responses to a subsequent biotic challenge": Supplementary_Figures__04__DiffGeneExp__submission__2026-06-26.pdf

### Supplementary figures (combined)

Build (UTC): 2026−06−26 12:36:45 UTC

Git: 55d6b9a4a7bf134b3db15f0777c389e2da8c31be

Module: 04__DiffGeneExp

Figures dir: /Sieler2026/Results/04__DiffGeneExp/Figures

Figures (combined PDF order; p. = first page of that figure in this PDF):

p. 2: Figure S4.4.1 − heatmap_top_50_genes_parasite_exposure

p. 3: Figure S4.3.1 − significant_genes_by_treatment_bar

#### Top 50 Differentially Expressed Genes: Parasite Exposure by Stressor History

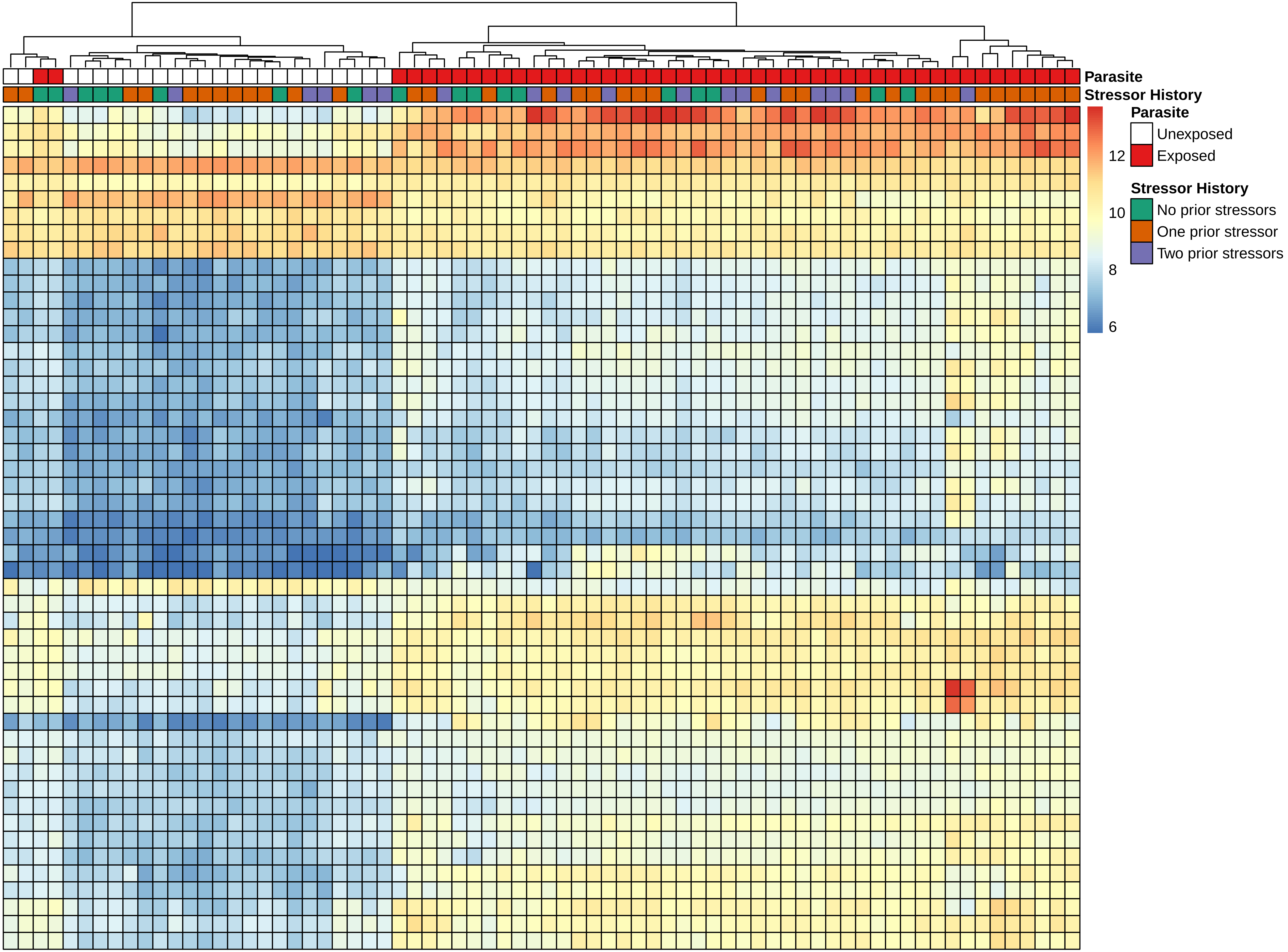

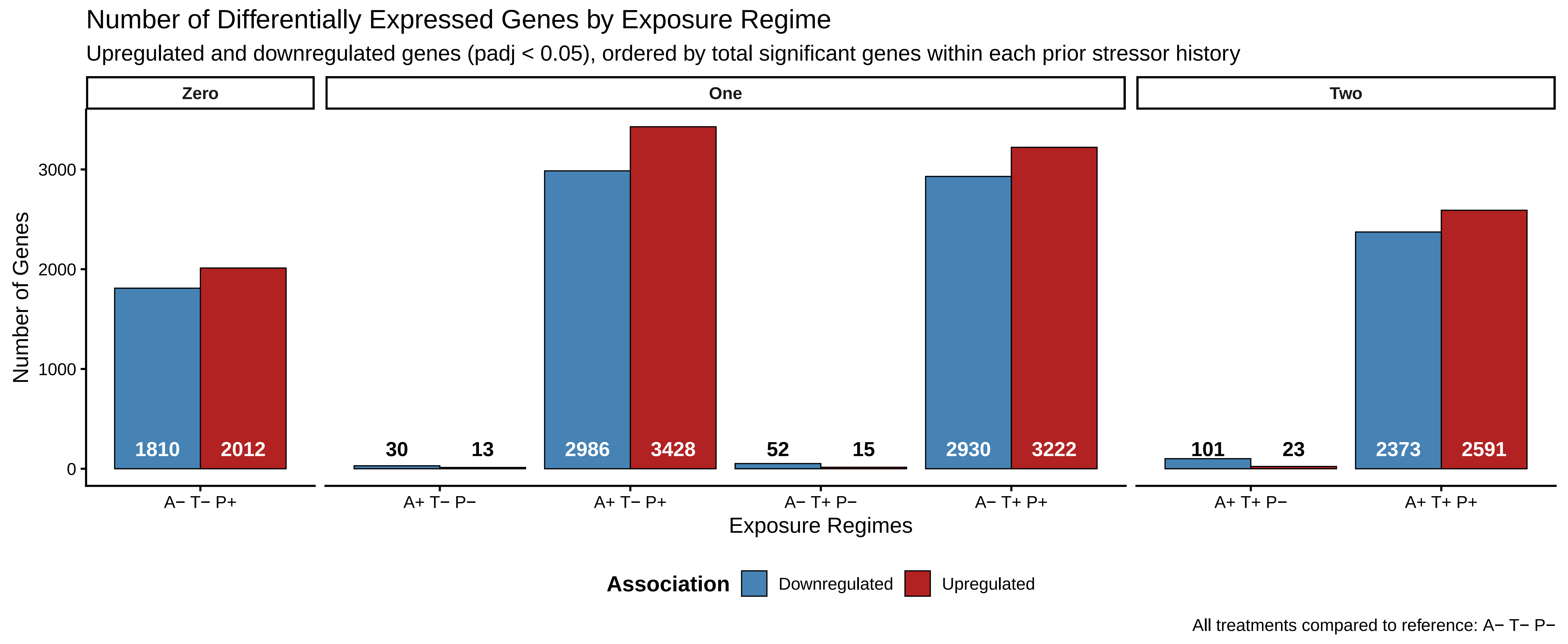

Number of Differentially Expressed Genes by Exposure Regime
Upregulated and downregulated genes (padj < 0.05), ordered by total significant genes within each prior stressor history
Zero
One
Two
Number of Genes
0
1000
2000
3000
1810
2012
30
13
2986
3428
52
15
2930
3222
101
23
2373
2591
A− T− P+
A+ T− P−
A+ T− P+
A− T+ P−
A− T+ P+
A+ T+ P−
A+ T+ P+
Exposure Regimes
Association
Downregulated
Upregulated
All treatments compared to reference: A− T− P−
