## Supplementary Figures for "Historical contingency shapes zebrafish host-microbiome responses to a subsequent biotic challenge": Supplementary_Figures__05__Mort-Inf__submission__2026-06-26.pdf

### Supplementary figures (combined)

Build (UTC): 2026−06−26 12:36:46 UTC

Git: 55d6b9a4a7bf134b3db15f0777c389e2da8c31be

Module: 05__Mort−Inf

Figures dir: /Sieler2026/Results/05__Mort−Inf/Figures

Figures (combined PDF order; p. = first page of that figure in this PDF):

p. 2: Figure S5.2.1 − infection_mean_worm_burden_infected_by_exposure_regime_prior_history_square

p. 3: Figure S5.2.2 − infection_mean_worm_burden_infected_by_prior_stressor_history_bars_square

p. 4: Figure S5.2.3 − infection_total_worm_count_by_exposure_regime_prior_history_square

p. 5: Figure S5.2.4 − infection_total_worm_count_by_prior_stressor_history_bars_square

p. 6: Figure S5.7.1 − infection_prevalence_trend_predicted_square_quasirandom

p. 7: Figure S5.7.2 − infection_prevalence_by_exposure_regime_prior_history_square

p. 8: Figure S5.17.1 − mortality_prior_stressor_trend_square_quasirandom

p. 9: Figure S5.17.2 − mortality_percent_by_exposure_regime_prior_history_square

p. 10: Figure S5.17.3 − mortality_parasite_trend_comparison_square

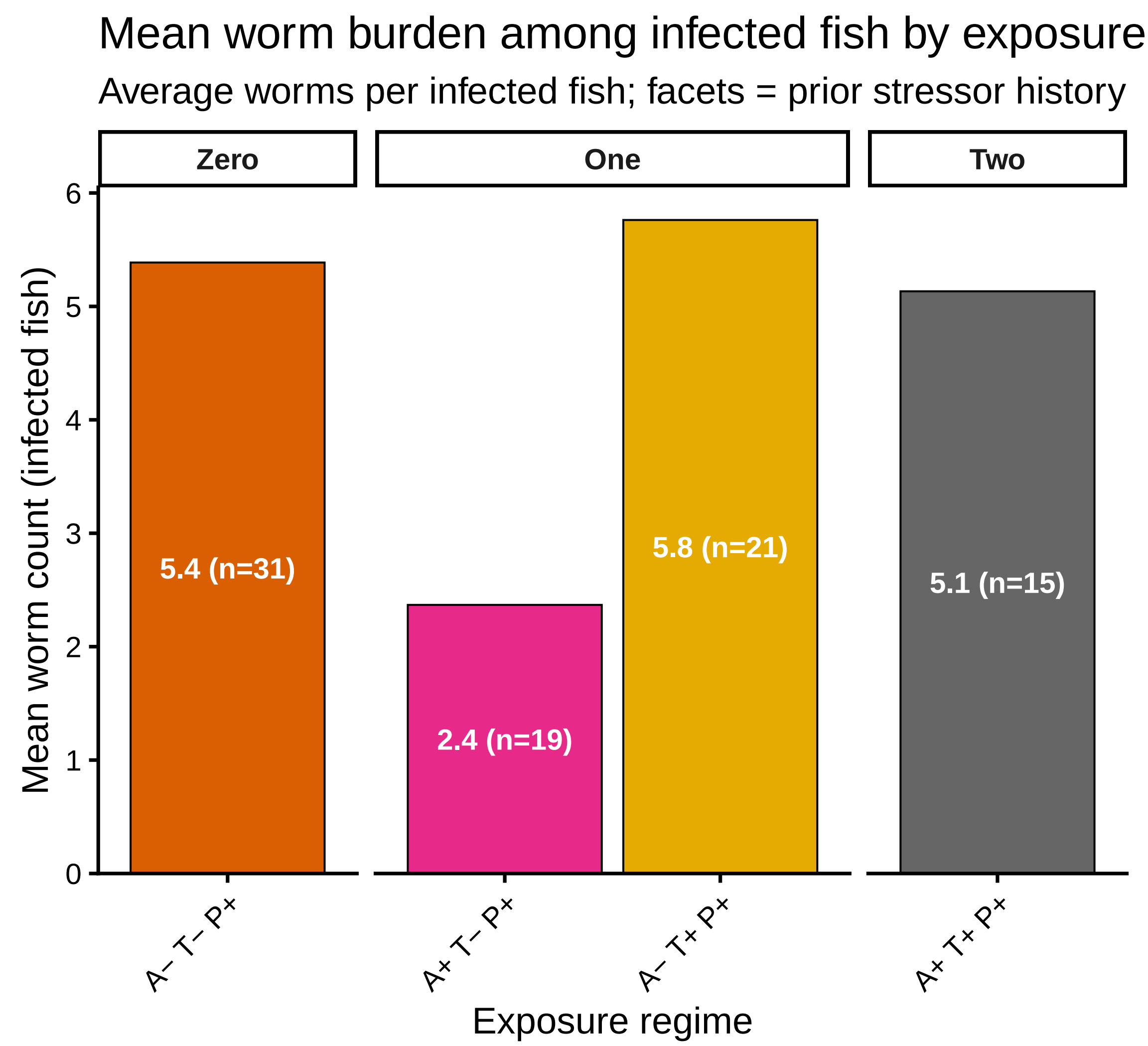

Mean among fish with Total.Worm.Count > 0 only; n = infected fish per cell.

Facets: prior stressor history; colors match exposure regime.

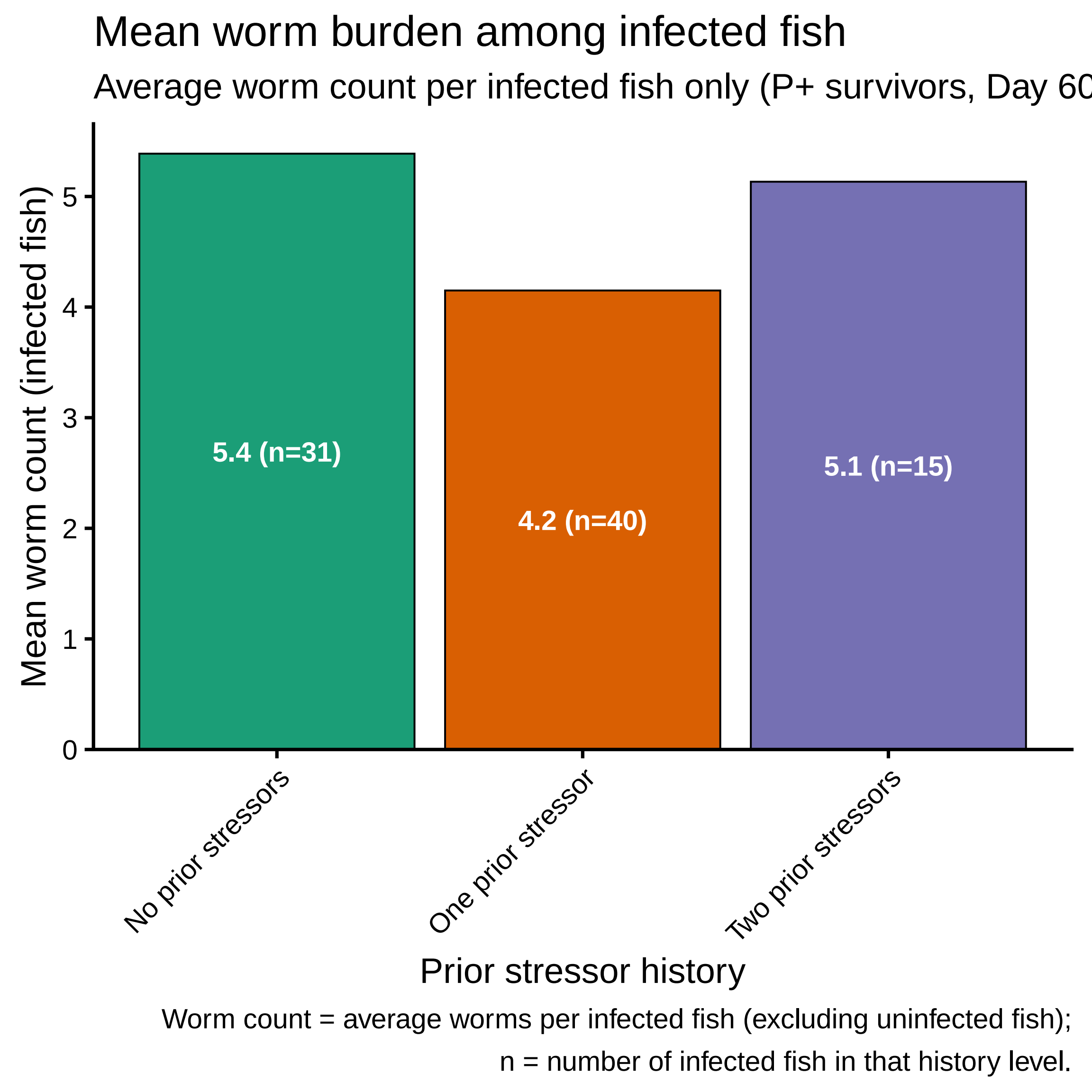
Mean worm burden among infected fish
Average worm count per infected fish only (P+ survivors, Day 60
Mean worm count (infected fish)
0
1
2
3
4
5
5.4 (n=31)
4.2 (n=40)
5.1 (n=15)
No prior stressors
One prior stressor
Two prior stressors
Prior stressor history
Worm count = average worms per infected fish (excluding uninfected fish);
n = number of infected fish in that history level.

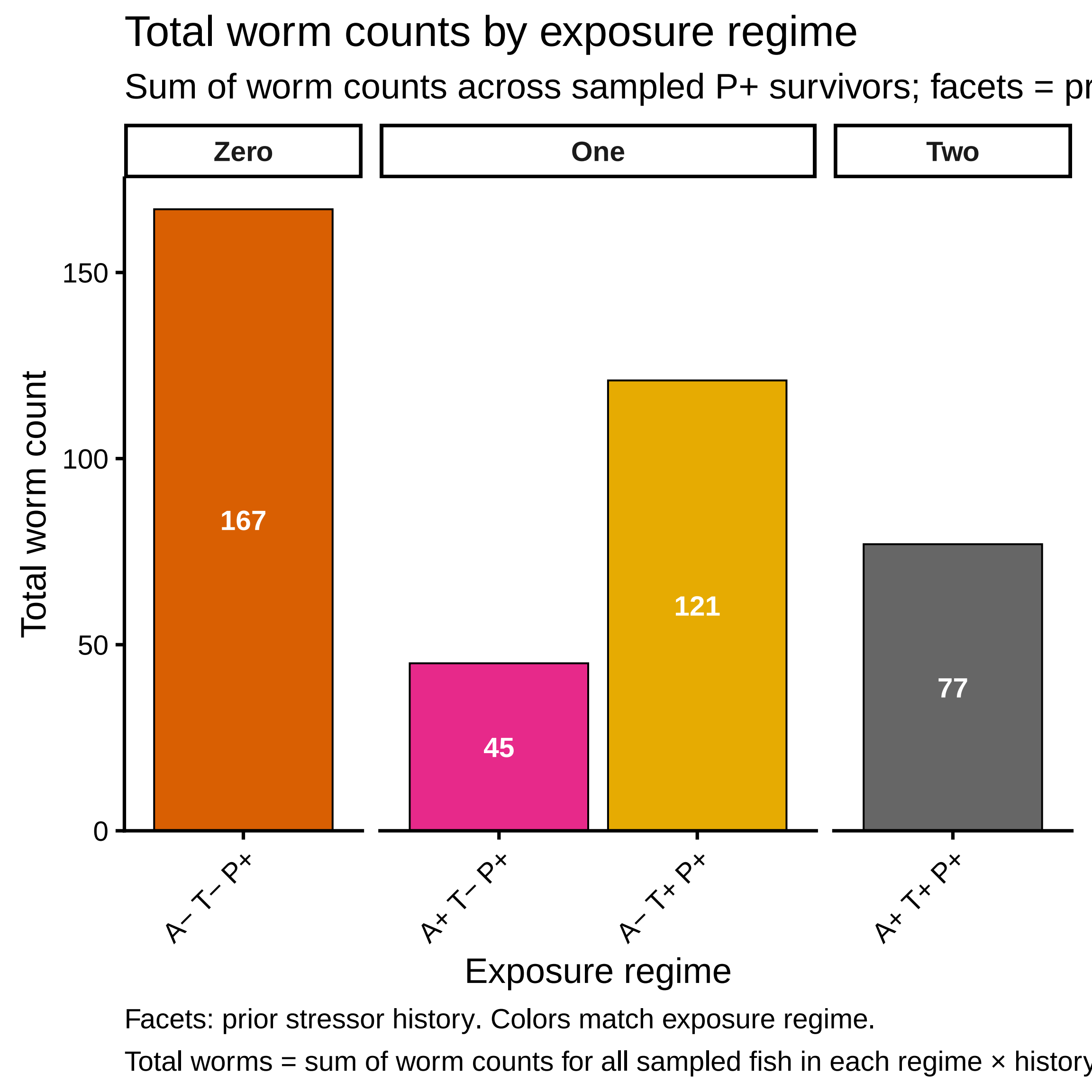

Total worm counts by exposure regime
Sum of worm counts across sampled P+ survivors; facets = pr
Zero
One
Two
Total worm count
150
100
50
0
167
45
121
77
A− T− P+
A+ T− P+
A− T+ P+
A+ T+ P+
Exposure regime
Facets: prior stressor history. Colors match exposure regime.
Total worms = sum of worm counts for all sampled fish in each regime × history

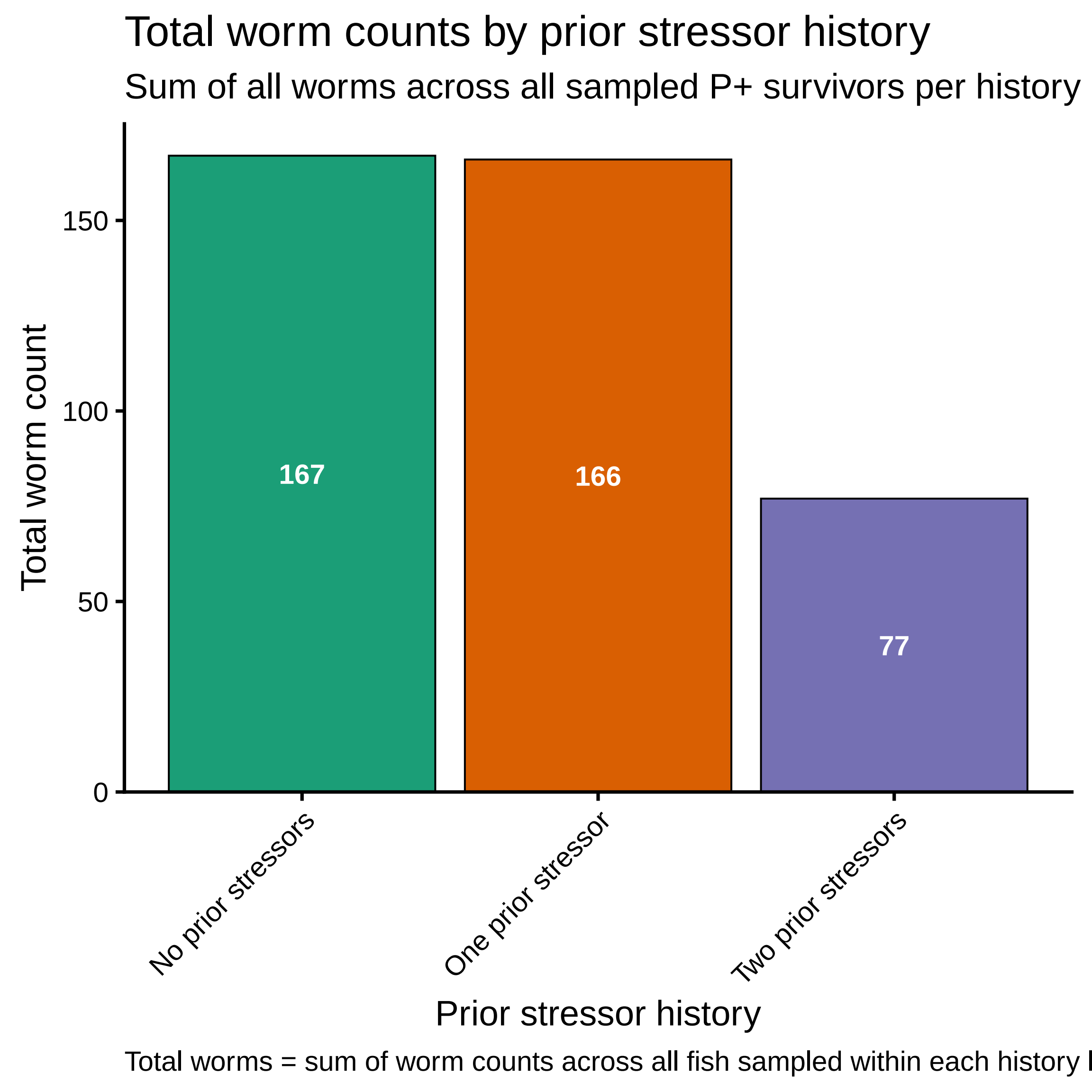
Total worm counts by prior stressor history
Sum of all worms across all sampled P+ survivors per history
Total worm count
150
100
50
0
167
166
77
No prior stressors
One prior stressor
Two prior stressors
Prior stressor history
Total worms = sum of worm counts across all fish sampled within each history l

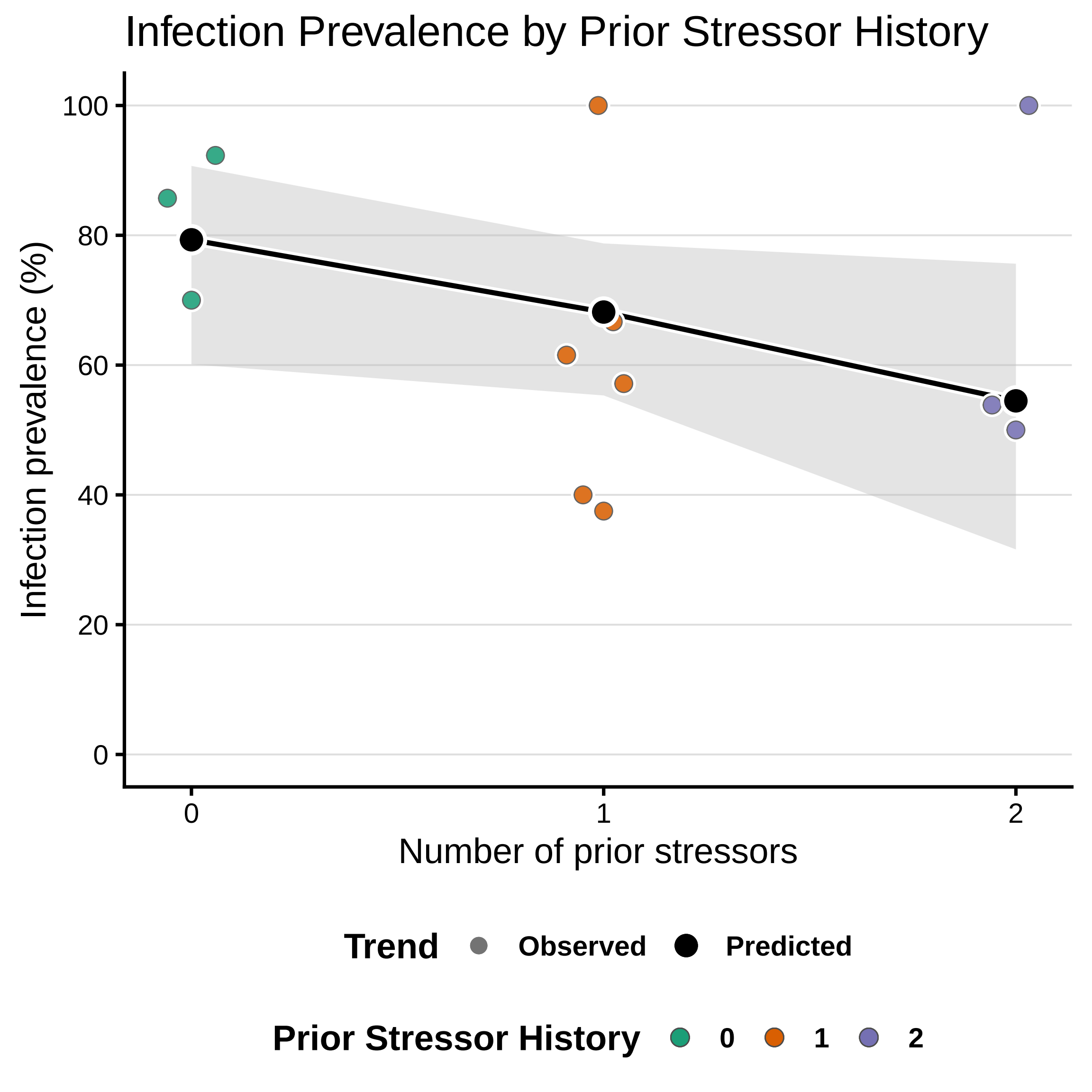
Infection Prevalence by Prior Stressor History
Infection prevalence (%)
100
80
60
40
20
0
0
1
2
Number of prior stressors
Trend
Observed
Predicted
Prior Stressor History
0
1
2

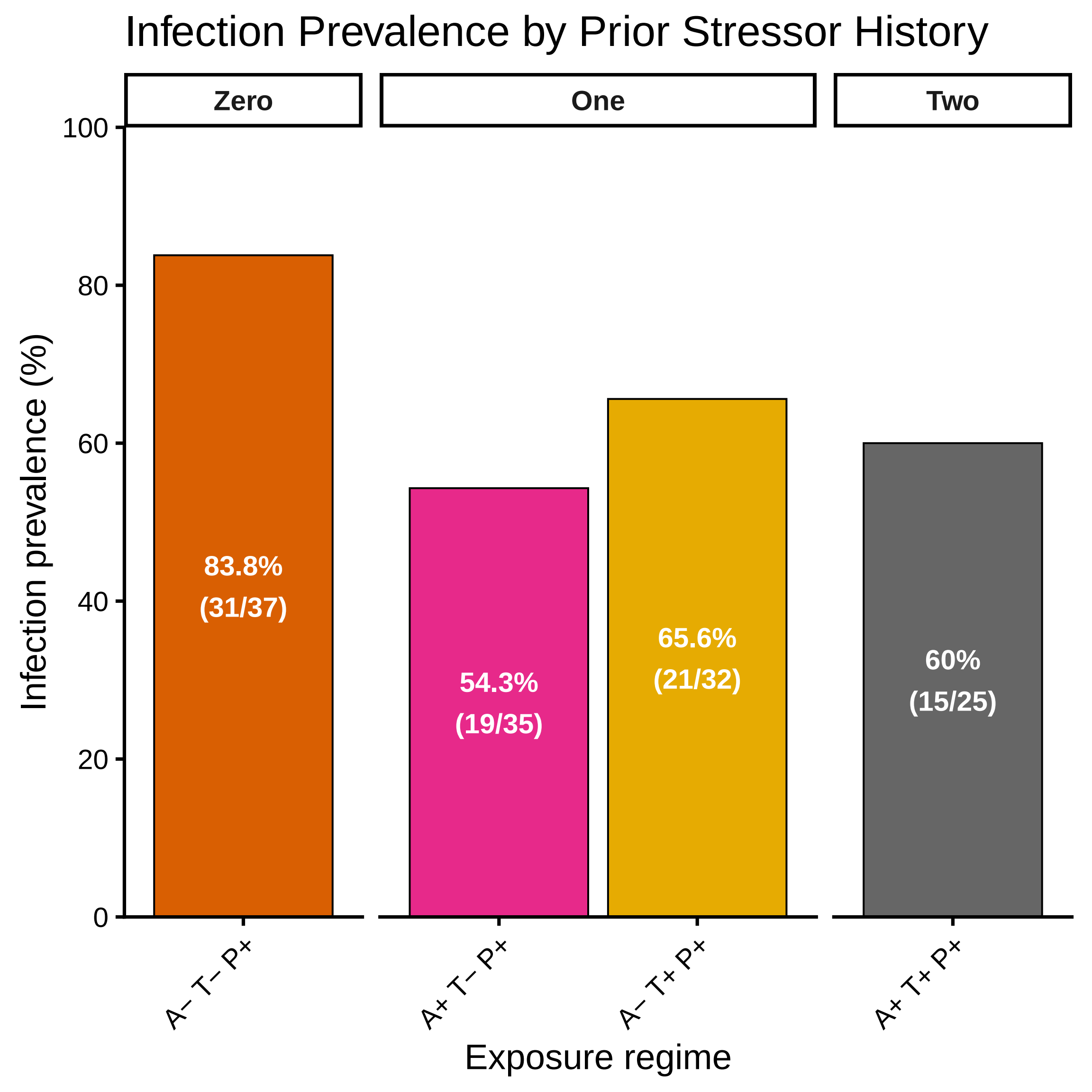
Infection Prevalence by Prior Stressor History
Zero
One
Two
Infection prevalence (%)
0
20
40
60
80
100
83.8%
(31/37)
54.3%
(19/35)
65.6%
(21/32)
60%
(15/25)
A− T− P+
A+ T− P+
A− T+ P+
A+ T+ P+
Exposure regime

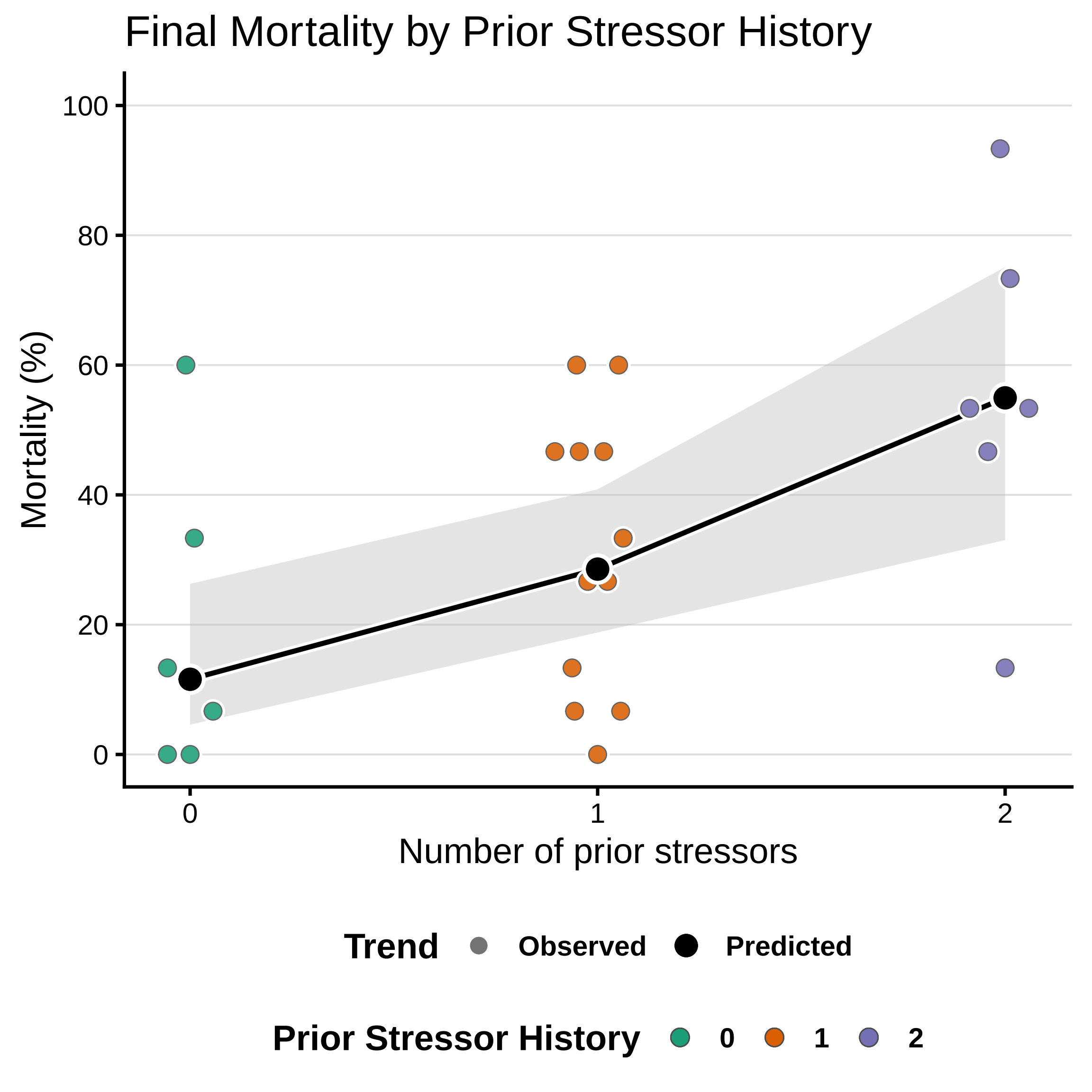

Final Mortality by Prior Stressor History
Mortality (%)
100
80
60
40
20
0
0
1
2
Number of prior stressors
Trend
Observed
Predicted
Prior Stressor History
0
1
2

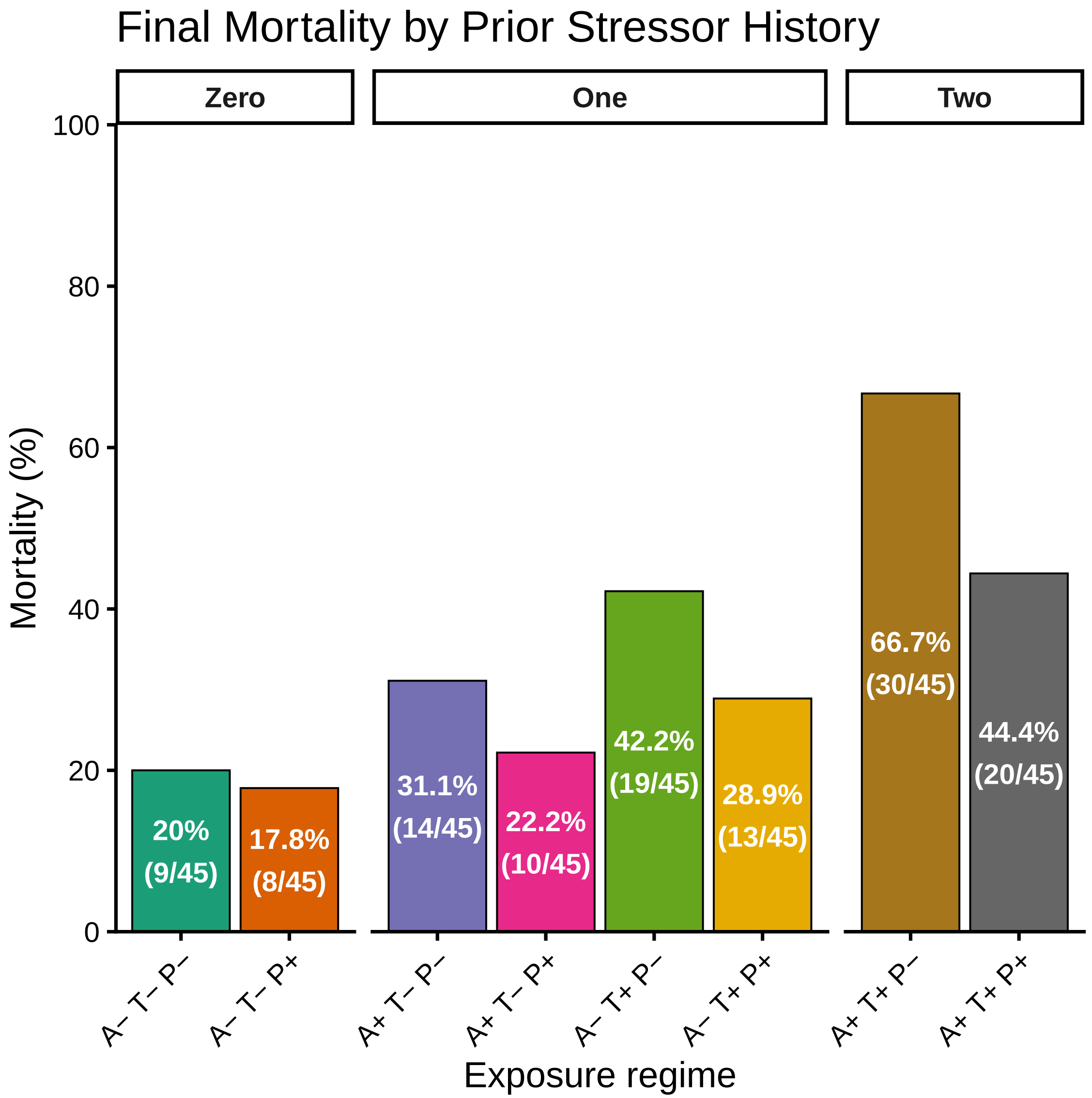

Final Mortality by Prior Stressor History
Zero
One
Two
Mortality (%)
100
80
60
40
20
0
20%
(9/45)
17.8%
(8/45)
31.1%
(14/45)
22.2%
(10/45)
42.2%
(19/45)
28.9%
(13/45)
66.7%
(30/45)
44.4%
(20/45)
A− T− P−
A− T− P+
A+ T− P−
A+ T− P+
A− T+ P−
A− T+ P+
A+ T+ P−
A+ T+ P+
Exposure regime

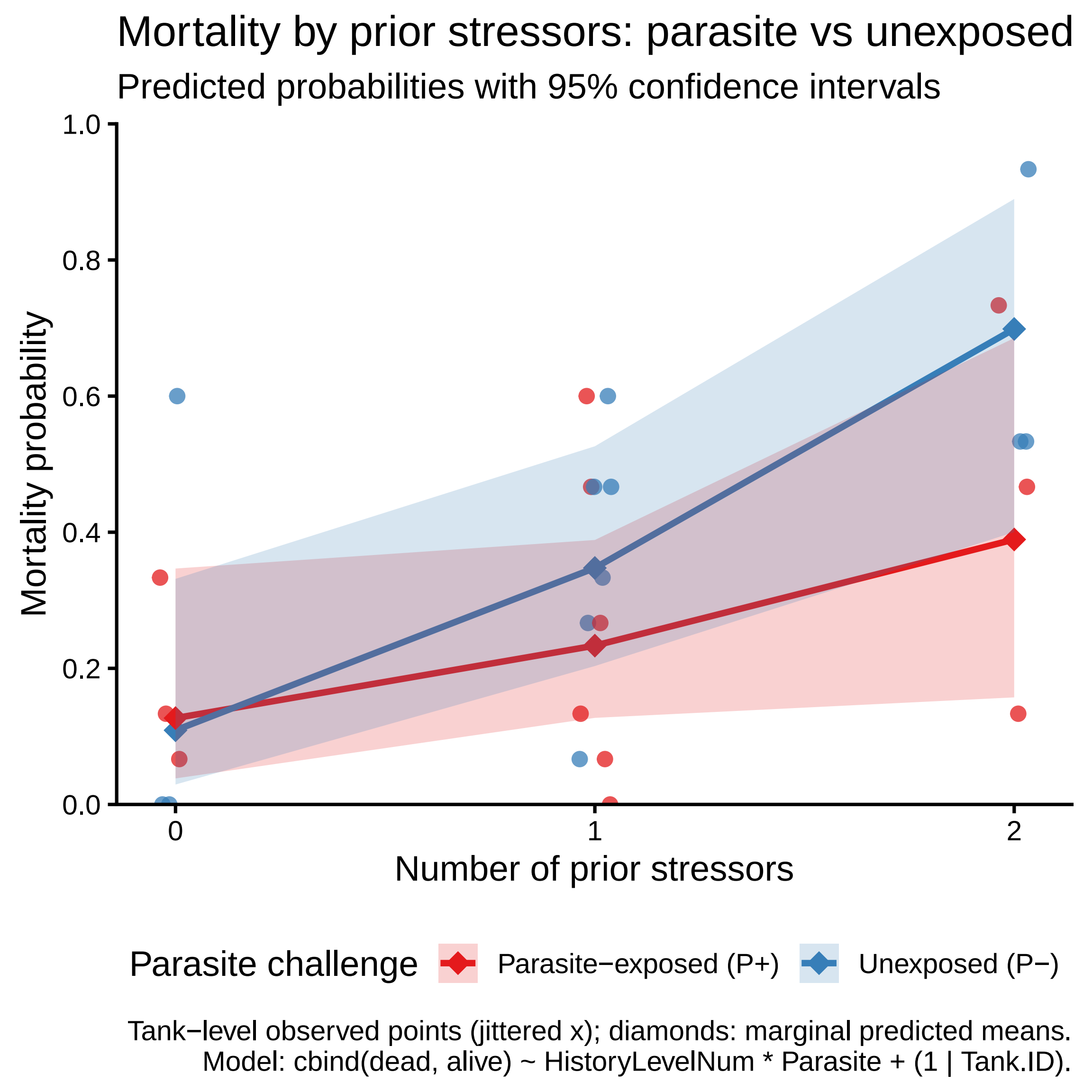
Mortality by prior stressors: parasite vs unexposed
Predicted probabilities with 95% confidence intervals
Mortality probability
1.0
0.8
0.6
0.4
0.2
0.0
0
1
2
Number of prior stressors
Parasite challenge
Parasite−exposed (P+)
Unexposed (P−)
Tank−level observed points (jittered x); diamonds: marginal predicted means.
Model: cbind(dead, alive) ~ HistoryLevelNum * Parasite + (1 | Tank.ID).
