## Supplementary Figures for "Historical contingency shapes zebrafish host-microbiome responses to a subsequent biotic challenge": Supplementary_Figures__06__Taxon-DEG-Mort__submission__2026-06-26.pdf

### Supplementary figures (combined)

Build (UTC): 2026−06−26 12:36:46 UTC

Git: 55d6b9a4a7bf134b3db15f0777c389e2da8c31be

Module: 06__Taxon−DEG−Mort

Figures dir: /Sieler2026/Results/06__Taxon−DEG−Mort/Figures

Figures (combined PDF order; p. = first page of that figure in this PDF):

p. 2: Figure S6.7.1 − all_sig_taxa_partial_cor_scatter

p. 3: Figure S6.8.1 − bipartite_partial_correlations

p. 4: Figure S6.4.1 − focal_four_genes_venn_all

p. 5: Figure S6.4.2 − focal_four_genes_venn_negative

p. 6: Figure S6.4.3 − focal_four_genes_venn_positive

p. 7: Figure S6.5.1 − partial_correlation_histogram

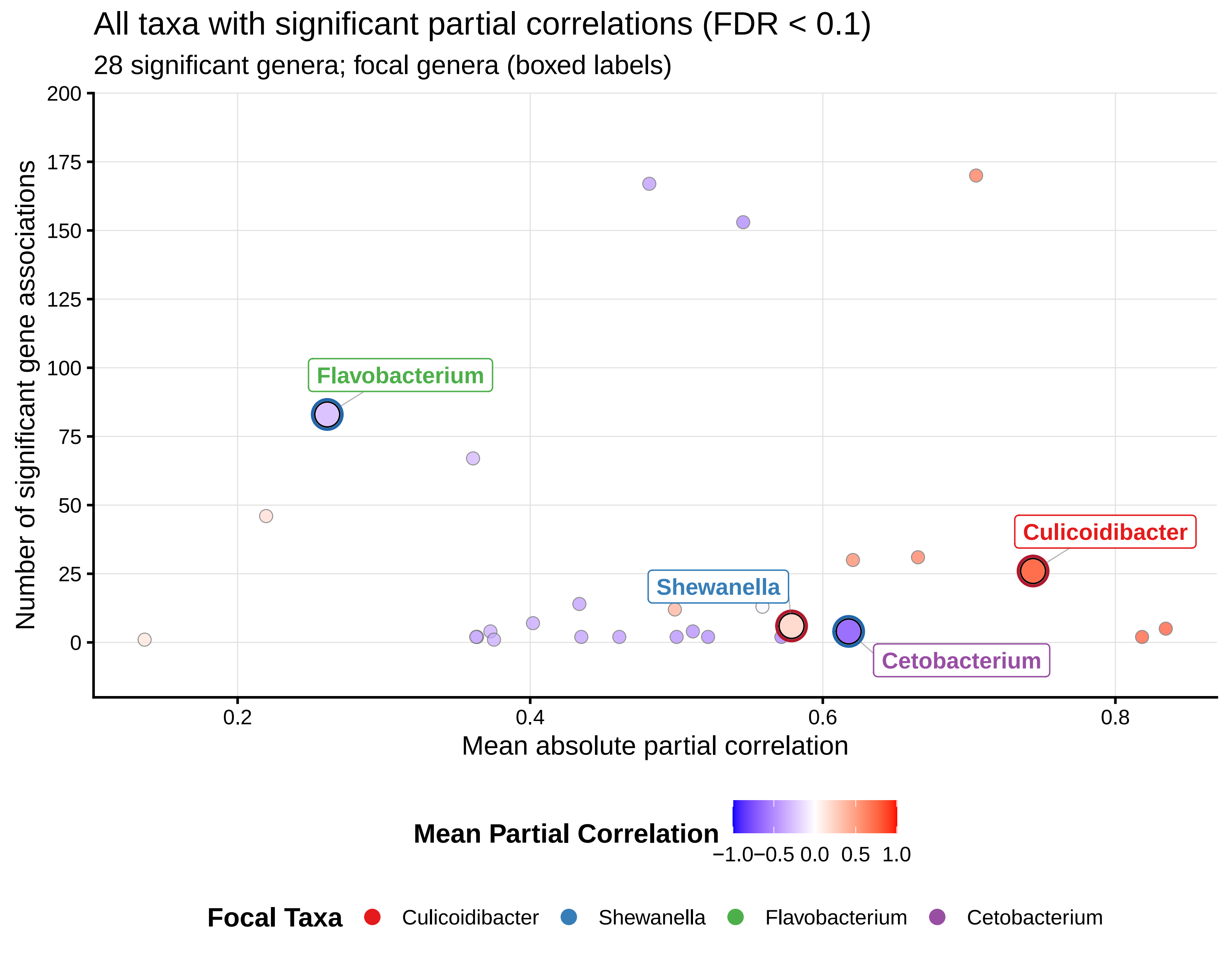

All taxa with significant partial correlations (FDR < 0.1)
28 significant genera; focal genera (boxed labels)
Number of significant gene associations
200
175
150
125
100
75
50
25
0
0.2
0.4
0.6
0.8
Mean absolute partial correlation
Flavobacterium
Shewanella
Culicoidibacter
Cetobacterium
Mean Partial Correlation
−1.0
−0.5
0.0
0.5
1.0
Focal Taxa
Culicoidibacter
Shewanella
Flavobacterium
Cetobacterium

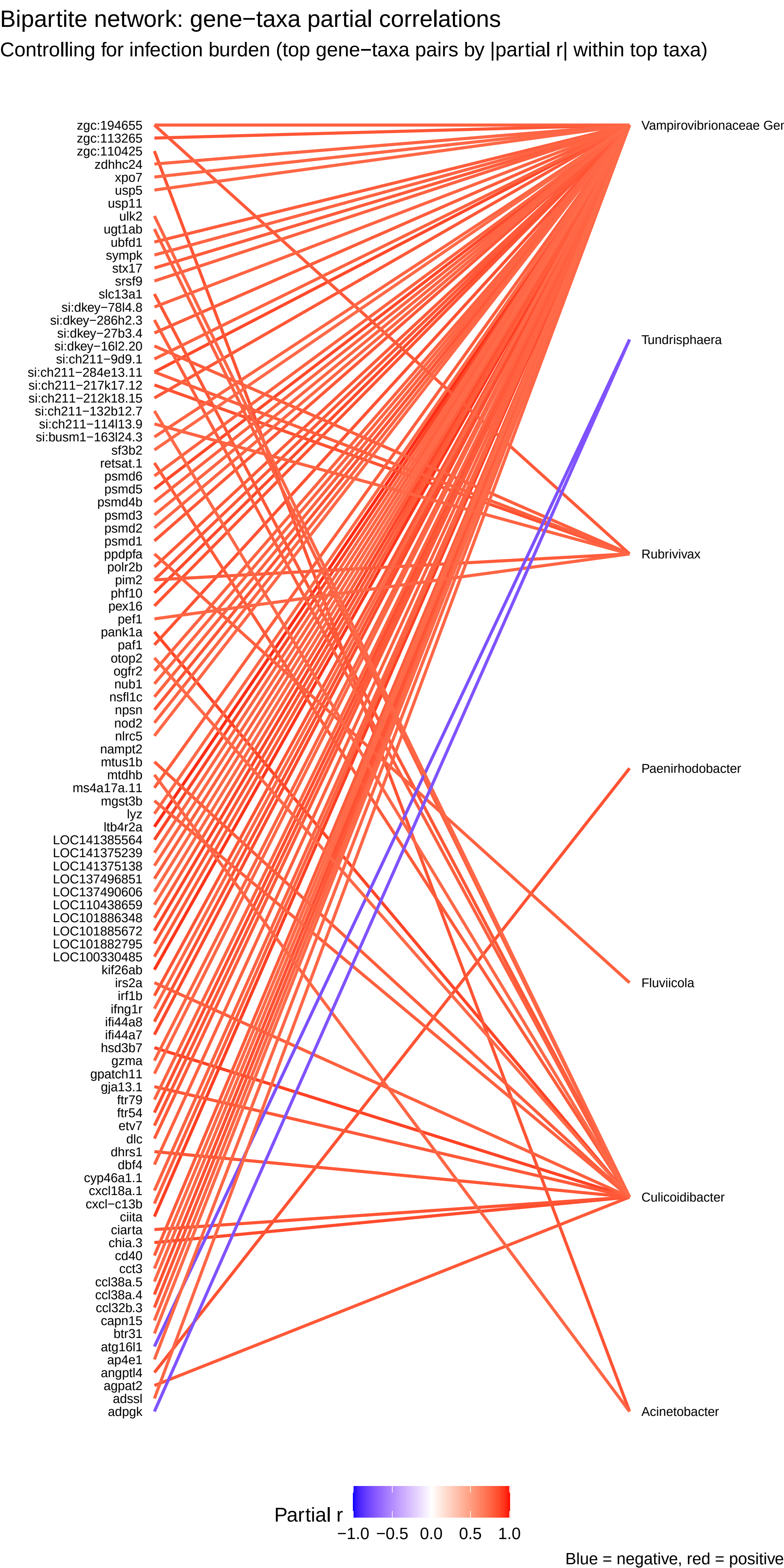
Bipartite network: gene−taxa partial correlations
Controlling for infection burden (top gene−taxa pairs by |partial r| within top taxa)
zgc:194655
zgc:113265
zgc:110425
zdhhc24
xpo7
usp5
usp11
ulk2
ugt1ab
ubfd1
sympk
stx17
srsf9
slc13a1
si:dkey−78l4.8
si:dkey−286h2.3
si:dkey−27b3.4
si:dkey−16l2.20
si:ch211−9d9.1
si:ch211−284e13.11
si:ch211−217k17.12
si:ch211−212k18.15
si:ch211−132b12.7
si:ch211−114l13.9
si:busm1−163l24.3
sf3b2
retsat.1
psmd6
psmd5
psmd4b
psmd3
psmd2
psmd1
ppdpfa
polr2b
pim2
phf10
pex16
pef1
pank1a
paf1
otop2
ogfr2
nub1
nsfl1c
npsn
nod2
nlrc5
nampt2
mtus1b
mtdhb
ms4a17a.11
mgst3b
lyz
ltb4r2a
LOC141385564
LOC141375239
LOC141375138
LOC137496851
LOC137490606
LOC110438659
LOC101886348
LOC101885672
LOC101882795
LOC100330485
kif26ab
irs2a
irf1b
ifng1r
ifi44a8
ifi44a7
hsd3b7
gzma
gpatch11
gja13.1
ftr79
ftr54
etv7
dlc
dhrs1
dbf4
cyp46a1.1
cxcl18a.1
cxcl−c13b
ciita
ciarta
chia.3
cd40
cct3
ccl38a.5
ccl38a.4
ccl32b.3
capn15
btr31
atg16l1
ap4e1
angptl4
agpat2
adssl
adpgk
Vampirovibrionaceae Ger
Tundrisphaera
Rubrivivax
Paenirhodobacter
Fluviicola
Culicoidibacter
Acinetobacter
Partial r
−1.0 −0.5 0.0 0.5 1.0
Blue = negative, red = positive

#### Gene overlap: focal genera (all significant edges)

FDR−significant partial correlations to each genus
(module 06).

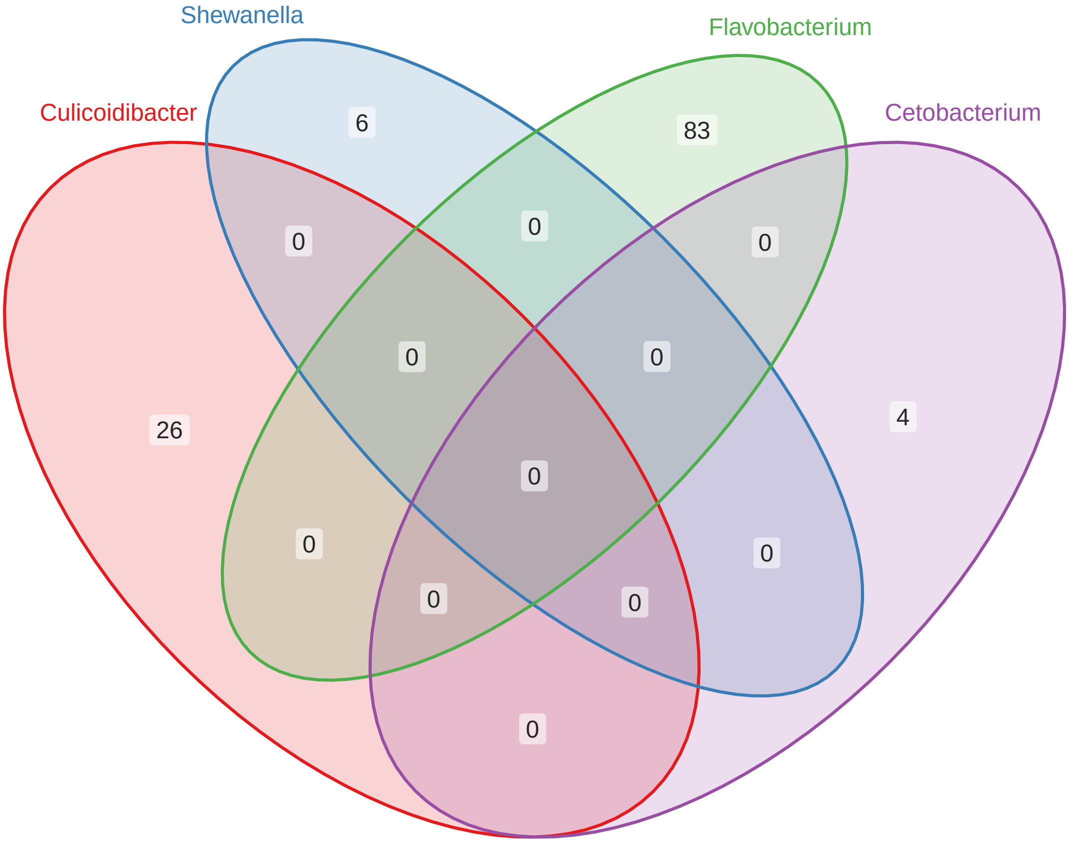

#### Gene overlap: focal genera (negative partial correlation)

At least one negative partial correlation per genus (FDR < 0.1 edges).

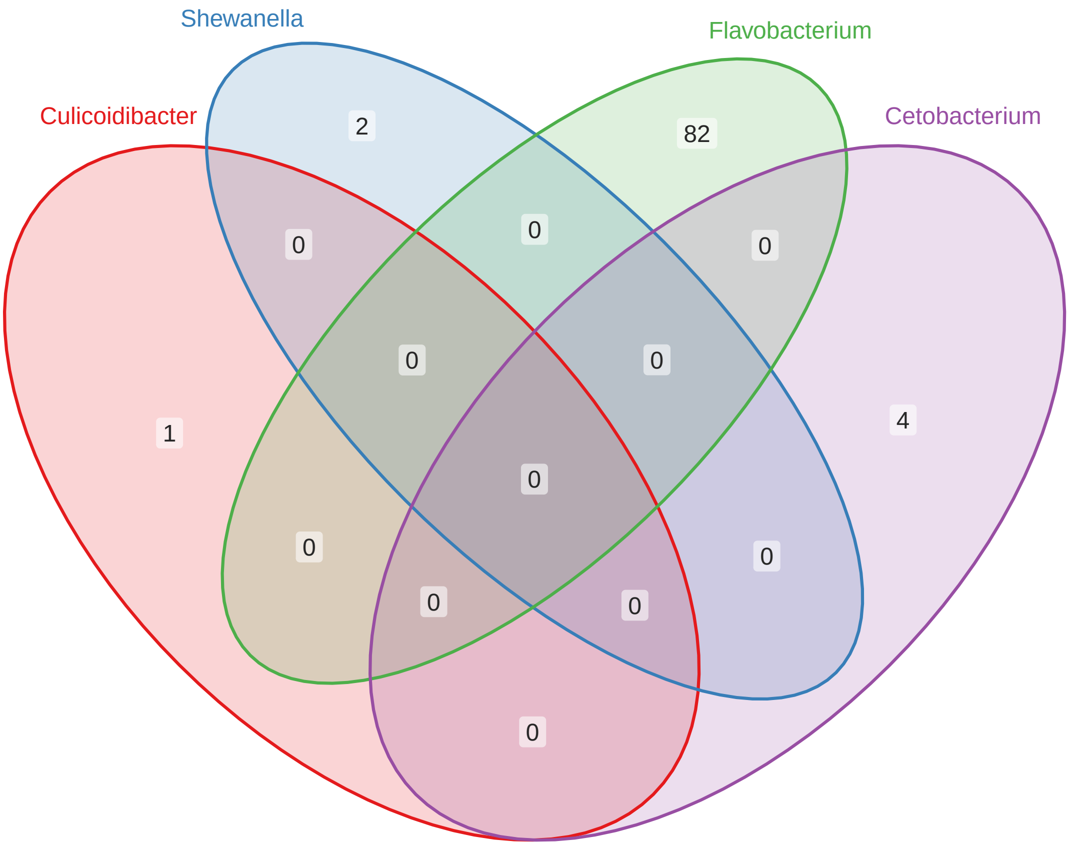

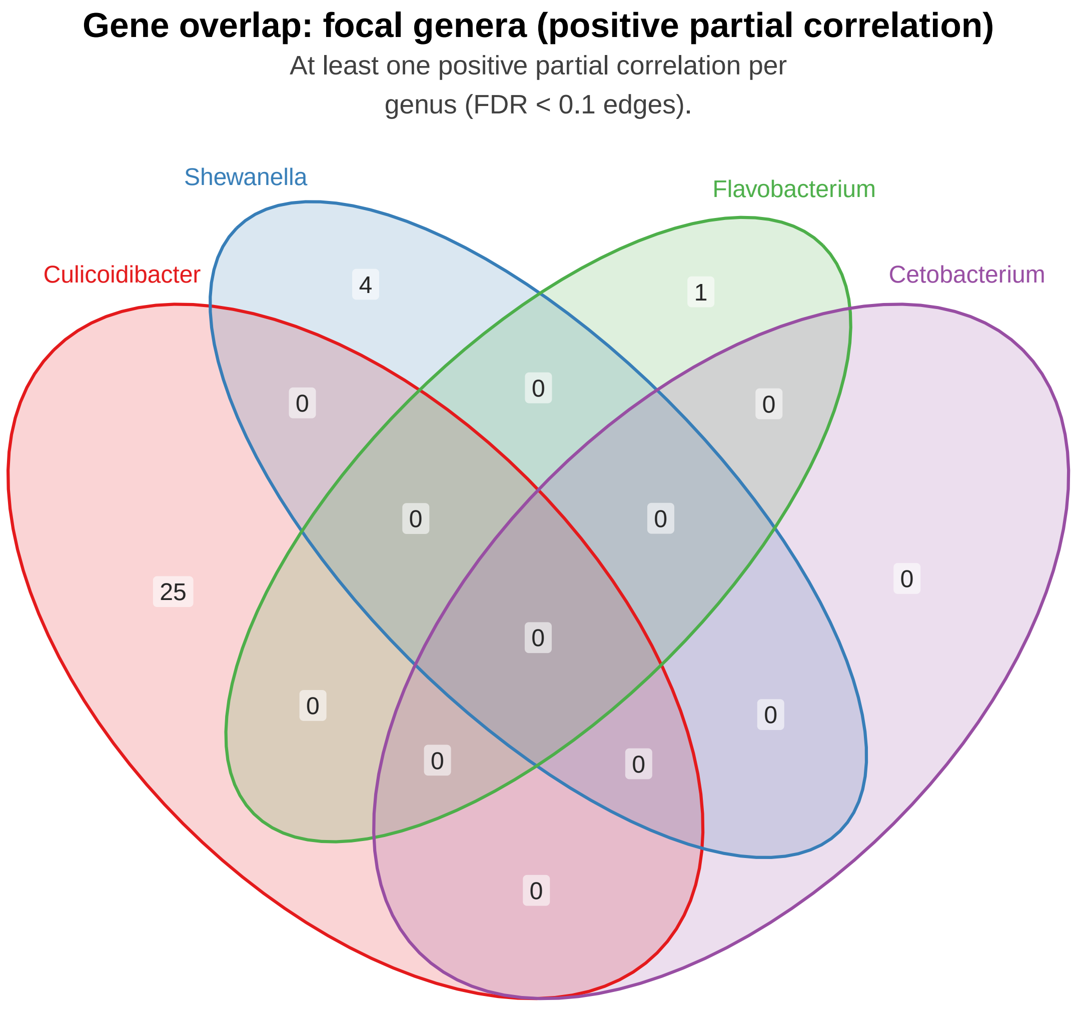
Gene overlap: focal genera (positive partial correlation)
At least one positive partial correlation per
genus (FDR < 0.1 edges).
Shewanella
Flavobacterium
Culicoidibacter
Cetobacterium
4
1
0
0
0
0
0
25
0
0
0
0
0
0
0

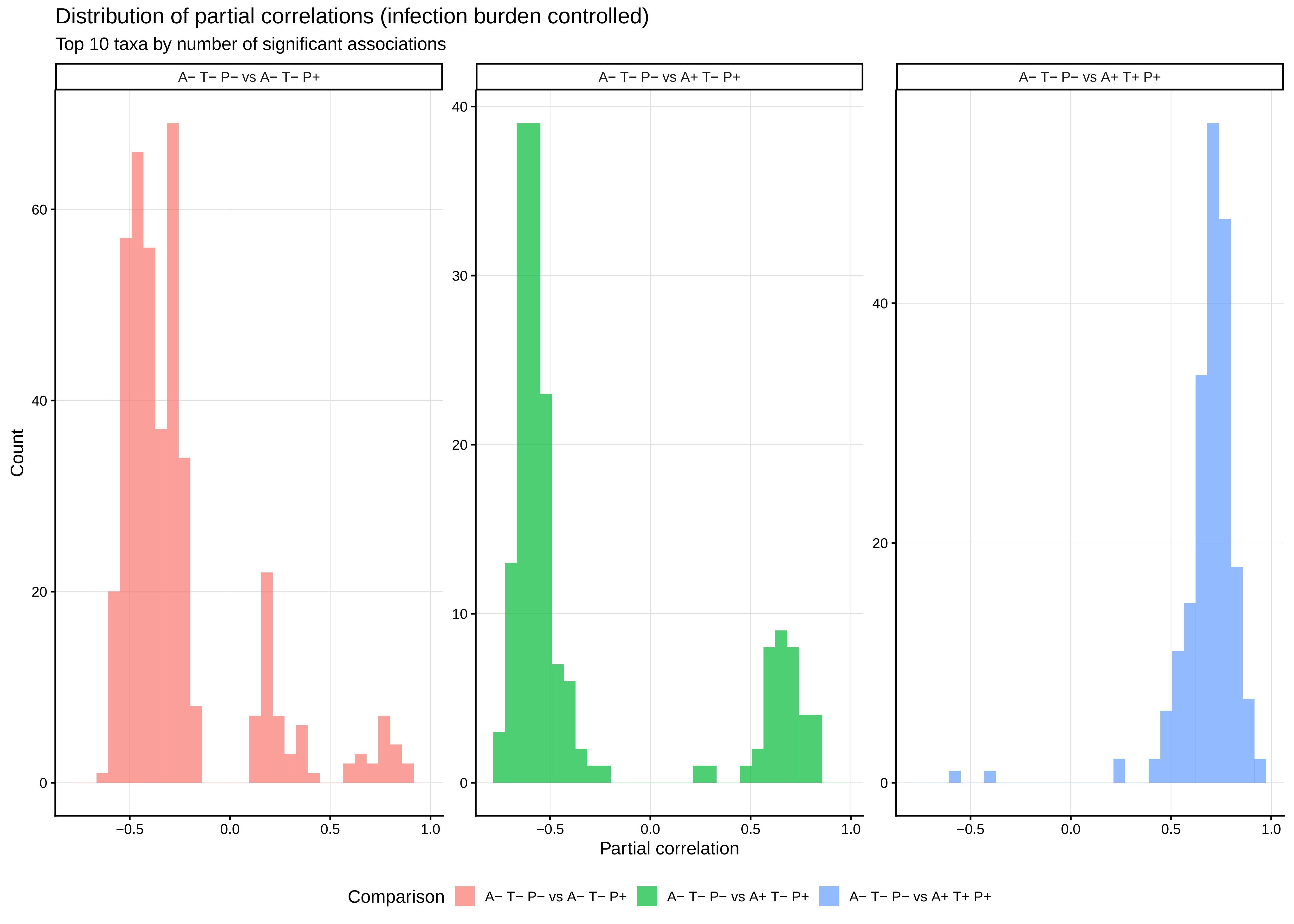

Distribution of partial correlations (infection burden controlled)
Top 10 taxa by number of significant associations
A− T− P− vs A− T− P+
A− T− P− vs A+ T− P+
A− T− P− vs A+ T+ P+
Count
Partial correlation
0
20
40
60
10
30
−0.5
0.0
0.5
1.0
Comparison
A− T− P− vs A− T− P+
A− T− P− vs A+ T− P+
A− T− P− vs A+ T+ P+
