## Supplementary Figures for "Historical contingency shapes zebrafish host-microbiome responses to a subsequent biotic challenge": Supplementary_Figures__07__FunctionalAnno__submission__2026-06-26.pdf

### Supplementary figures (combined)

Build (UTC): 2026−06−26 12:36:47 UTC

Git: 55d6b9a4a7bf134b3db15f0777c389e2da8c31be

Module: 07__FunctionalAnno

Figures dir: /Sieler2026/Results/07__FunctionalAnno/Figures

Figures (combined PDF order; p. = first page of that figure in this PDF):

p. 2: Figure S7.14.1 − go_bp_cetobacterium_dotplot

p. 3: Figure S7.16.1 − go_bp_flavobacterium_dotplot

p. 4: Figure S7.18.1 − go_bp_focal_four_dotplot

p. 5: Figure S7.19.1 − go_bp_shewanella_dotplot

p. 6: Figure S7.21.1 − go_cc_cetobacterium_dotplot

p. 7: Figure S7.22.1 − go_cc_culicoidibacter_dotplot

p. 8: Figure S7.23.1 − go_cc_flavobacterium_dotplot

p. 9: Figure S7.25.1 − go_cc_focal_four_dotplot

p. 10: Figure S7.26.1 − go_cc_shewanella_dotplot

p. 11: Figure S7.28.1 − go_mf_cetobacterium_dotplot

p. 12: Figure S7.29.1 − go_mf_culicoidibacter_dotplot

p. 13: Figure S7.30.1 − go_mf_flavobacterium_dotplot

p. 14: Figure S7.32.1 − go_mf_focal_four_dotplot

p. 15: Figure S7.33.1 − go_mf_shewanella_dotplot

p. 16: Figure S7.35.1 − kegg_cetobacterium_dotplot

p. 17: Figure S7.36.1 − kegg_culicoidibacter_dotplot

p. 18: Figure S7.37.1 − kegg_flavobacterium_dotplot

p. 19: Figure S7.39.1 − kegg_focal_four_dotplot

p. 20: Figure S7.40.1 − kegg_shewanella_dotplot

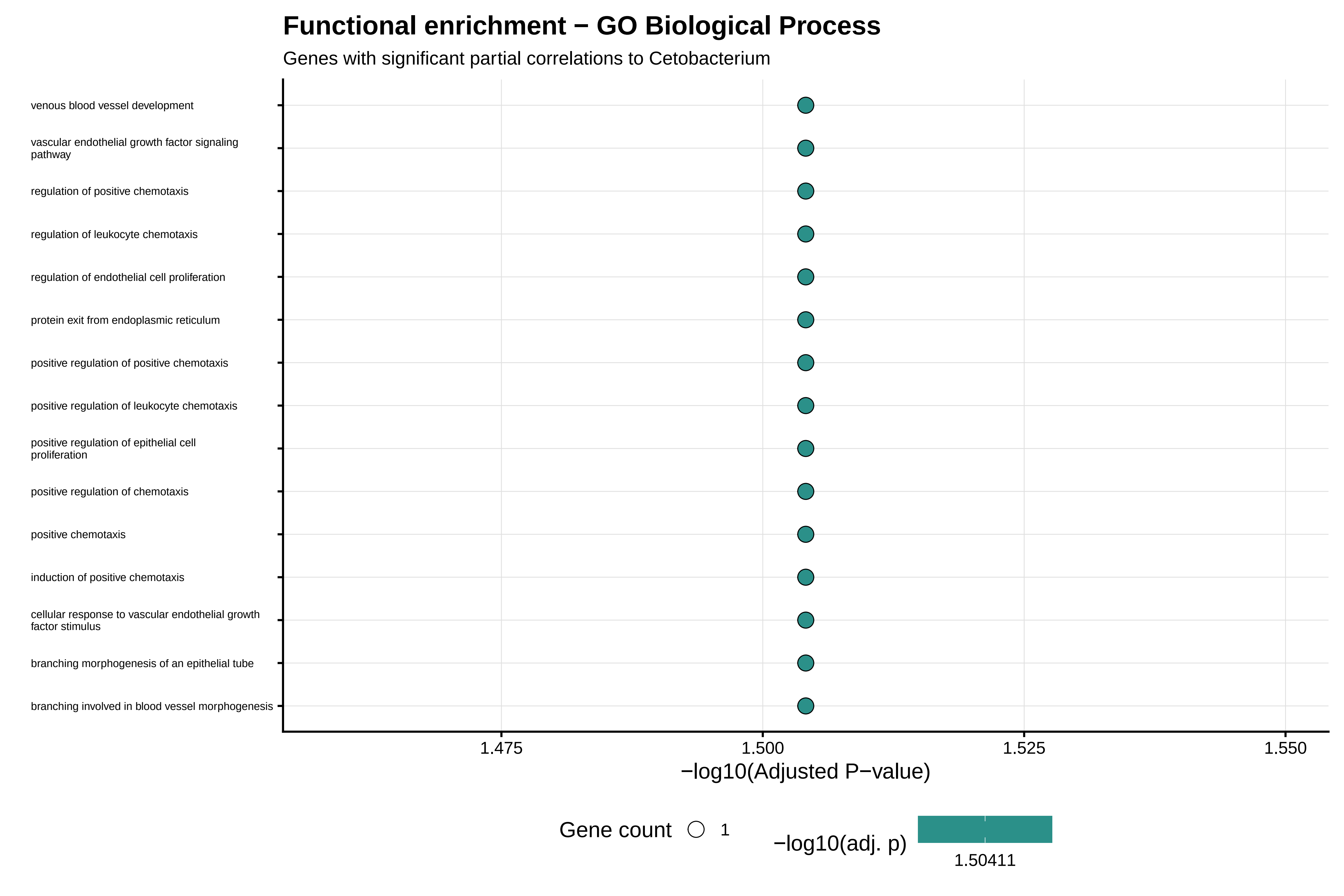

Functional enrichment − GO Biological Process
Genes with significant partial correlations to Cetobacterium
venous blood vessel development
vascular endothelial growth factor signaling pathway
regulation of positive chemotaxis
regulation of leukocyte chemotaxis
regulation of endothelial cell proliferation
protein exit from endoplasmic reticulum
positive regulation of positive chemotaxis
positive regulation of leukocyte chemotaxis
positive regulation of epithelial cell proliferation
positive regulation of chemotaxis
positive chemotaxis
induction of positive chemotaxis
cellular response to vascular endothelial growth factor stimulus
branching morphogenesis of an epithelial tube
branching involved in blood vessel morphogenesis
1.475
1.500
1.525
1.550
−log10(Adjusted P−value)
Gene count
1
−log10(adj. p)
1.50411

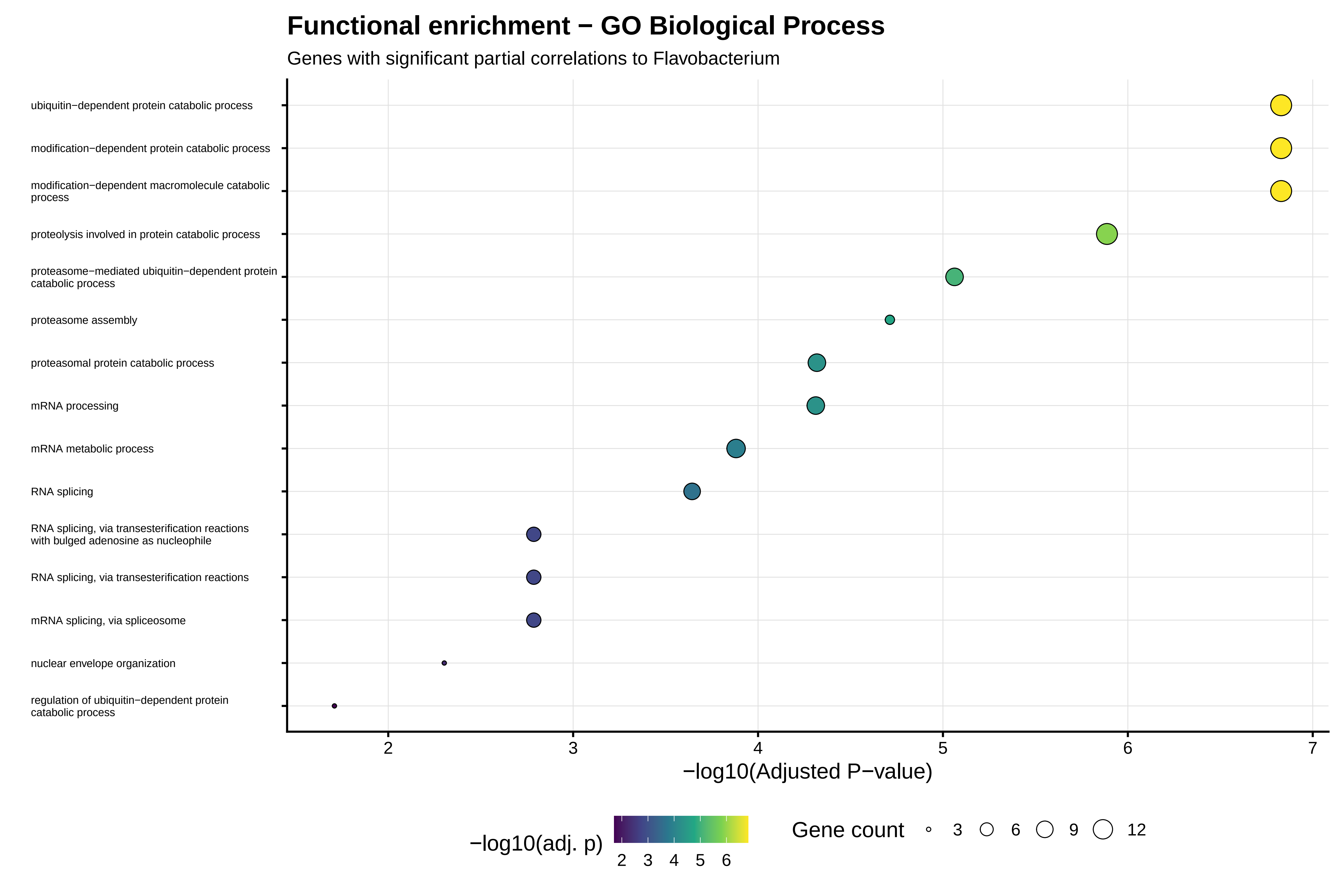
Functional enrichment − GO Biological Process
Genes with significant partial correlations to Flavobacterium
ubiquitin−dependent protein catabolic process
modification−dependent protein catabolic process
modification−dependent macromolecule catabolic process
proteolysis involved in protein catabolic process
proteasome−mediated ubiquitin−dependent protein catabolic process
proteasome assembly
proteasomal protein catabolic process
mRNA processing
mRNA metabolic process
RNA splicing
RNA splicing, via transesterification reactions with bulged adenosine as nucleophile
RNA splicing, via transesterification reactions
mRNA splicing, via spliceosome
nuclear envelope organization
regulation of ubiquitin−dependent protein catabolic process
2
3
4
5
6
7
−log10(Adjusted P−value)
−log10(adj. p)
2 3 4 5 6
Gene count
3
6
9
12

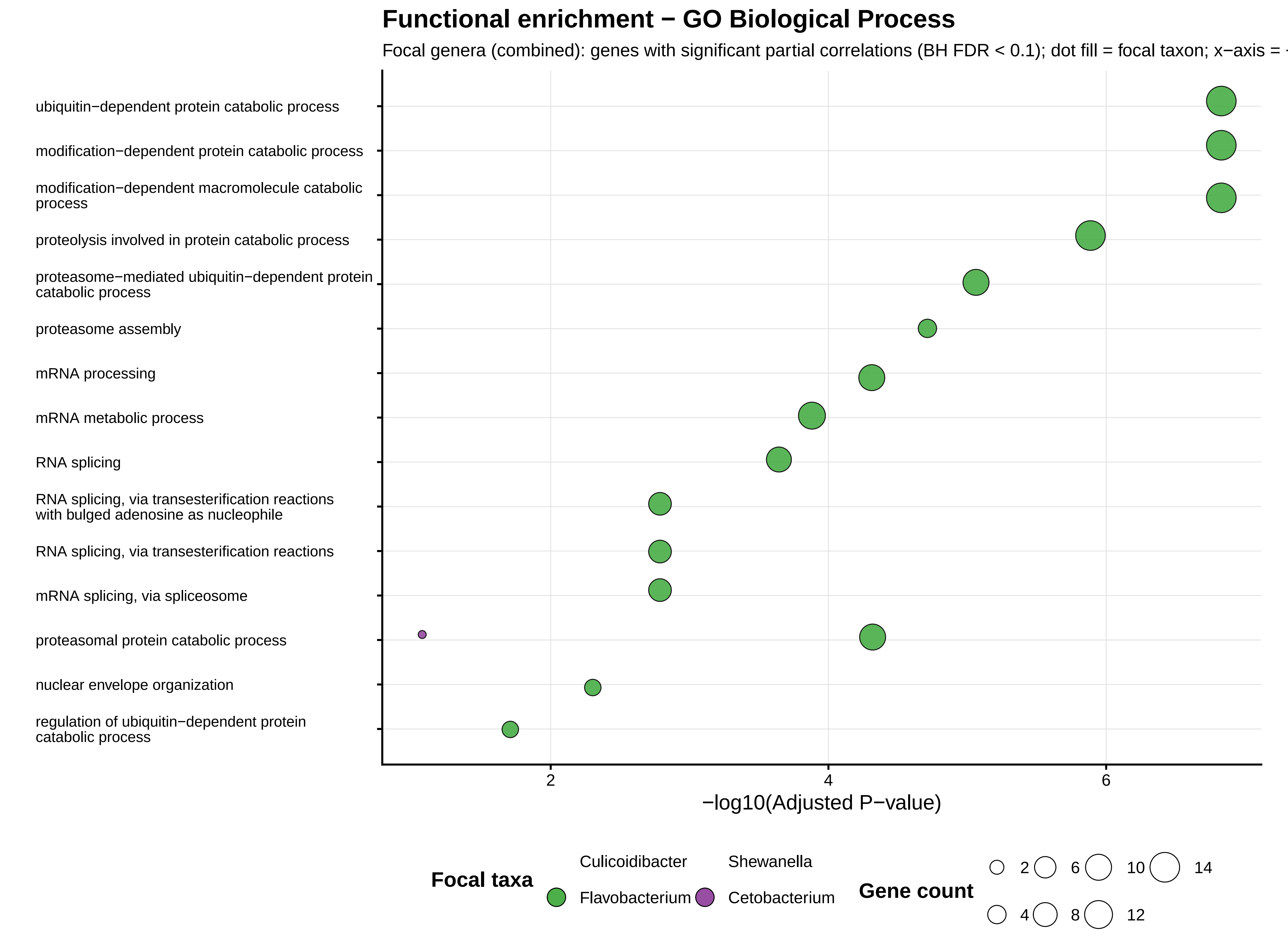
Functional enrichment − GO Biological Process
Focal genera (combined): genes with significant partial correlations (BH FDR < 0.1); dot fill = focal taxon; x−axis =
ubiquitin−dependent protein catabolic process
modification−dependent protein catabolic process
modification−dependent macromolecule catabolic process
proteolysis involved in protein catabolic process
proteasome−mediated ubiquitin−dependent protein catabolic process
proteasome assembly
mRNA processing
mRNA metabolic process
RNA splicing
RNA splicing, via transesterification reactions with bulged adenosine as nucleophile
RNA splicing, via transesterification reactions
mRNA splicing, via spliceosome
proteasomal protein catabolic process
nuclear envelope organization
regulation of ubiquitin−dependent protein catabolic process
2
4
6
−log10(Adjusted P−value)
Focal taxa
Culicoidibacter
Shewanella
Flavobacterium
Cetobacterium
Gene count
2
4
6
8
10
12
14

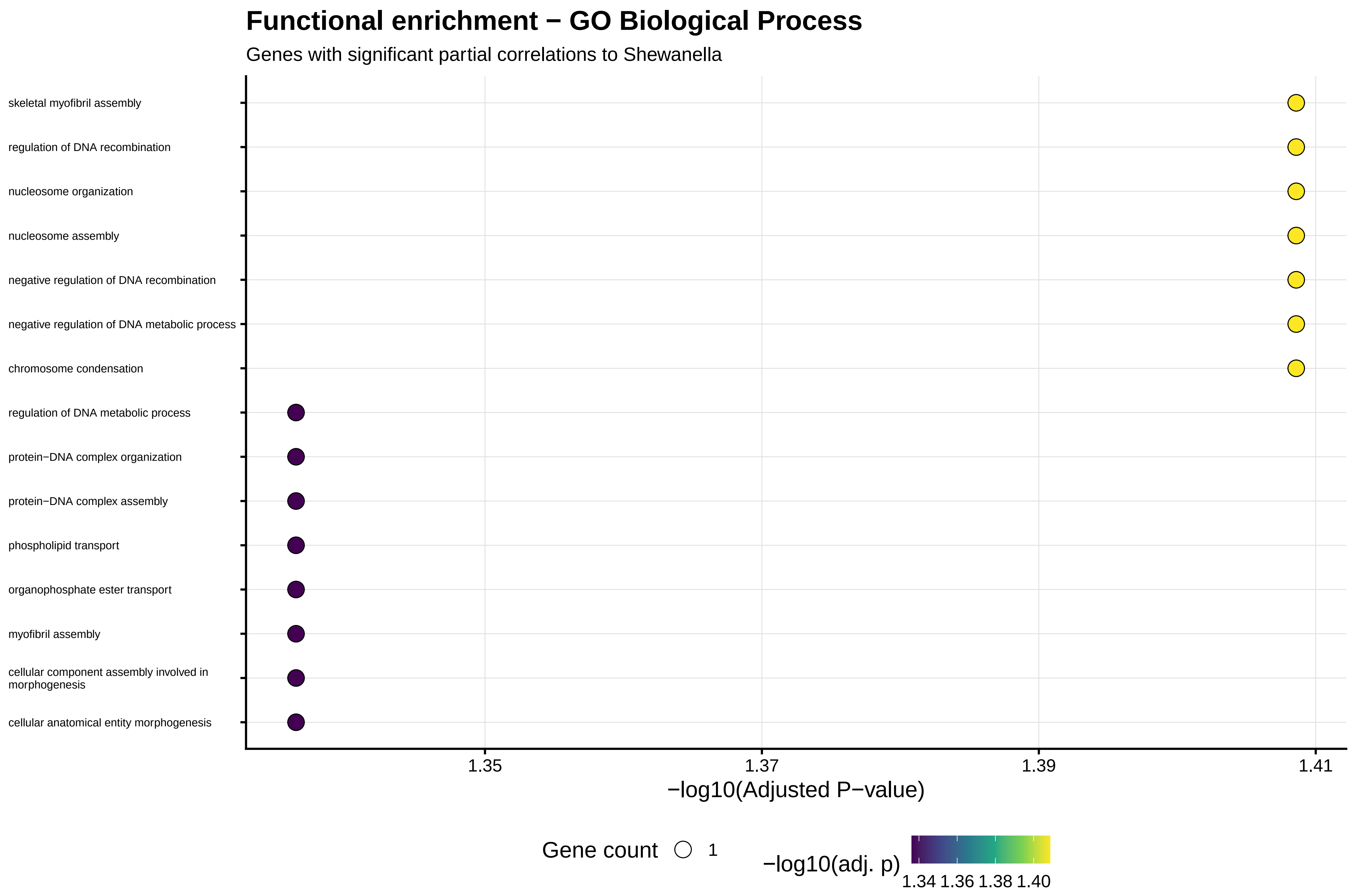
Functional enrichment − GO Biological Process
Genes with significant partial correlations to Shewanella
skeletal myofibril assembly
regulation of DNA recombination
nucleosome organization
nucleosome assembly
negative regulation of DNA recombination
negative regulation of DNA metabolic process
chromosome condensation
regulation of DNA metabolic process
protein−DNA complex organization
protein−DNA complex assembly
phospholipid transport
organophosphate ester transport
myofibril assembly
cellular component assembly involved in morphogenesis
cellular anatomical entity morphogenesis
1.35
1.37
1.39
1.41
−log10(Adjusted P−value)
Gene count
1
−log10(adj. p)
1.34 1.36 1.38 1.40

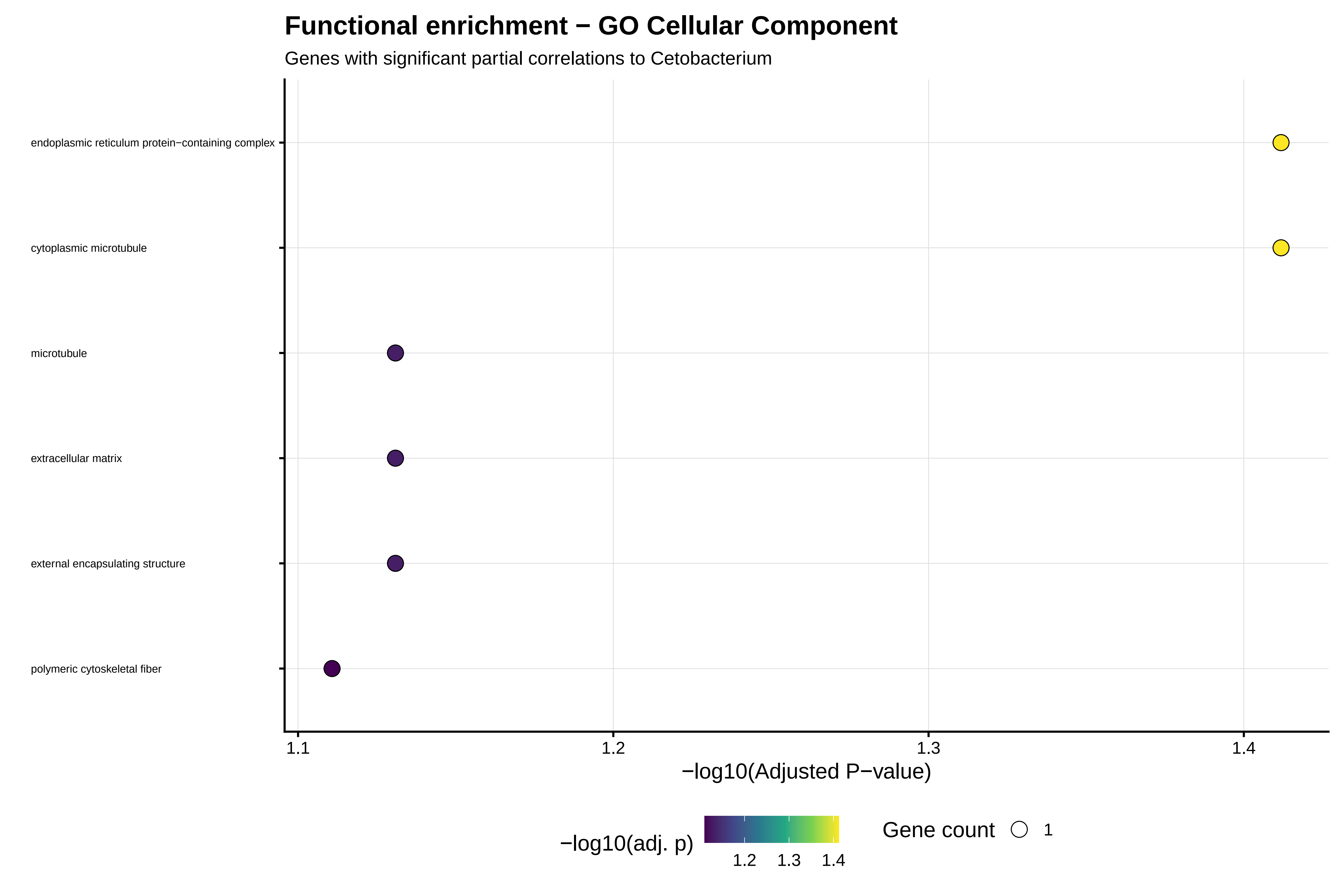
Functional enrichment − GO Cellular Component
Genes with significant partial correlations to Cetobacterium
endoplasmic reticulum protein−containing complex
cytoplasmic microtubule
microtubule
extracellular matrix
external encapsulating structure
polymeric cytoskeletal fiber
1.1
1.2
1.3
1.4
−log10(Adjusted P−value)
−log10(adj. p)
1.2
1.3
1.4
Gene count
1

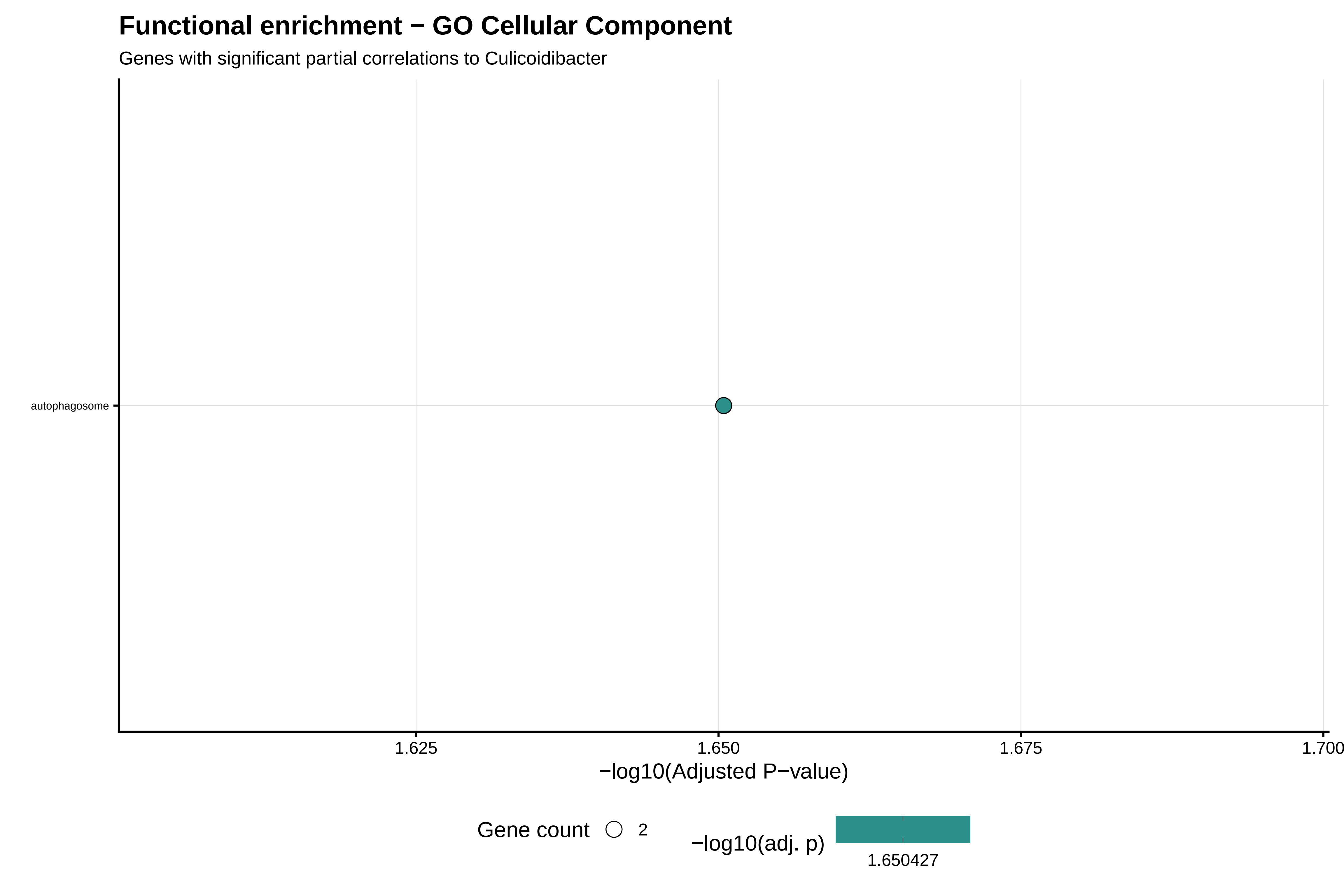
Functional enrichment − GO Cellular Component
Genes with significant partial correlations to Culicoidibacter
autophagosome
1.625
1.650
1.675
1.700
−log10(Adjusted P−value)
Gene count
2
−log10(adj. p)
1.650427

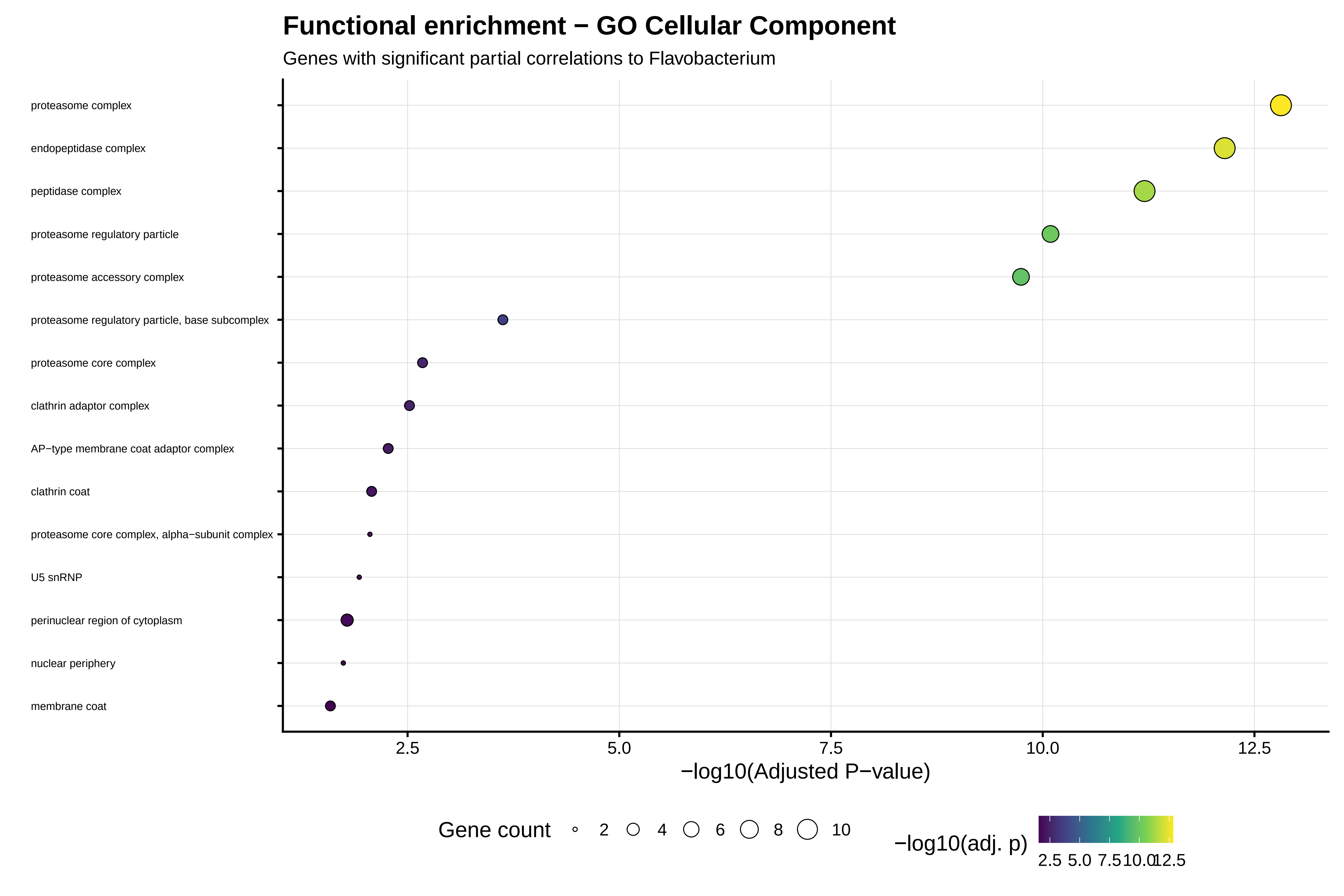
Functional enrichment − GO Cellular Component
Genes with significant partial correlations to Flavobacterium
proteasome complex
endopeptidase complex
peptidase complex
proteasome regulatory particle
proteasome accessory complex
proteasome regulatory particle, base subcomplex
proteasome core complex
clathrin adaptor complex
AP−type membrane coat adaptor complex
clathrin coat
proteasome core complex, alpha−subunit complex
U5 snRNP
perinuclear region of cytoplasm
nuclear periphery
membrane coat
2.5
5.0
7.5
10.0
12.5
−log10(Adjusted P−value)
Gene count
2
4
6
8
10
−log10(adj. p)
2.5 5.0 7.5 10.0 12.5

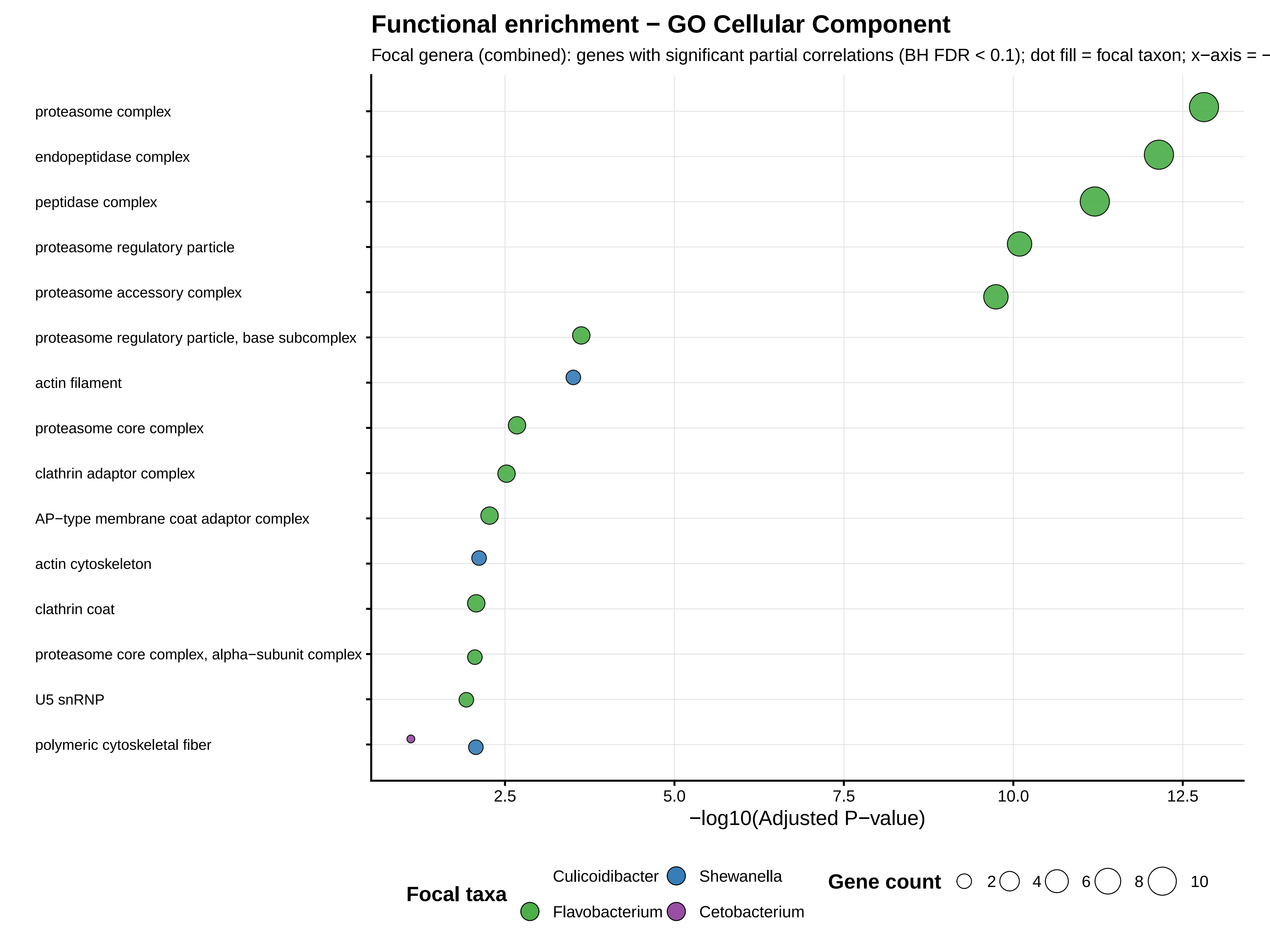
Functional enrichment − GO Cellular Component
Focal genera (combined): genes with significant partial correlations (BH FDR < 0.1); dot fill = focal taxon; x−axis = −
proteasome complex
endopeptidase complex
peptidase complex
proteasome regulatory particle
proteasome accessory complex
proteasome regulatory particle, base subcomplex
actin filament
proteasome core complex
clathrin adaptor complex
AP−type membrane coat adaptor complex
actin cytoskeleton
clathrin coat
proteasome core complex, alpha−subunit complex
U5 snRNP
polymeric cytoskeletal fiber
2.5
5.0
7.5
10.0
12.5
−log10(Adjusted P−value)
Focal taxa
Culicoidibacter
Shewanella
Flavobacterium
Cetobacterium
Gene count
2
4
6
8
10

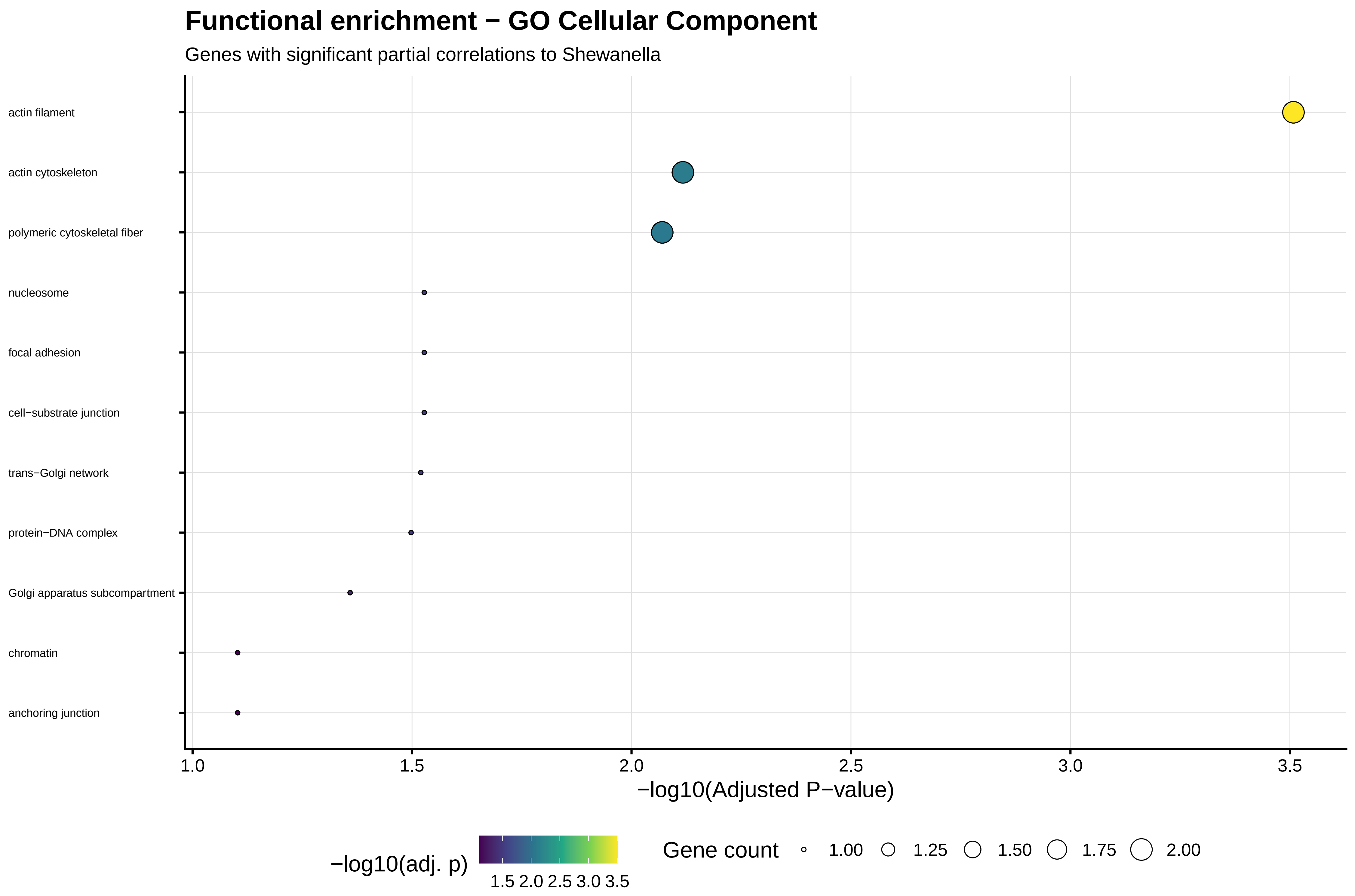
Functional enrichment − GO Cellular Component
Genes with significant partial correlations to Shewanella
actin filament
actin cytoskeleton
polymeric cytoskeletal fiber
nucleosome
focal adhesion
cell−substrate junction
trans−Golgi network
protein−DNA complex
Golgi apparatus subcompartment
chromatin
anchoring junction
1.0
1.5
2.0
2.5
3.0
3.5
−log10(Adjusted P−value)
−log10(adj. p)
1.5 2.0 2.5 3.0 3.5
Gene count
1.00
1.25
1.50
1.75
2.00

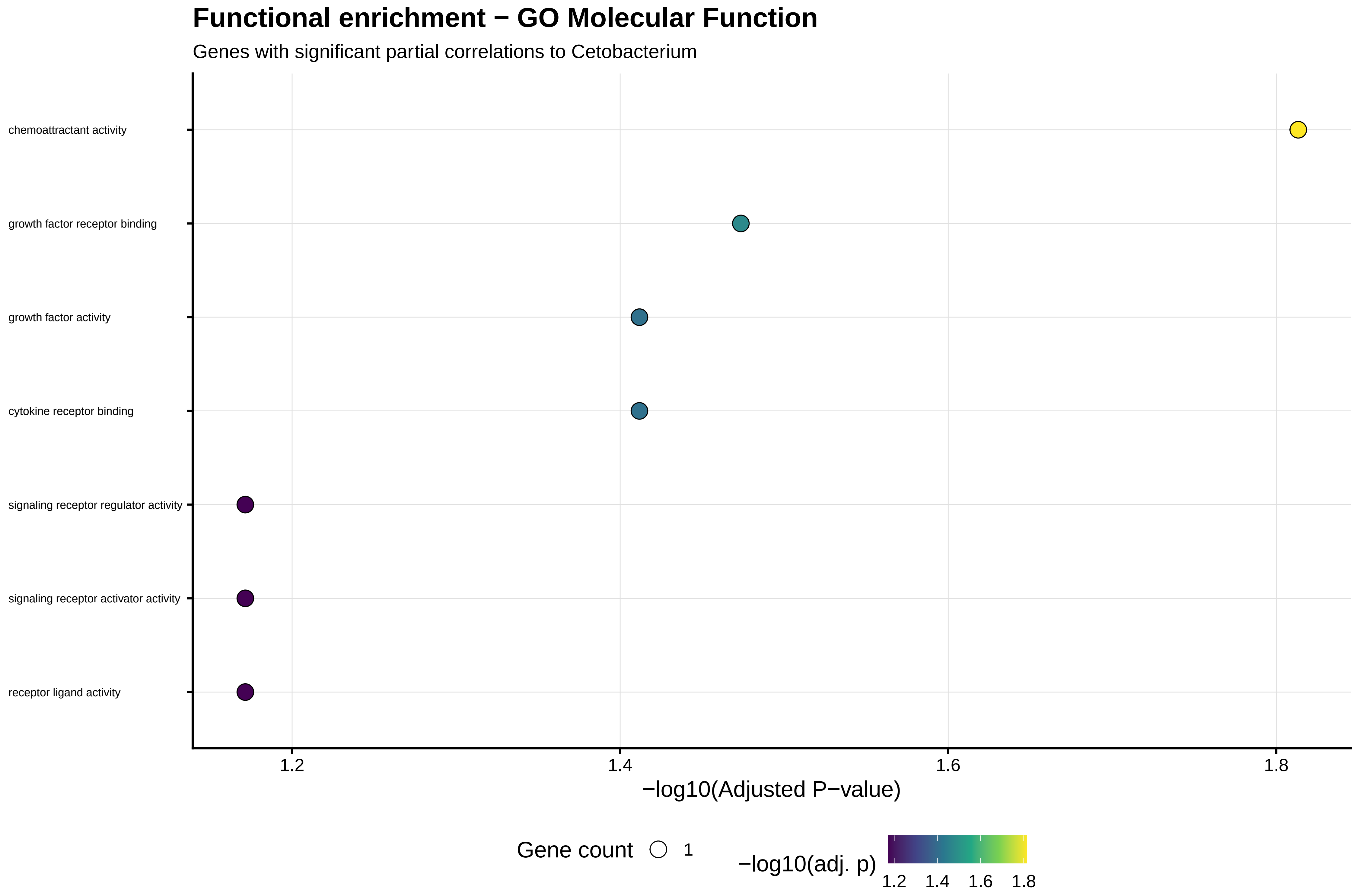
Functional enrichment − GO Molecular Function
Genes with significant partial correlations to Cetobacterium
chemoattractant activity
growth factor receptor binding
growth factor activity
cytokine receptor binding
signaling receptor regulator activity
signaling receptor activator activity
receptor ligand activity
1.2
1.4
1.6
1.8
−log10(Adjusted P−value)
Gene count
1
−log10(adj. p)
1.2
1.4
1.6
1.8

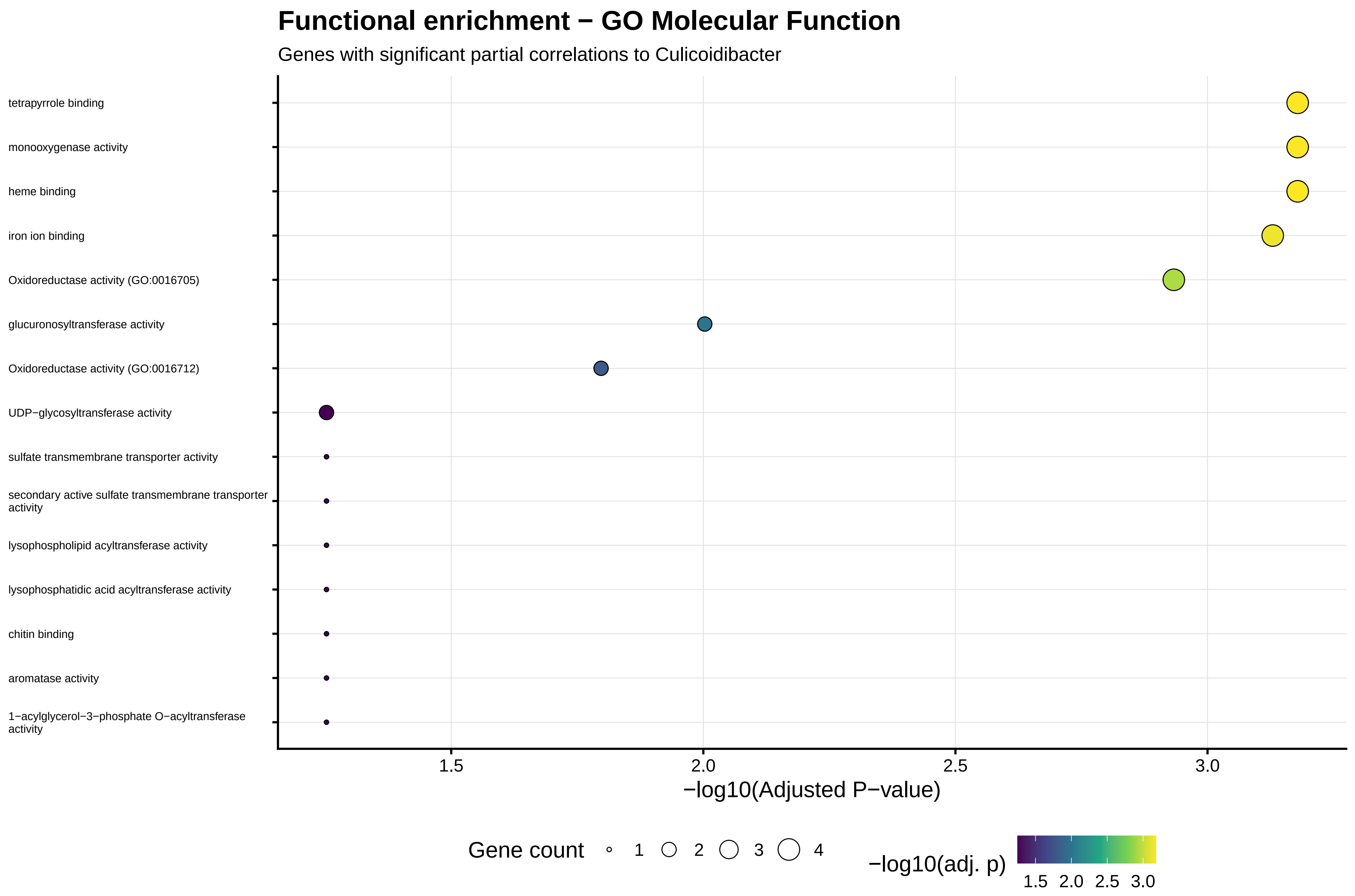

Functional enrichment − GO Molecular Function
Genes with significant partial correlations to Culicoidibacter
tetrapyrrole binding
monooxygenase activity
heme binding
iron ion binding
Oxidoreductase activity (GO:0016705)
glucuronosyltransferase activity
Oxidoreductase activity (GO:0016712)
UDP−glycosyltransferase activity
sulfate transmembrane transporter activity
secondary active sulfate transmembrane transporter activity
lysophospholipid acyltransferase activity
lysophosphatidic acid acyltransferase activity
chitin binding
aromatase activity
1−acylglycerol−3−phosphate O−acyltransferase activity
1.5
2.0
2.5
3.0
−log10(Adjusted P−value)
Gene count
1
2
3
4
−log10(adj. p)
1.5
2.0
2.5
3.0

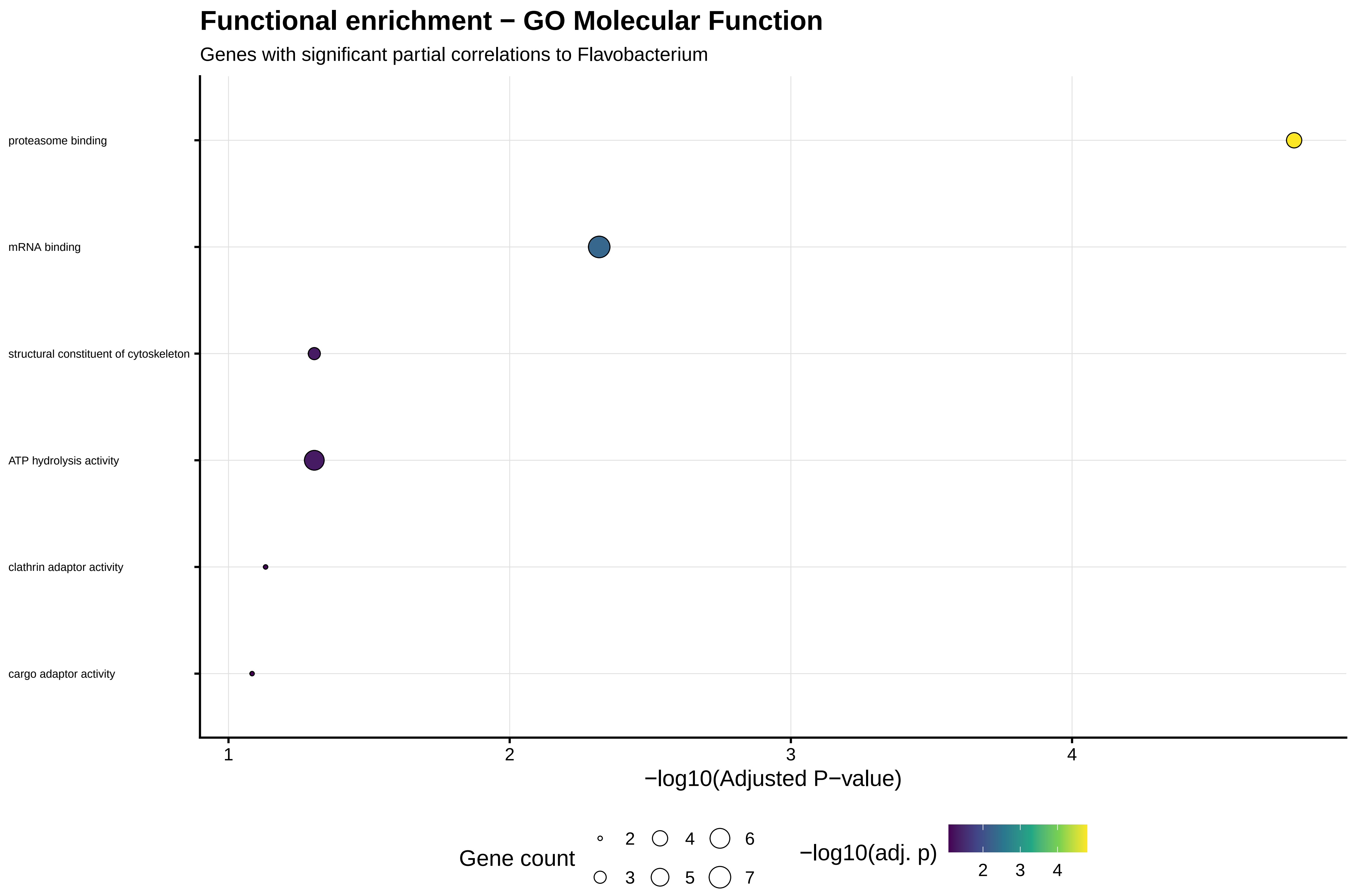

Functional enrichment − GO Molecular Function
Genes with significant partial correlations to Flavobacterium
proteasome binding
mRNA binding
structural constituent of cytoskeleton
ATP hydrolysis activity
clathrin adaptor activity
cargo adaptor activity
1
2
3
4
−log10(Adjusted P−value)
Gene count
2
3
4
5
6
7
−log10(adj. p)
2
3
4

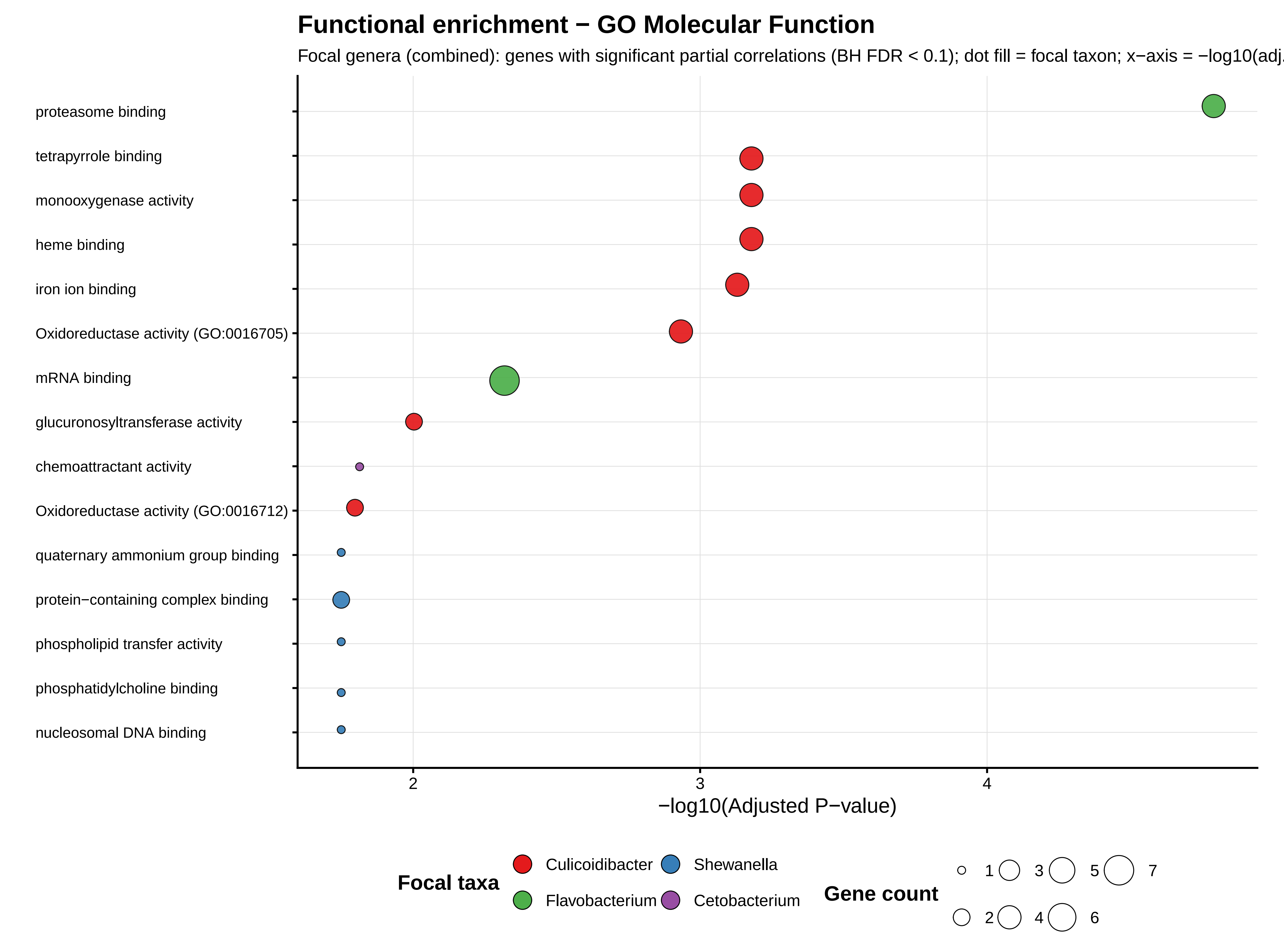

Functional enrichment − GO Molecular Function
Focal genera (combined): genes with significant partial correlations (BH FDR < 0.1); dot fill = focal taxon; x−axis = −log10(adj.
proteasome binding
tetrapyrrole binding
monooxygenase activity
heme binding
iron ion binding
Oxidoreductase activity (GO:0016705)
mRNA binding
glucuronosyltransferase activity
chemoattractant activity
Oxidoreductase activity (GO:0016712)
quaternary ammonium group binding
protein−containing complex binding
phospholipid transfer activity
phosphatidylcholine binding
nucleosomal DNA binding
2
3
4
−log10(Adjusted P−value)
Focal taxa
Culicoidibacter
Shewanella
Flavobacterium
Cetobacterium
Gene count
1
2
3
4
5
6
7

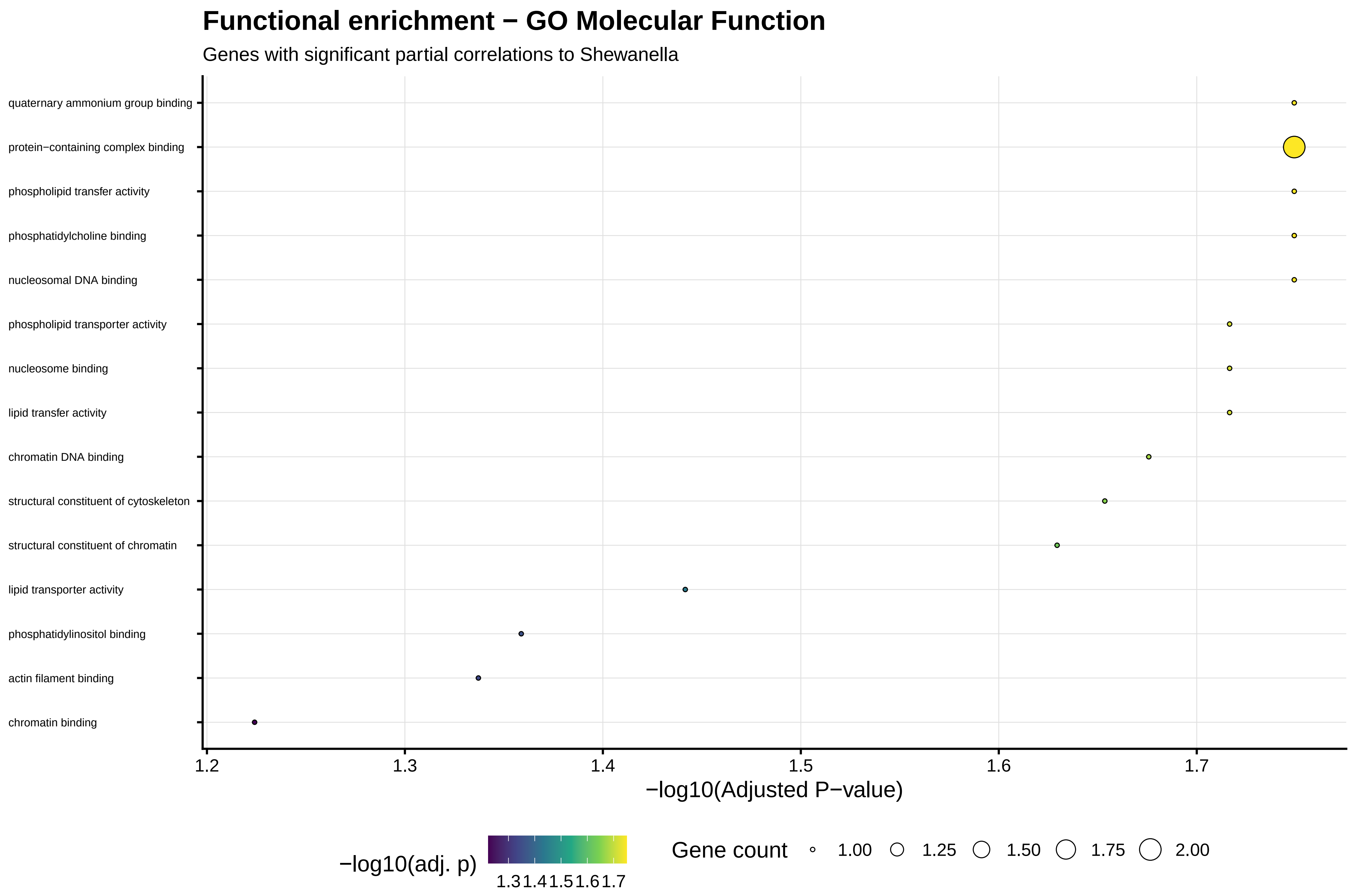
Functional enrichment − GO Molecular Function
Genes with significant partial correlations to Shewanella
quaternary ammonium group binding
protein−containing complex binding
phospholipid transfer activity
phosphatidylcholine binding
nucleosomal DNA binding
phospholipid transporter activity
nucleosome binding
lipid transfer activity
chromatin DNA binding
structural constituent of cytoskeleton
structural constituent of chromatin
lipid transporter activity
phosphatidylinositol binding
actin filament binding
chromatin binding
1.2
1.3
1.4
1.5
1.6
1.7
−log10(Adjusted P−value)
−log10(adj. p)
1.3 1.4 1.5 1.6 1.7
Gene count
1.00
1.25
1.50
1.75
2.00

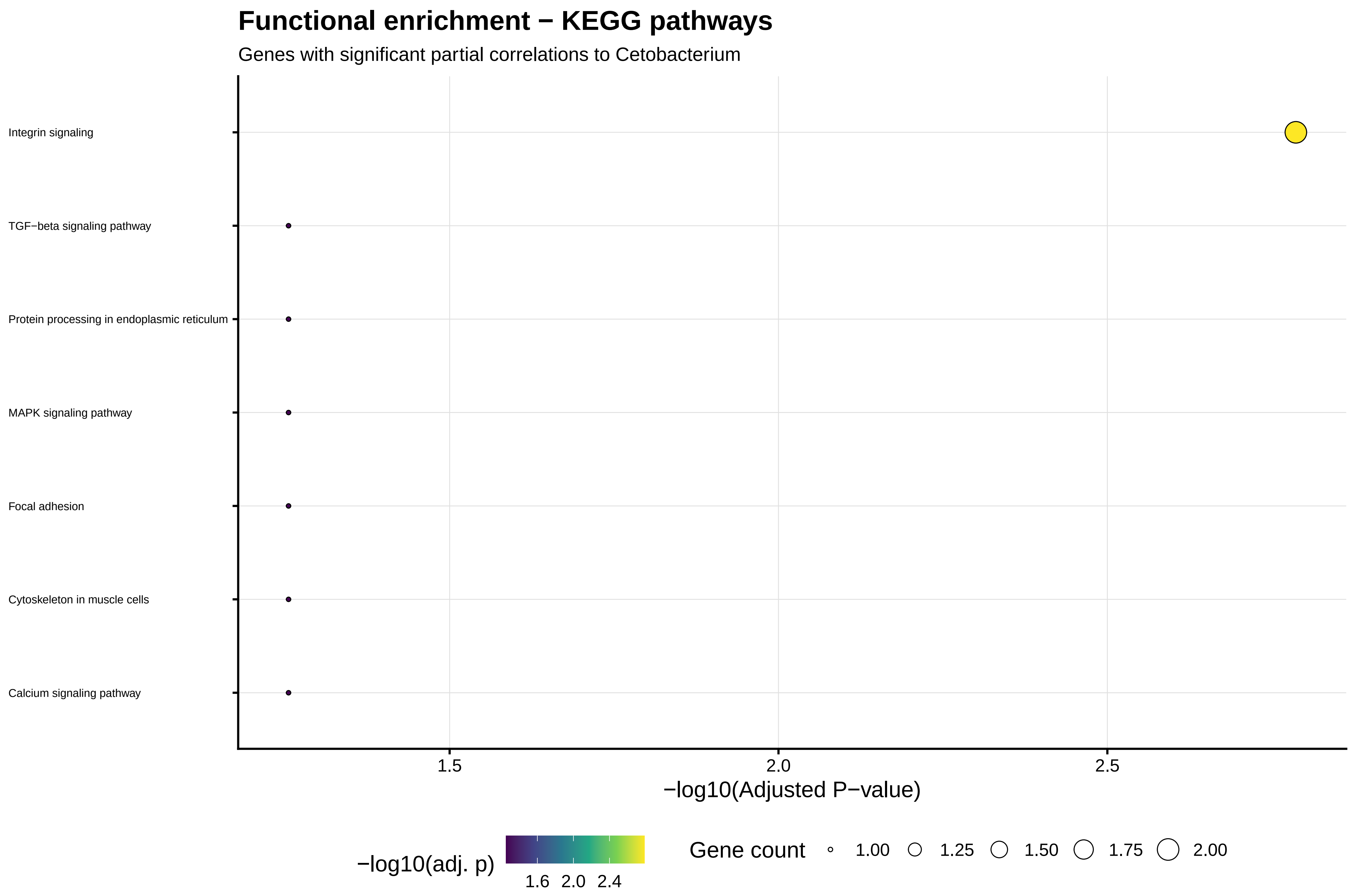

Functional enrichment − KEGG pathways
Genes with significant partial correlations to Cetobacterium
Integrin signaling
TGF−beta signaling pathway
Protein processing in endoplasmic reticulum
MAPK signaling pathway
Focal adhesion
Cytoskeleton in muscle cells
Calcium signaling pathway
1.5
2.0
2.5
−log10(Adjusted P−value)
−log10(adj. p)
1.6
2.0
2.4
Gene count
1.00
1.25
1.50
1.75
2.00

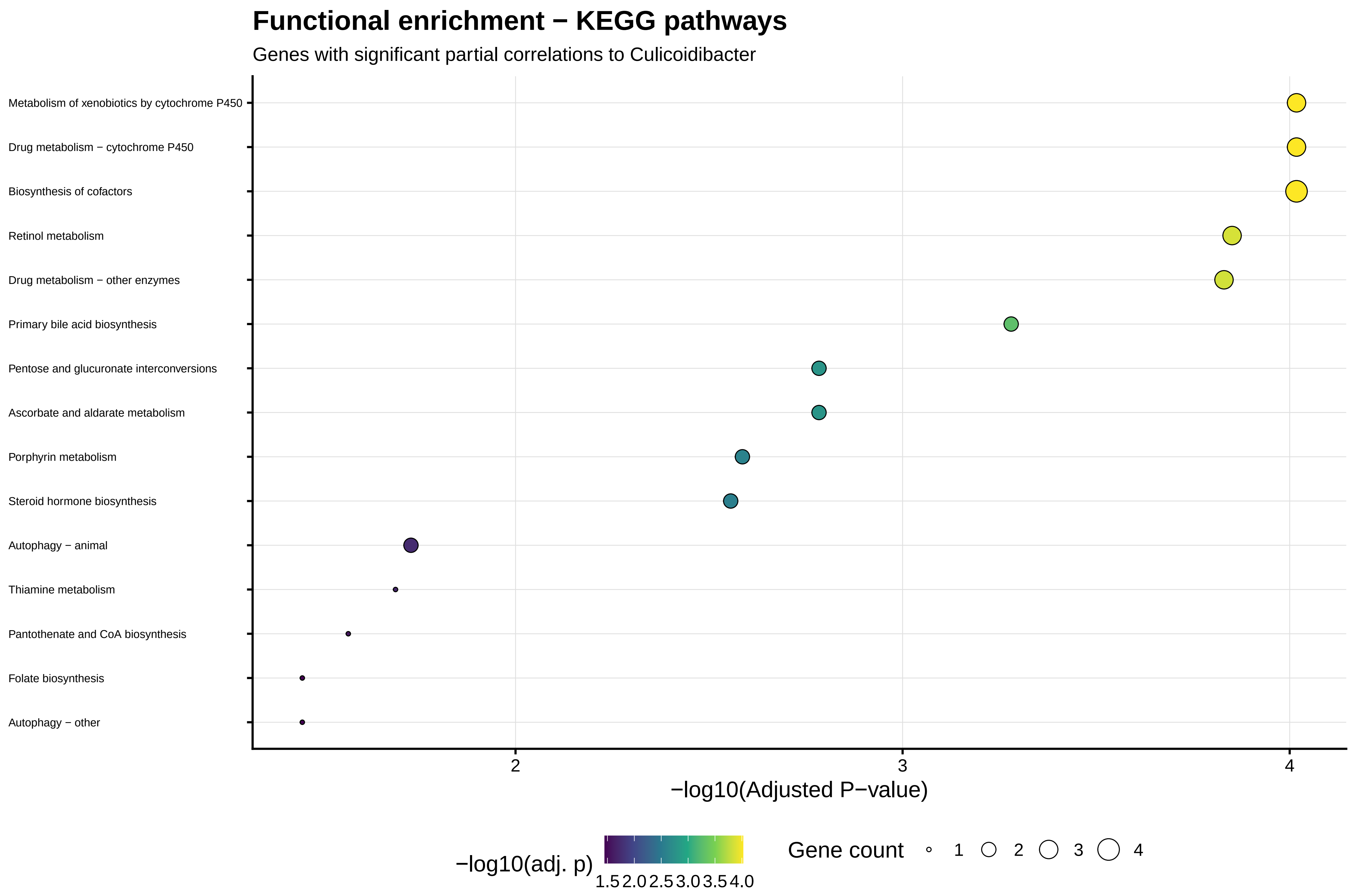

Functional enrichment − KEGG pathways
Genes with significant partial correlations to Culicoidibacter
Metabolism of xenobiotics by cytochrome P450
Drug metabolism − cytochrome P450
Biosynthesis of cofactors
Retinol metabolism
Drug metabolism − other enzymes
Primary bile acid biosynthesis
Pentose and glucuronate interconversions
Ascorbate and aldarate metabolism
Porphyrin metabolism
Steroid hormone biosynthesis
Autophagy − animal
Thiamine metabolism
Pantothenate and CoA biosynthesis
Folate biosynthesis
Autophagy − other
2
3
4
−log10(Adjusted P−value)
−log10(adj. p)
1.5 2.0 2.5 3.0 3.5 4.0
Gene count
1
2
3
4

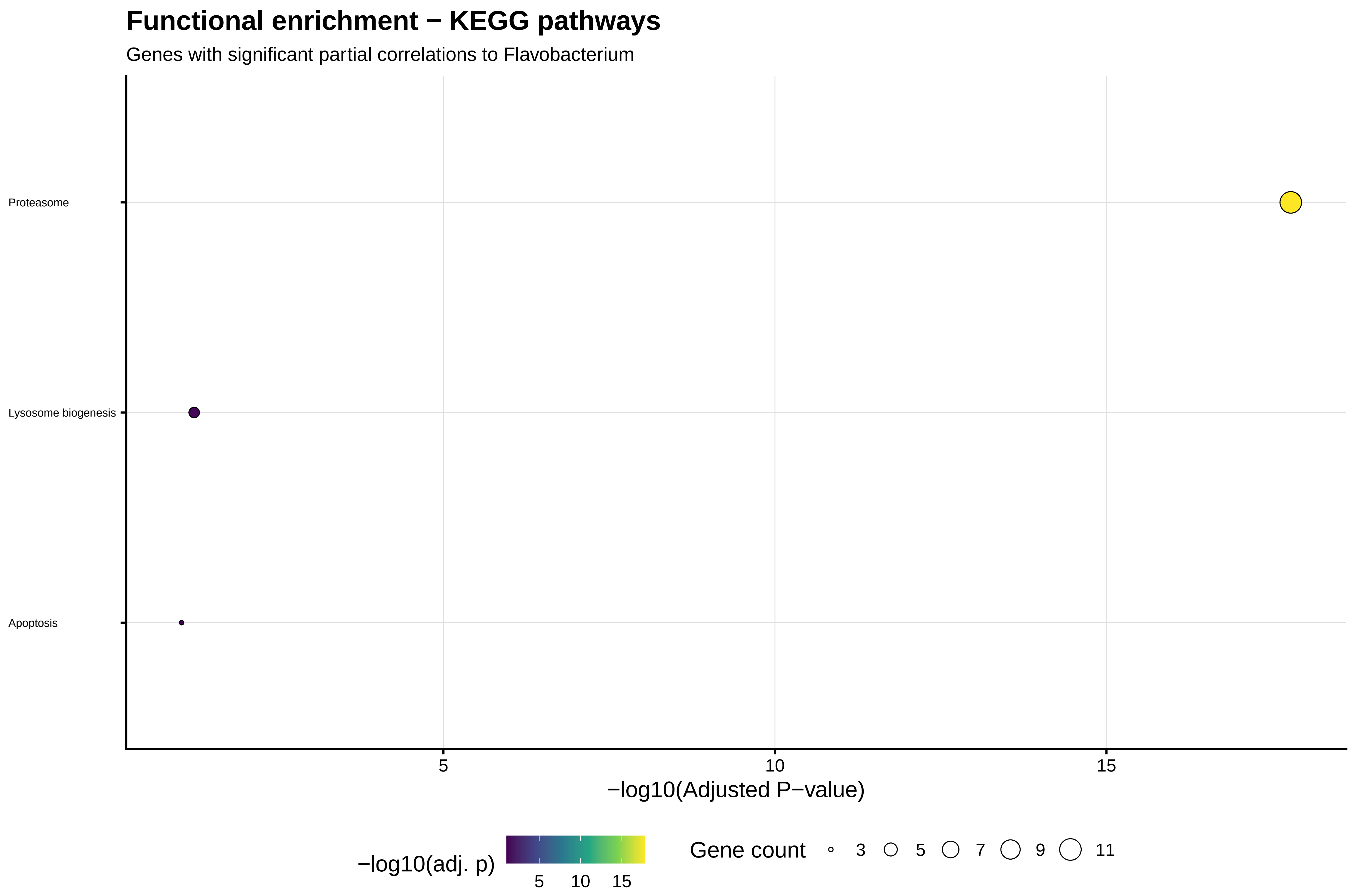

Functional enrichment − KEGG pathways
Genes with significant partial correlations to Flavobacterium
Proteasome
Lysosome biogenesis
Apoptosis
5
10
15
−log10(Adjusted P−value)
−log10(adj. p)
5
10
15
Gene count
3
5
7
9
11

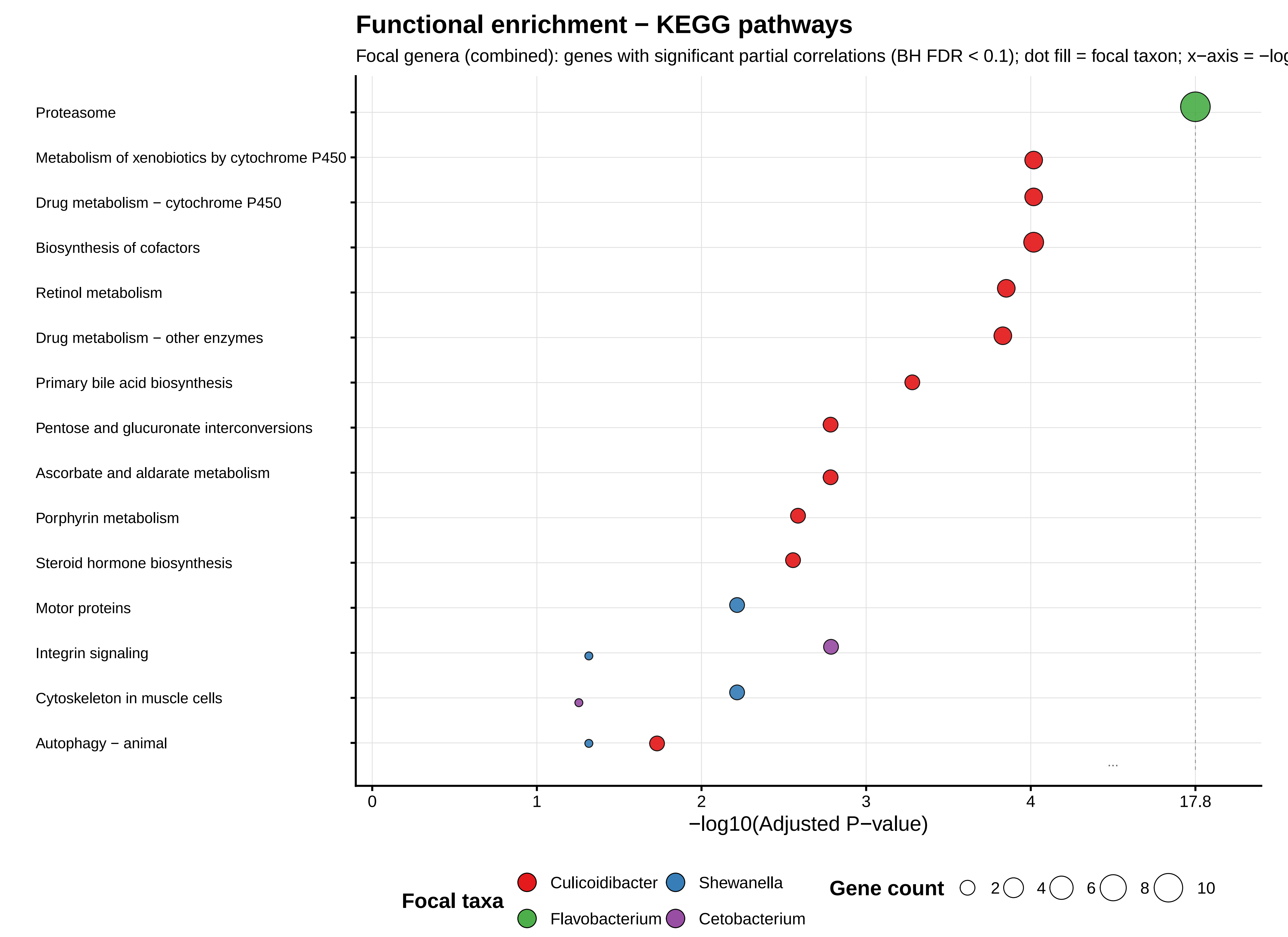
Functional enrichment − KEGG pathways
Focal genera (combined): genes with significant partial correlations (BH FDR < 0.1); dot fill = focal taxon; x−axis = −log
Proteasome
Metabolism of xenobiotics by cytochrome P450
Drug metabolism − cytochrome P450
Biosynthesis of cofactors
Retinol metabolism
Drug metabolism − other enzymes
Primary bile acid biosynthesis
Pentose and glucuronate interconversions
Ascorbate and aldarate metabolism
Porphyrin metabolism
Steroid hormone biosynthesis
Motor proteins
Integrin signaling
Cytoskeleton in muscle cells
Autophagy − animal
...
0
1
2
3
4
17.8
−log10(Adjusted P−value)
Focal taxa
Culicoidibacter
Shewanella
Flavobacterium
Cetobacterium
Gene count
2
4
6
8
10

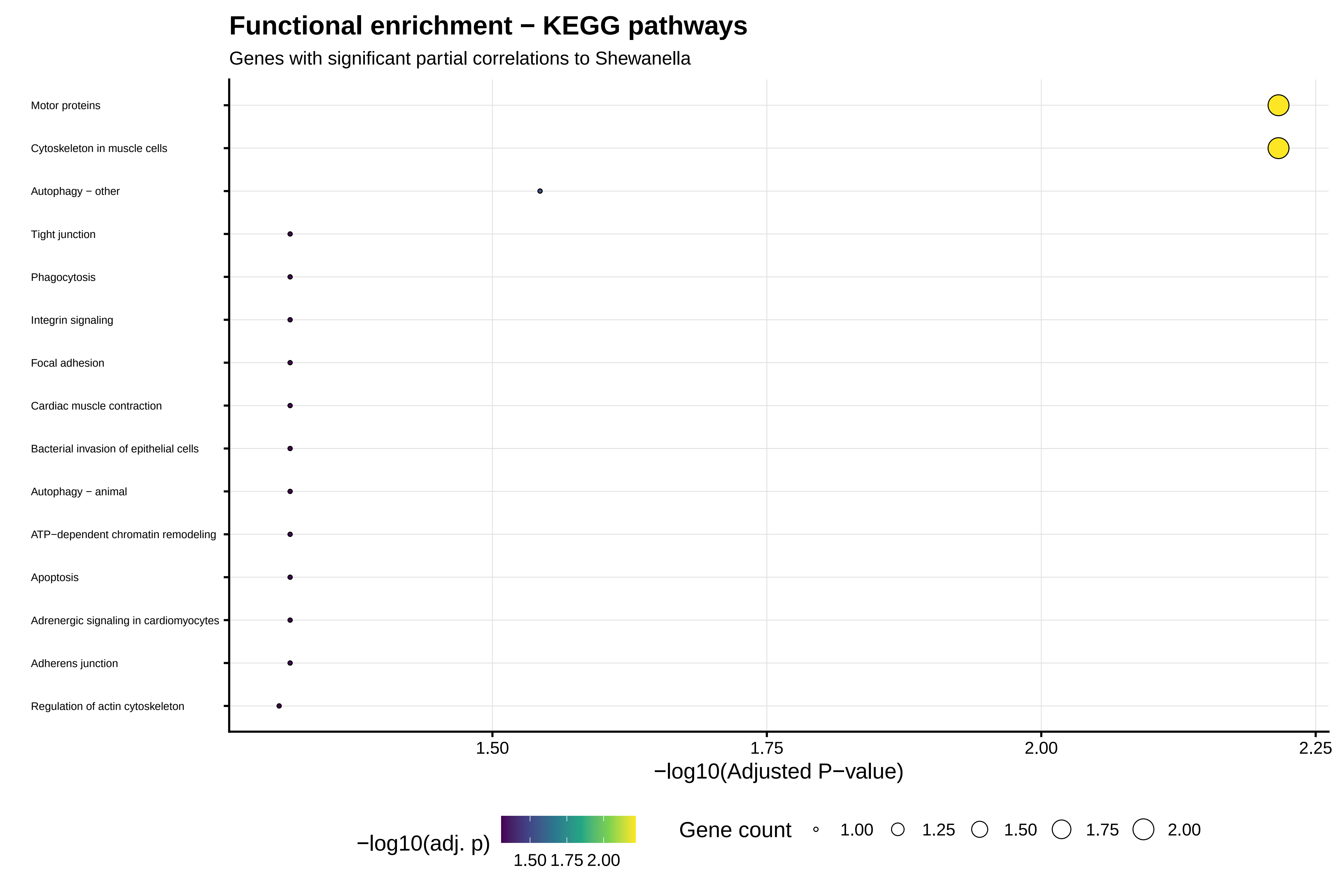

Functional enrichment − KEGG pathways
Genes with significant partial correlations to Shewanella
Motor proteins
Cytoskeleton in muscle cells
Autophagy − other
Tight junction
Phagocytosis
Integrin signaling
Focal adhesion
Cardiac muscle contraction
Bacterial invasion of epithelial cells
Autophagy − animal
ATP−dependent chromatin remodeling
Apoptosis
Adrenergic signaling in cardiomyocytes
Adherens junction
Regulation of actin cytoskeleton
1.50
1.75
2.00
2.25
−log10(Adjusted P−value)
−log10(adj. p)
1.50 1.75 2.00
Gene count
1.00
1.25
1.50
1.75
2.00
