## Supplementary Figures for "Historical contingency shapes zebrafish host-microbiome responses to a subsequent biotic challenge": Supplementary_Figures__08__NeutralModel__submission__2026-06-26.pdf

### Supplementary figures (combined)

Build (UTC): 2026−06−26 12:36:51 UTC

Git: 55d6b9a4a7bf134b3db15f0777c389e2da8c31be

Module: 08__NeutralModel

Figures dir: /Sieler2026/Results/08__NeutralModel/Figures

Figures (combined PDF order; p. = first page of that figure in this PDF):

p. 2: Figure S8.10.1 − neutral_model_Time60__Genus__focal_four_genera

p. 3: Figure S8.10.2 − neutral_model_Time60__ASV

p. 4: Figure S8.10.3 − neutral_model_Time60__ASV__cetobacterium

p. 5: Figure S8.10.4 − neutral_model_Time60__ASV__culicoidibacter

p. 6: Figure S8.10.5 − neutral_model_Time60__ASV__flavobacterium

p. 7: Figure S8.10.6 − neutral_model_Time60__ASV__shewanella

p. 8: Figure S8.10.7 − neutral_model_Time60__Genus

p. 9: Figure S8.10.8 − neutral_model_Time60__Genus__cetobacterium

p. 10: Figure S8.10.9 − neutral_model_Time60__Genus__culicoidibacter

p. 11: Figure S8.10.10 − neutral_model_Time60__Genus__flavobacterium

p. 12: Figure S8.10.11 − neutral_model_Time60__Genus__shewanella

p. 13: Figure S8.10.12 − neutral_model_Time60__ASV__focal_two_cetobacterium

p. 14: Figure S8.10.13 − neutral_model_Time60__ASV__focal_two_Culici

p. 15: Figure S8.10.14 − neutral_model_Time60__ASV__focal_two_flavobacterium

p. 16: Figure S8.10.15 − neutral_model_Time60__ASV__focal_two_shewanella

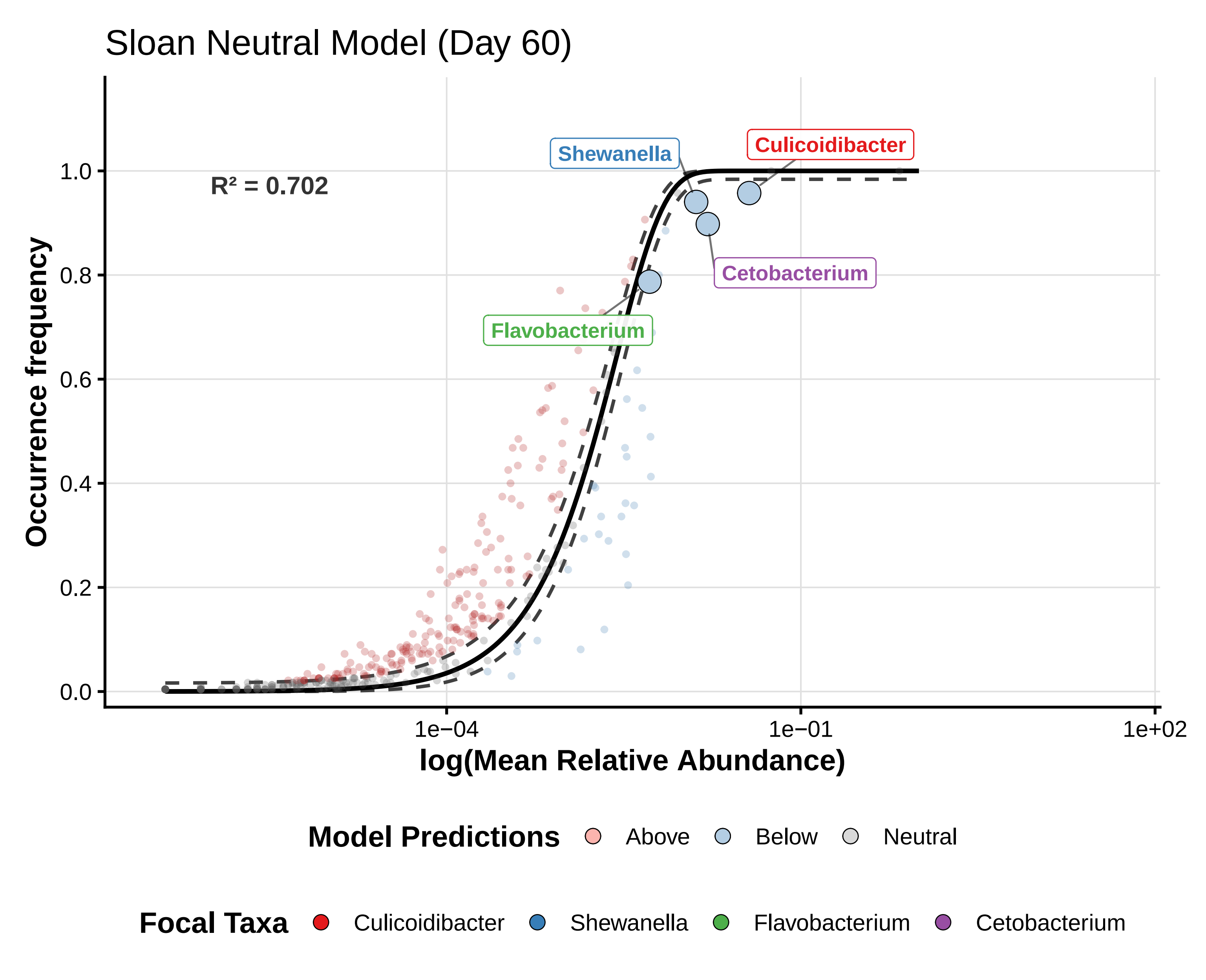

Sloan Neutral Model (Day 60)
R² = 0.702
Shewanella
Culicoidibacter
Cetobacterium
Flavobacterium
Occurrence frequency
0.0
0.2
0.4
0.6
0.8
1.0
1e−04
1e−01
1e+02
log(Mean Relative Abundance)
Model Predictions
Above
Below
Neutral
Focal Taxa
Culicoidibacter
Shewanella
Flavobacterium
Cetobacterium

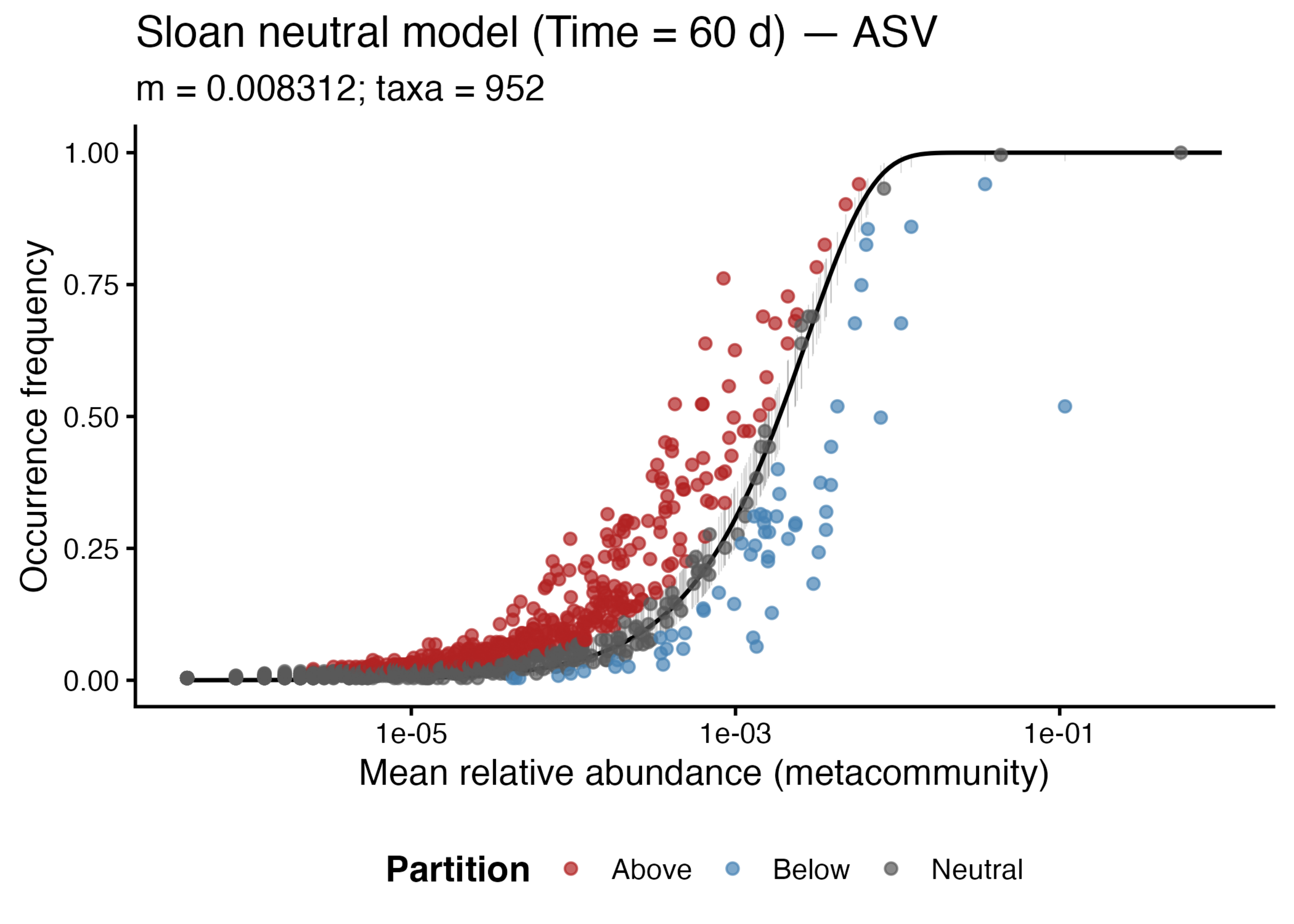

Sloan neutral model (Time = 60 d) — ASV
m = 0.008312; taxa = 952
Occurrence frequency
1.00
0.75
0.50
0.25
0.00
1e-05
1e-03
1e-01
Mean relative abundance (metacommunity)
Partition
Above
Below
Neutral

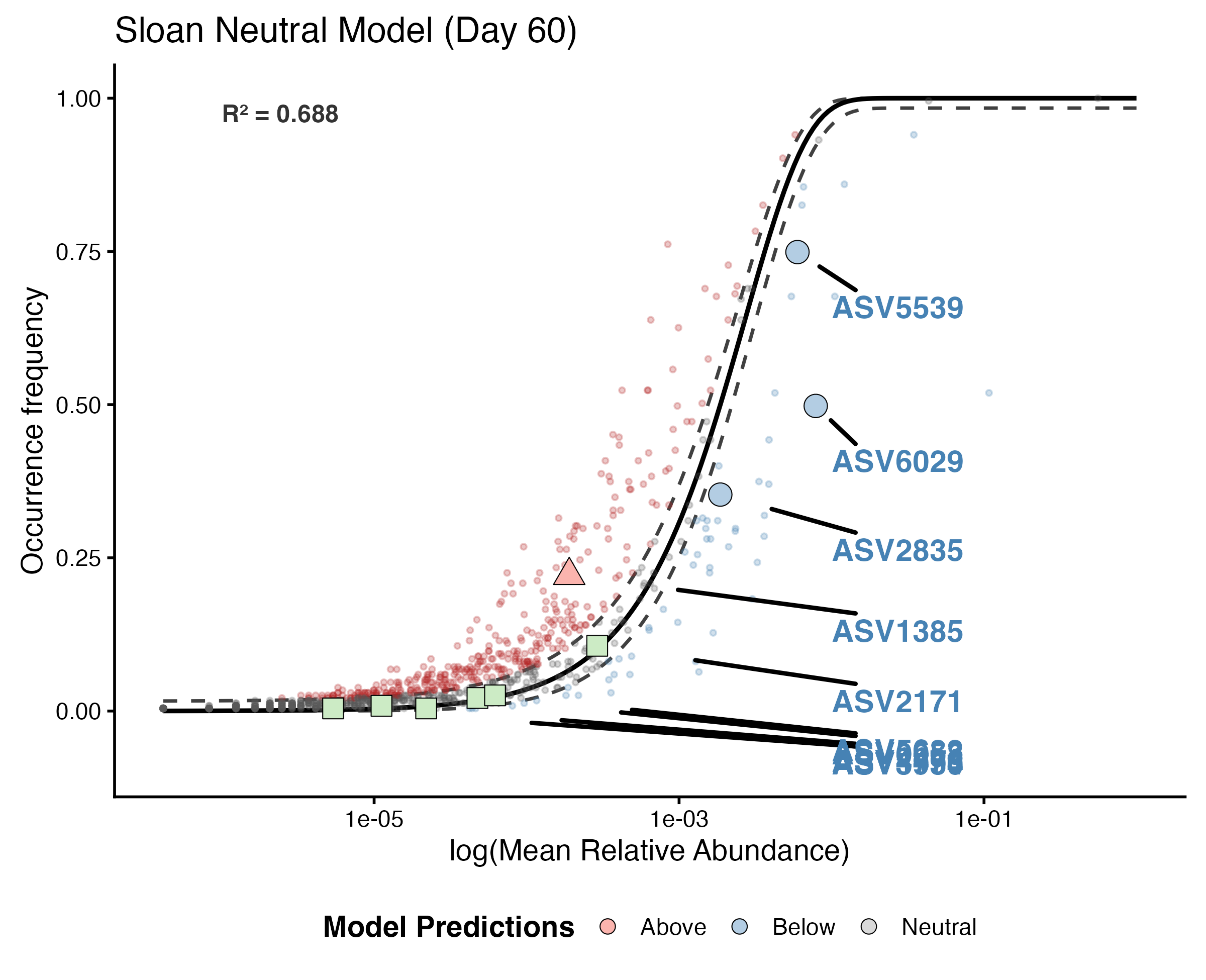
Sloan Neutral Model (Day 60)
$R^2$ = 0.688
1.00
0.75
0.50
0.25
0.00
Occurrence frequency
1e-05
1e-03
1e-01
log(Mean Relative Abundance)
ASV5539
ASV6029
ASV2835
ASV1385
ASV2171
Model Predictions
Above
Below
Neutral

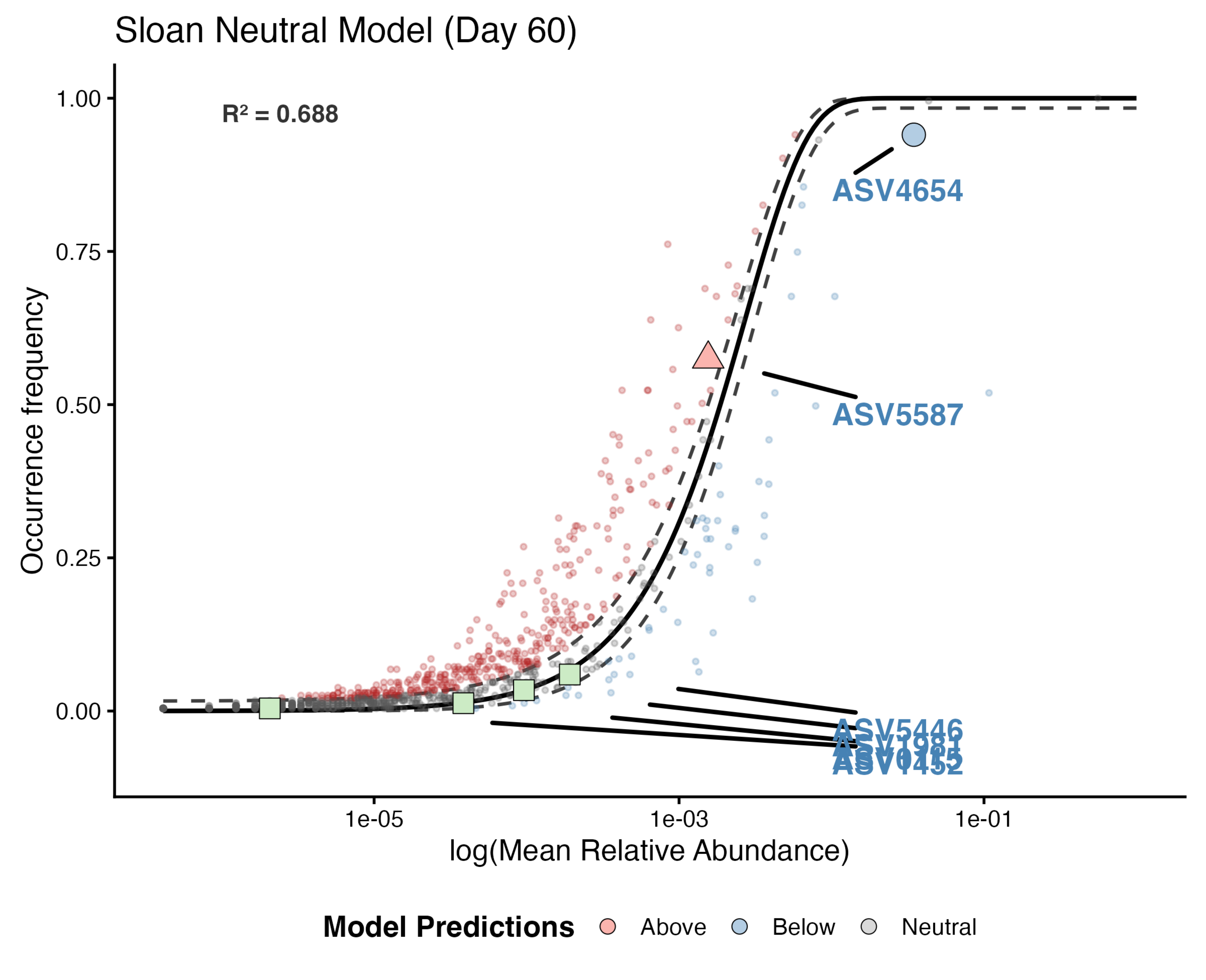
Sloan Neutral Model (Day 60)
R² = 0.688
1.00
0.75
0.50
0.25
0.00
Occurrence frequency
1e-05
1e-03
1e-01
log(Mean Relative Abundance)
ASV4654
ASV5587
ASV5446
Model Predictions
Above
Below
Neutral

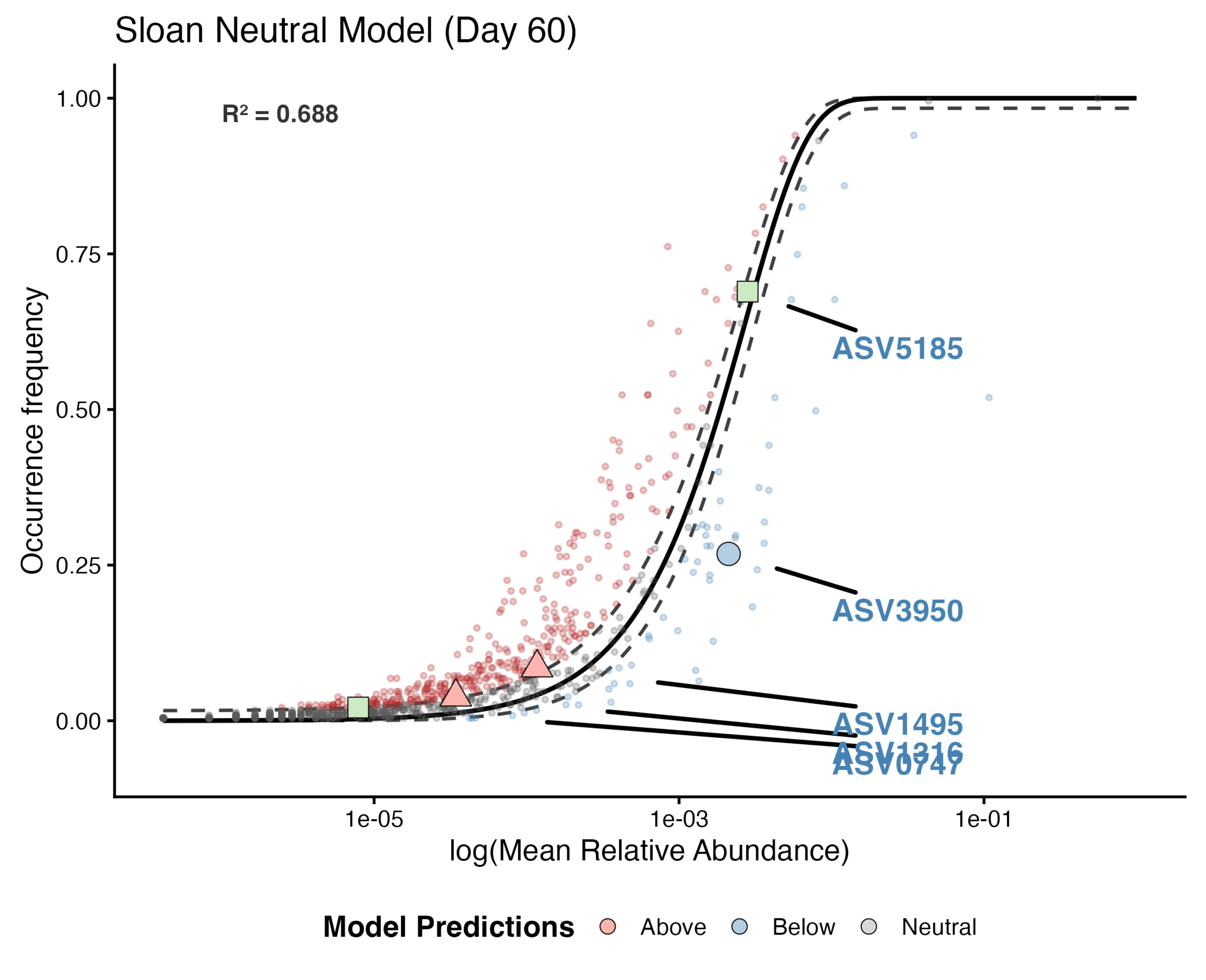

Sloan Neutral Model (Day 60)
$R^2 = 0.688$
Occurrence frequency
1.00
0.75
0.50
0.25
0.00
ASV5185
ASV3950
ASV1495
ASV1316
ASV0747
1e-05
1e-03
1e-01
log(Mean Relative Abundance)
Model Predictions
Above
Below
Neutral

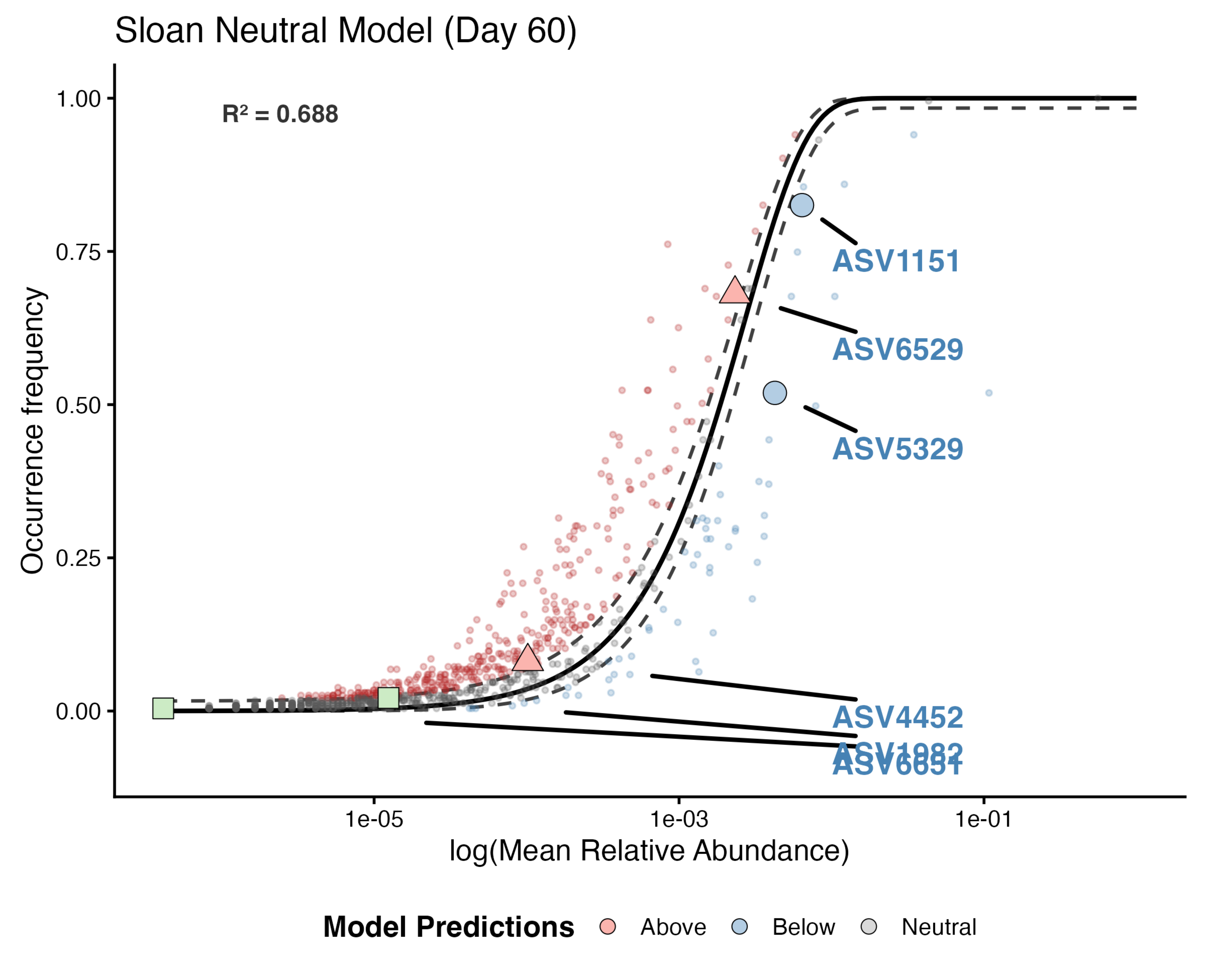

Sloan Neutral Model (Day 60)
R² = 0.688
1.00
0.75
0.50
0.25
0.00
Occurrence frequency
ASV1151
ASV6529
ASV5329
ASV4452
ASV1982
ASV6651
1e-05
1e-03
1e-01
log(Mean Relative Abundance)
Model Predictions
Above
Below
Neutral

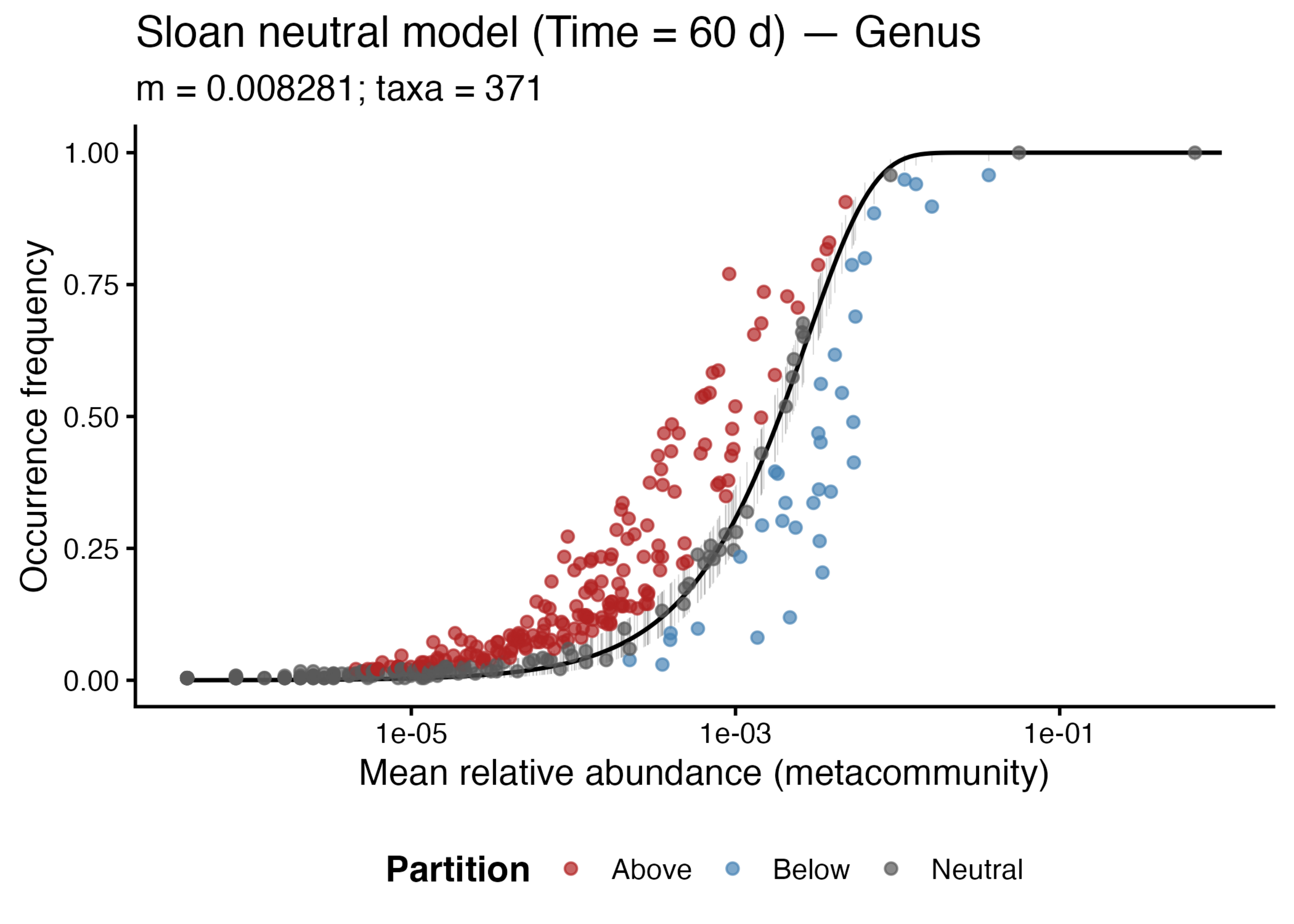

Sloan neutral model (Time = 60 d) — Genus
m = 0.008281; taxa = 371
Occurrence frequency
1.00
0.75
0.50
0.25
0.00
1e-05
1e-03
1e-01
Mean relative abundance (metacommunity)
Partition
Above
Below
Neutral

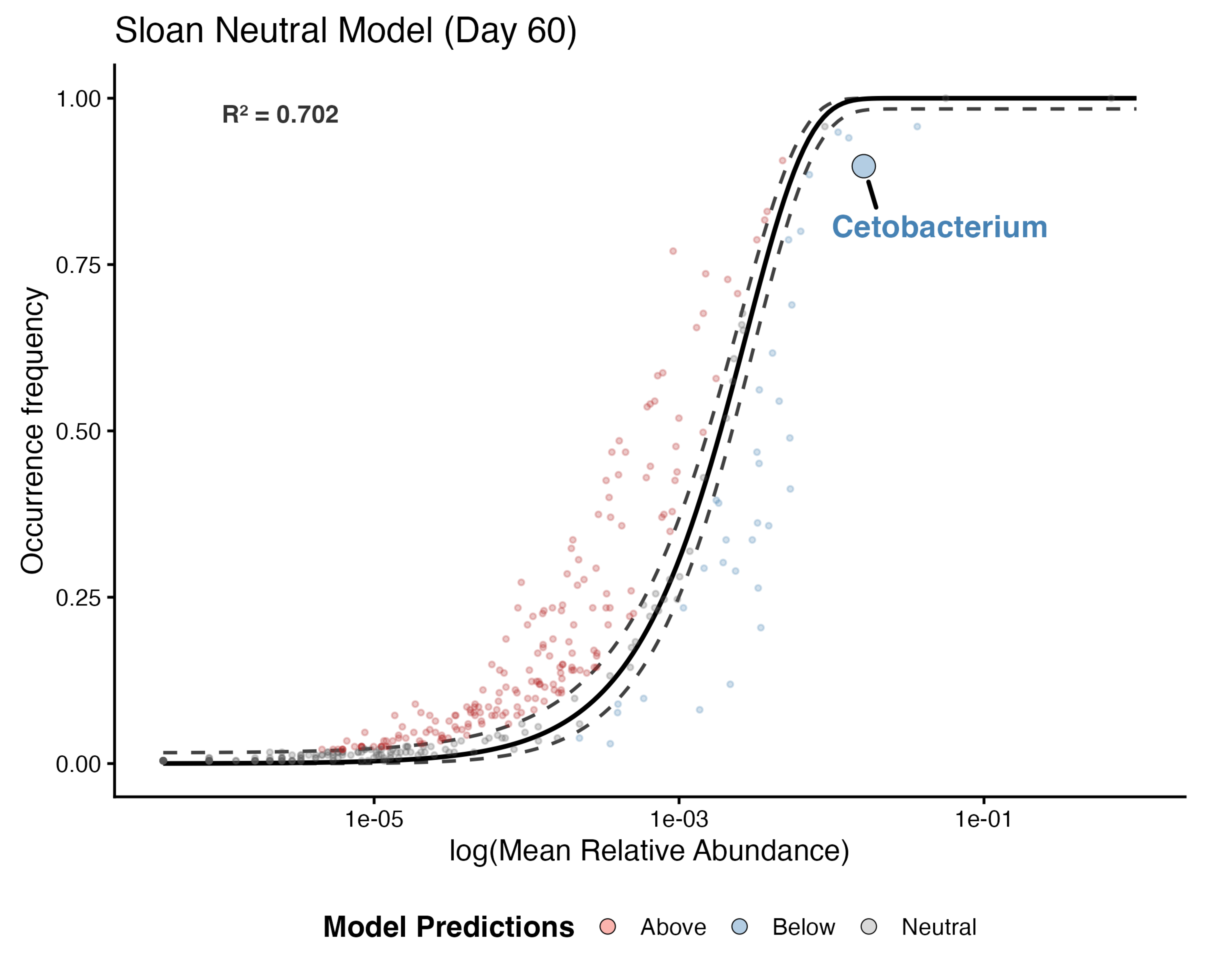

Sloan Neutral Model (Day 60)
R² = 0.702
Cetobacterium
Occurrence frequency
1.00
0.75
0.50
0.25
0.00
1e-05
1e-03
1e-01
log(Mean Relative Abundance)
Model Predictions
Above
Below
Neutral

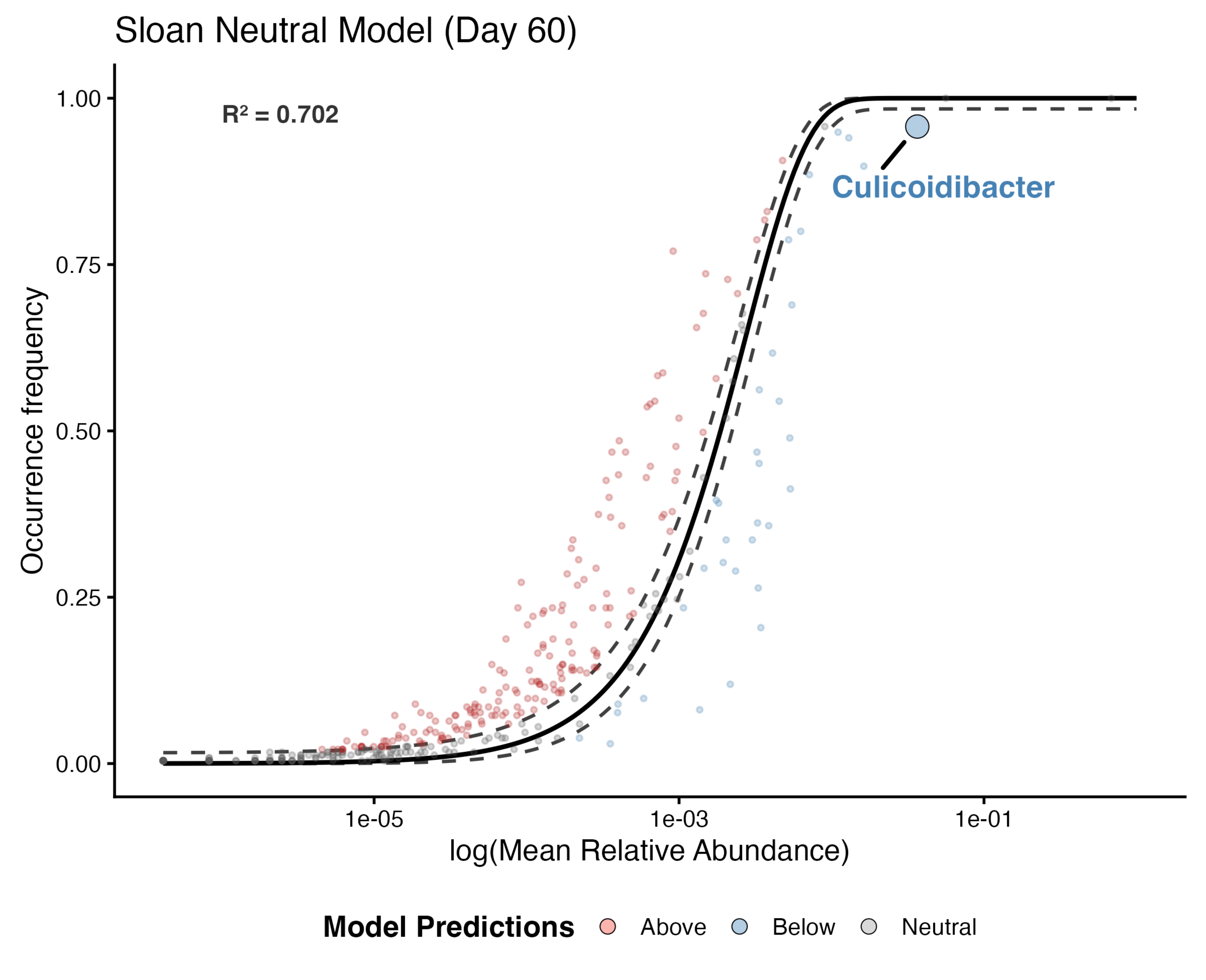

Sloan Neutral Model (Day 60)
R² = 0.702
Culicoidibacter
1.00
0.75
0.50
0.25
0.00
Occurrence frequency
1e-05
1e-03
1e-01
log(Mean Relative Abundance)
Model Predictions
Above
Below
Neutral

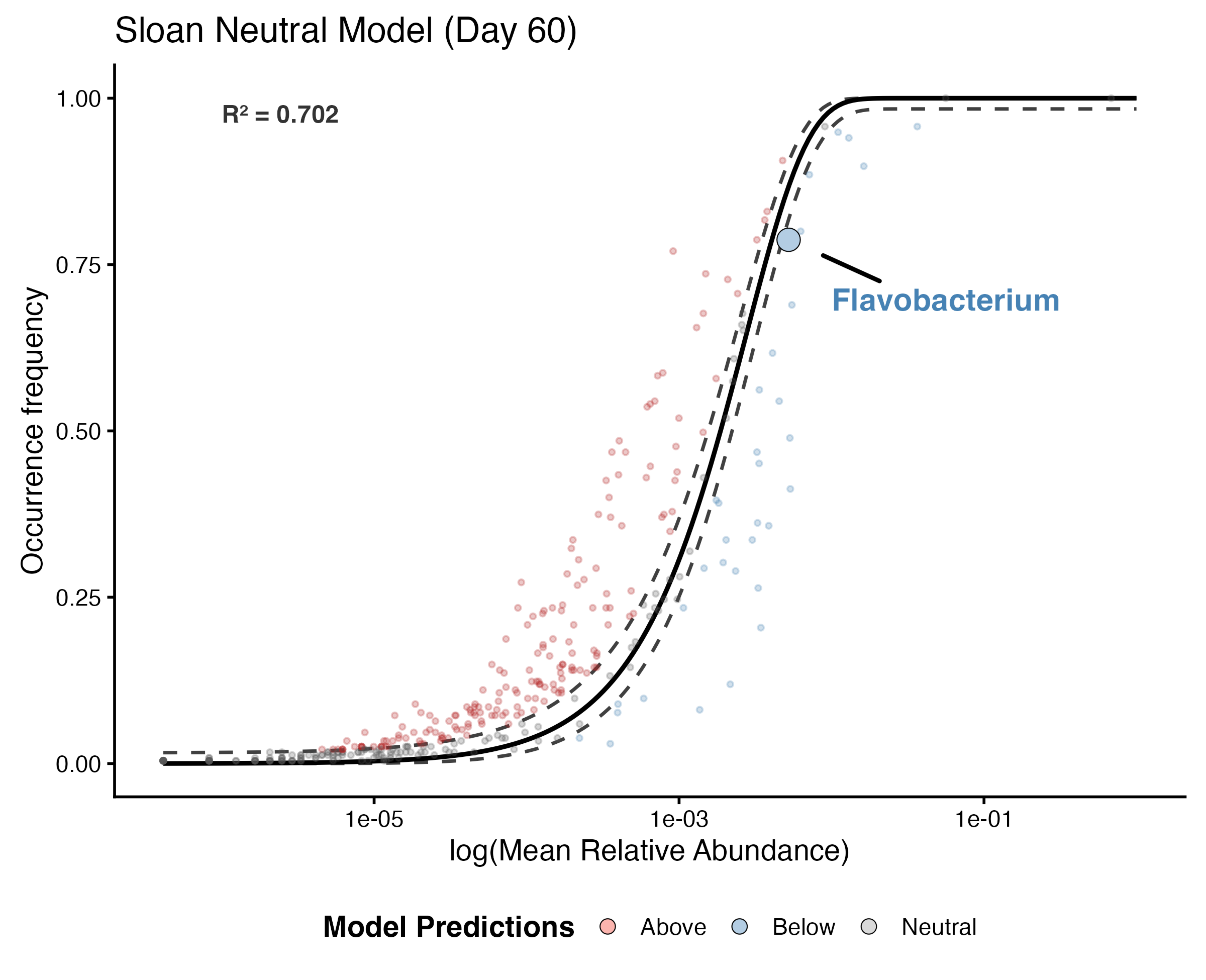

Sloan Neutral Model (Day 60)
R² = 0.702
Flavobacterium
1.00
0.75
0.50
0.25
0.00
Occurrence frequency
1e-05
1e-03
1e-01
log(Mean Relative Abundance)
Model Predictions
Above
Below
Neutral

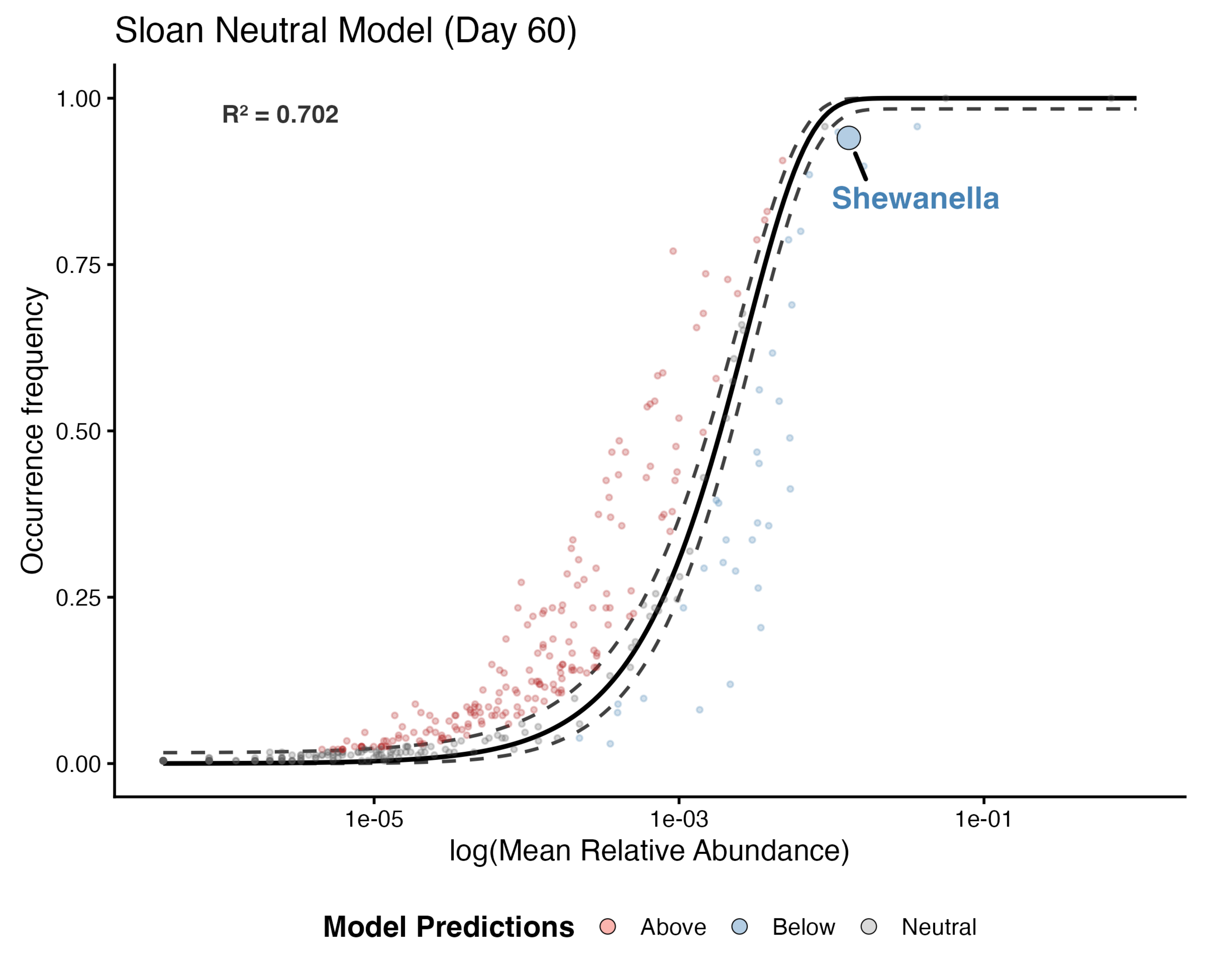
Sloan Neutral Model (Day 60)
R² = 0.702
Shewanella
Occurrence frequency
1.00
0.75
0.50
0.25
0.00
1e-05
1e-03
1e-01
log(Mean Relative Abundance)
Model Predictions
Above
Below
Neutral

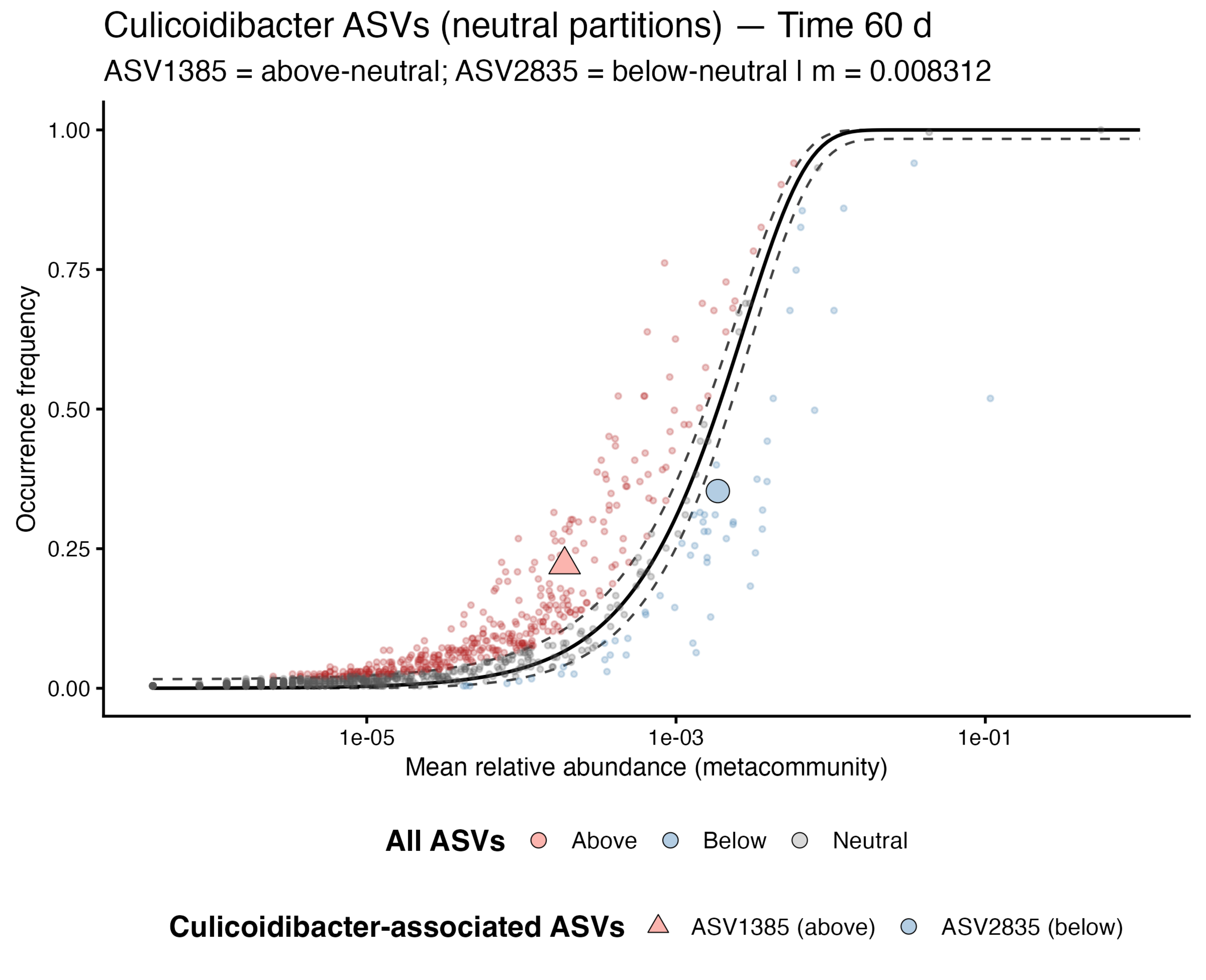

Culicoidibacter ASVs (neutral partitions) — Time 60 d
ASV1385 = above-neutral; ASV2835 = below-neutral | m = 0.008312
Occurrence frequency
0.00
0.25
0.50
0.75
1.00
1e-05
1e-03
1e-01
Mean relative abundance (metacommunity)
All ASVs
Above
Below
Neutral
Culicoidibacter-associated ASVs
ASV1385 (above)
ASV2835 (below)

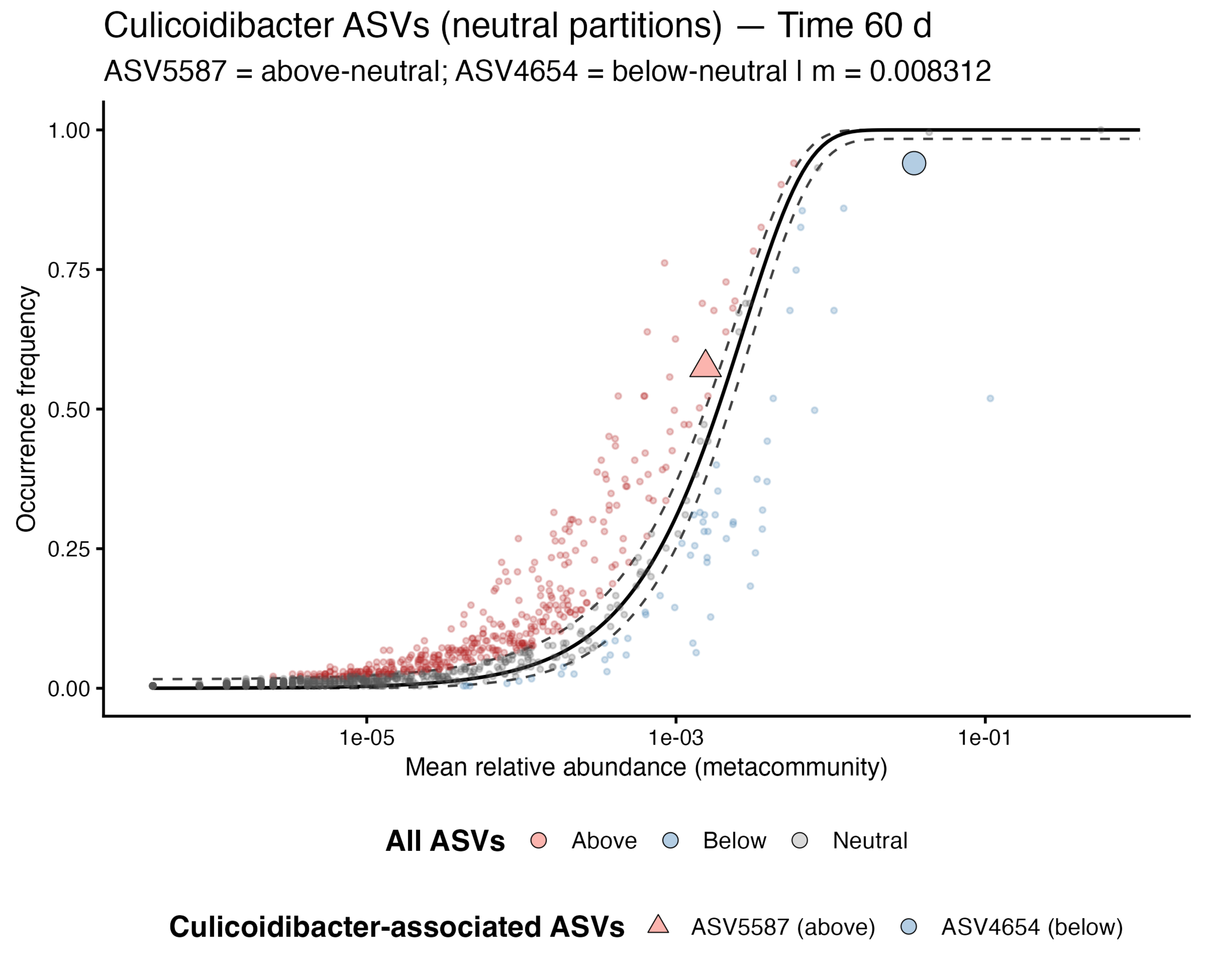
Culicoidibacter ASVs (neutral partitions) — Time 60 d
ASV5587 = above-neutral; ASV4654 = below-neutral | m = 0.008312
Occurrence frequency
1.00
0.75
0.50
0.25
0.00
1e-05
1e-03
1e-01
Mean relative abundance (metacommunity)
All ASVs
Above
Below
Neutral
Culicoidibacter-associated ASVs
ASV5587 (above)
ASV4654 (below)

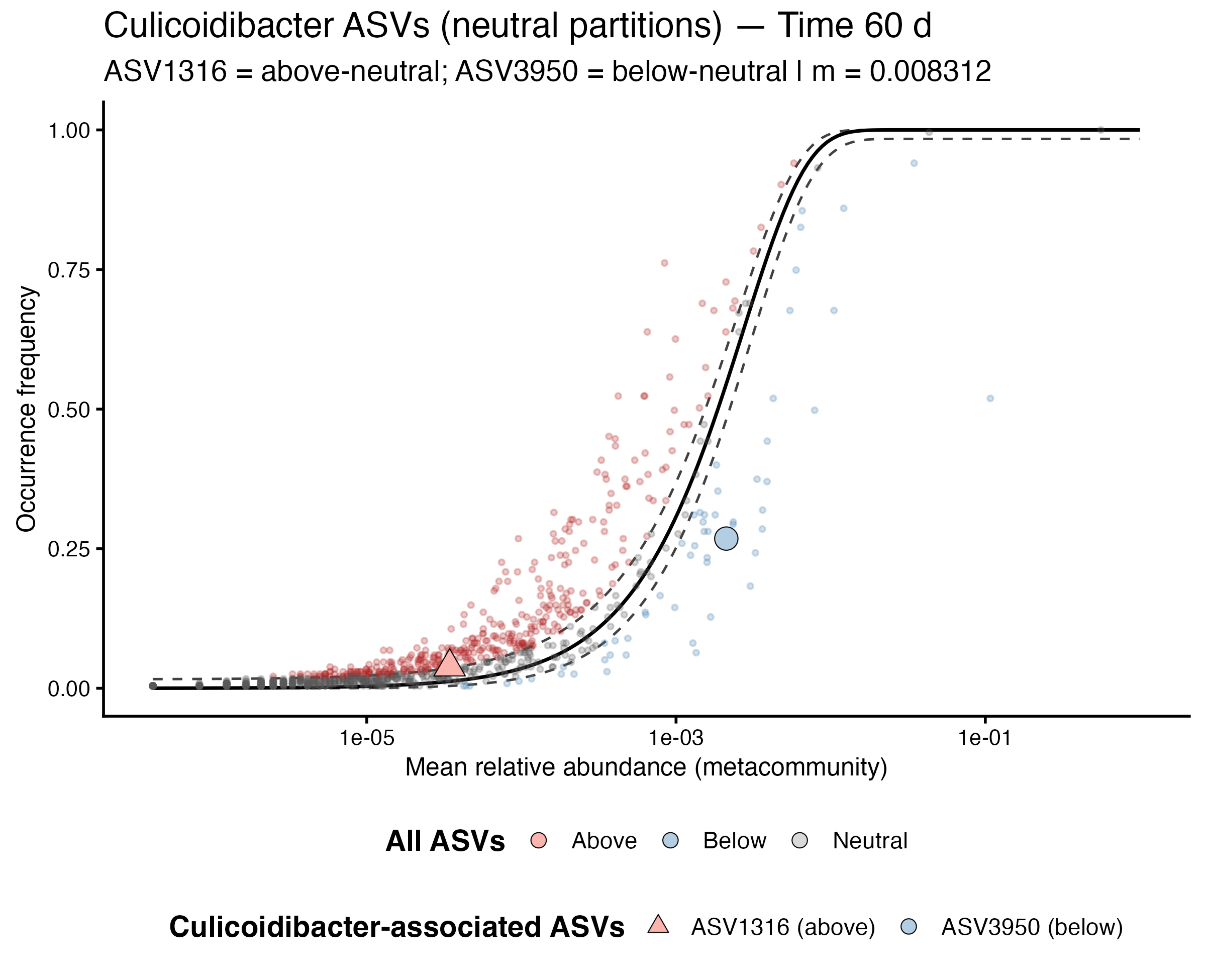
Culicoidibacter ASVs (neutral partitions) — Time 60 d
ASV1316 = above-neutral; ASV3950 = below-neutral | m = 0.008312
Occurrence frequency
1.00
0.75
0.50
0.25
0.00
1e-05
1e-03
1e-01
Mean relative abundance (metacommunity)
All ASVs
Above
Below
Neutral
Culicoidibacter-associated ASVs
ASV1316 (above)
ASV3950 (below)

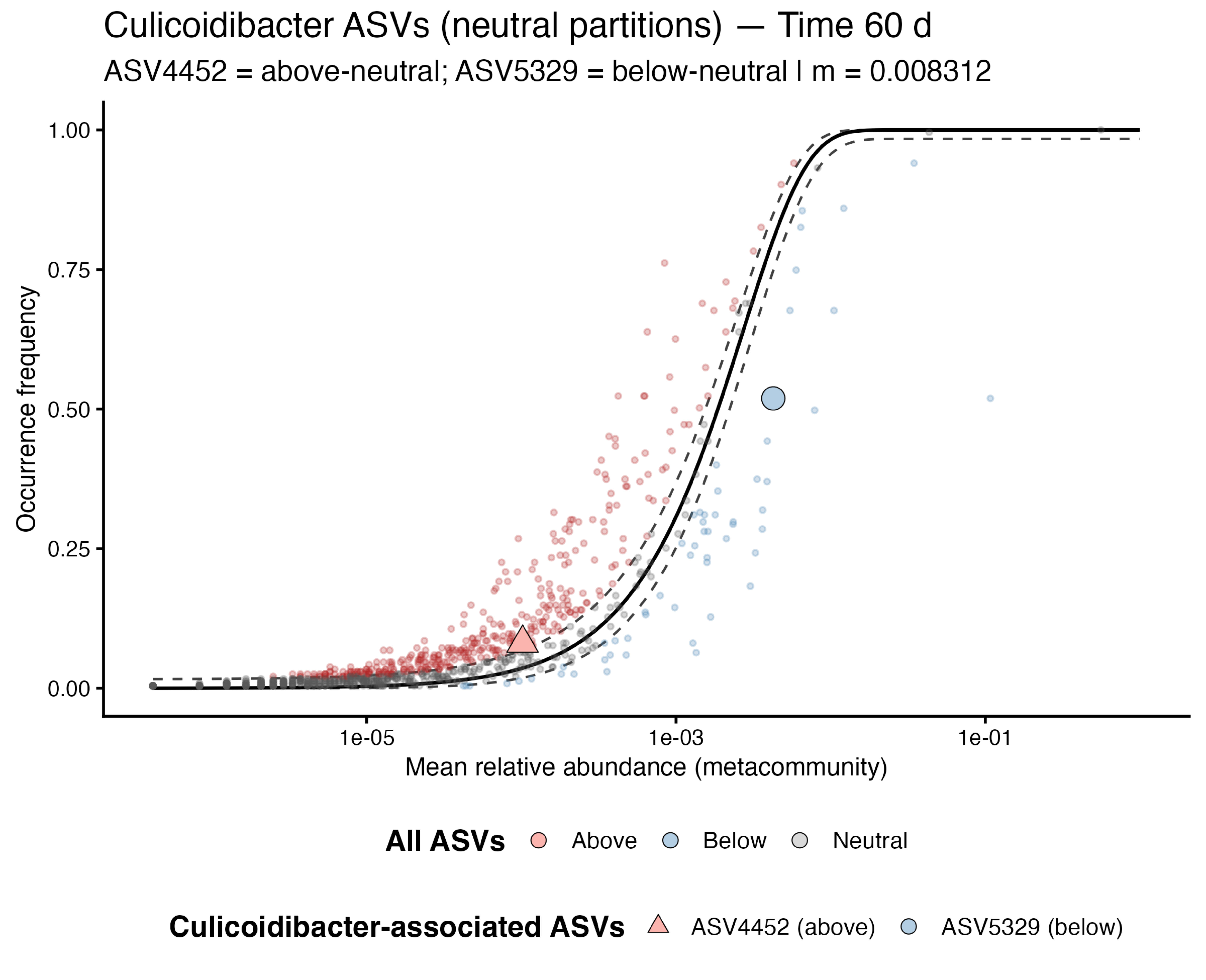

Culicoidibacter ASVs (neutral partitions) — Time 60 d
ASV4452 = above-neutral; ASV5329 = below-neutral | m = 0.008312
Occurrence frequency
1.00
0.75
0.50
0.25
0.00
1e-05
1e-03
1e-01
Mean relative abundance (metacommunity)
All ASVs
Above
Below
Neutral
Culicoidibacter-associated ASVs
ASV4452 (above)
ASV5329 (below)
